# WAGO-1 is a sexually dimorphic Argonaute protein required for proper germ granule structure and gametogenesis

**DOI:** 10.1101/2025.05.29.656876

**Authors:** Acadia L. DiNardo, Nicole A. Kurhanewicz, Hannah R. Wilson, Veronica Berg, Diana E. Libuda

## Abstract

Germ cell proliferation and proper genome inheritance are critical for maintaining fertility through generations. To promote proper germ cell development, small RNA pathways employ Argonaute proteins (AGOs) to modulate gene expression and protect against deleterious genomic elements while not silencing against self. Here we identify sexual dimorphisms in localization and function of protein structural features of the Argonaute WAGO-1 that affects sex-specific gene regulation during *C. elegans* germ cell development. During meiotic prophase I progression, we find that germ granule structural proteins and the PIWI AGO, PRG-1, display dynamic and distinct localization patterns between egg and sperm development which coincide with differential WAGO-1 localization and biophysical properties. Sexually dimorphic functions of specific WAGO-1 protein structural domains underpin these differences. Disruption or modification to the N-terminus intrinsically disordered region (IDR) of WAGO-1 leads to loss of PGL-1 phase separation only during spermatogenesis. Further, we find that these germ granule disruptions are likely due to prolonged association of the IDR with the RNA-binding pocket of WAGO-1. In addition, deletion of the MID and part of the PIWI domains causes male-specific sterility and disruption to WAGO-1 localization with PGL-1 during oogenesis. Finally, we demonstrate that these disruptions to WAGO-1 protein structure dynamically change the mRNA and sRNA landscape of adult males and hermaphrodites, in which the AGOs ALG-3/4 and VSRA-1 are misregulated. Together, these data suggest that WAGO-1 differentially regulates genes during oogenesis versus spermatogenesis, and that these differences in gene regulation may be due to the sex-specific configuration and biophysical properties of WAGO-1 within the germ granule.

## Introduction

Germ cell proliferation and immortality are critical for fertility and proper genome inheritance by the next generation. To generate viable egg and sperm cells, the regulation of the genome is finely tuned to express genes critical for meiotic progression and gametogenesis, while simultaneously suppressing deleterious genomic elements, such as pseudogenes, transposons, and somatic genes^1–5^. During oogenesis and spermatogenesis, the licensing and repression of genes is regulated by small RNA(sRNA) pathways. Widely conserved in eukaryotes, sRNA pathways modulate gene expression from transcription through translation both during periods of homeostasis and stress, and are essential for presevering fertility^6–9^.

Small RNA pathway function relies on the Argonaute(AGO) family of proteins. AGOs are bilobed proteins with four structural domains: the N-terminal lobe contains the intrinsically disordered(IDR) N-terminus and PAZ(PIWI-Argonaute-Zwille) domains, while the C-terminal lobe contains the MID and PIWI domains(Fig. S1)^10^. Small RNAs bind within the hydrophilic pocket formed between the two lobes, with the 3’ and 5’ ends of sRNAs binding with the PAZ and MID domains, respectively. Through binding of sRNAs, AGOs target sequences for licensing or silencing through direct base-pairing interactions with complementary target RNA or DNA sequences^11^. The PIWI domain has structural similarities to RNaseH, although the majority of AGOs lack catalytically active RNase function^12^. Mutations in Argonaute proteins and other machinery required for sRNA pathway regulation are tied to birth defects, developmental disorders, and infertility^13–19^.

Small RNA pathways employ non-coding sRNAs(18-30nt) for directed targeting of specific transcripts during germ cell development. In *Caenorhabditis elegans,* at least four distinct classes of endogenously encoded sRNAs are utilized for gene regulation: miRNAs, piRNAs, 26G- and 22G-RNAs. Pathways employing these sRNAs utilize two main mechanisms for target gene regulation: direct sRNA/target site binding interactions and a feedforward amplification of sRNAs triggered by primary sRNA interactions with AGOs^20^. The miRNA pathway utilizes this first mechanism, known as miRNA-mediated repression, which relies on specific sRNA/binding site interactions on the target transcript to trigger mRNA degradation and translation repression^20,21^. In contrast, piRNAs, 26G-RNAs, and 22G-RNAs are all utilized by small RNA pathways that amplify sRNAs when regulating target genes using primary and secondary AGOs.

piRNAs, known as 21U-sRNAs in *C. elegans,* and 26G-RNAs are upstream sRNAs that are loaded onto well-conserved primary AGOs. PRG-1, the only PIWI AGO in *C. elegans*, specifically binds to piRNAs and primarily targets transposons and pseudogenes in the germline^9,18,22,23^. ERGO-1, ALG-3, and ALG-4 all bind 26G-RNAs, a class of small interfering RNAs(siRNAs) that are primarily complementary to protein-coding genes expressed in the germline^13,17,24–26^. Following primary AGO binding, the sRNA-AGO complexes are recruited to target mRNA transcripts and transcribe a secondary class of small interfering RNAs, 22G-RNAs, with RNA-dependent RNA Polymerases(RdRPs)^27–30^. 22G-RNAs are loaded onto a worm-specific clade of AGOs, termed WAGOs, for targeted post-transcriptional and co-transcriptional gene regulation^17,28,30–36^. WAGOs bind specific 22G-RNAs based on the primary AGO utilized for mRNA targeting. For example, 22G-RNAs derived from sequences targeted by PRG-1 are primarily loaded onto the secondary AGOs from the WAGO Cluster, which includes WAGO-1 and PPW-2^17,18,31,37–40^. In contrast, 22G-RNAs derived from ALG-3/4 target sequences, which are primarily genes expressed during spermatogenesis, are loaded onto WAGO-10 and a sperm-specific CSR-1 isoform^17,24,32,34^.

In *C. elegans,* both the piRNA and 26G-RNA pathways display gamete-specific gene regulation. Most evident is the expression of the primary AGOs associated with the 26G-RNA pathway: ERGO-1, ALG-3, and ALG-4. ERGO-1 is only expressed in germlines undergoing oogenesis(adult hermaphrodites) while ALG-3 and ALG-4 are only expressed in germlines undergoing spermatogenesis(adult males and larval stage 4(L4) hermaphrodites)^13,17,24,26^. Regulation by piRNA pathways is similarly sexually dimorphic. Previous work in mammals demonstrated that proper regulation of piRNAs by PIWI-homologs is required for spermatocyte development, while being expendable for egg development^41,42^. Similarly in *C. elegans*, piRNAs and proper PIWI AGO PRG-1 function are required for spermatogenesis and male fertility^4,6,8,38^. Regulation by PRG-1 and downstream WAGOs, specifically WAGO-1 and CSR-1, ensures that meiotic genes are properly expressed during pachytene while simultaneously silencing deleterious genomic elements, such as transposons, in the male germline^6,8,32,34,37^. During oogenesis, PRG-1 and the piRNA pathway regulates transgenerational fertility by ensuring proper histone and ribosomal RNA expression^37,38,43^.

Numerous components of small RNA pathways localize to germ granules, which are liquid-like, phase separated compartments perinuclear to developing germ cells. These components include, but are not limited to: primary AGOs PRG-1, ALG-3 and ALG-4, secondary AGO WAGO-1, and RdRPs used for 22G-RNA transcription^8,13,17,22,24,30,44–46^. During germ cell development, germ granules separate into at least six distinct sub-compartments: the P-granule, the Z-granule, *Mutator* foci, SIMR foci, the E-granule, and the D-compartment^17,19,44,45,47–53^. Further, transgenerational sterility in *prg-1* mutants may be caused by disruption of P-granule structure^18^.

The P-granule is a sub-compartment identified by the presence of PGL-1, a scaffolding protein that aids in mRNA translation repression in developing germ cells^54–57^. Deletion of *pgl-1* is tied to transgenerational loss of fertility, underproliferation of germlines, and improper gene expression^55,56,58^. PGL-1 helps maintain fertility through translation suppression of mRNA via WAGO-1 and PRG-1^17,22,30,40,54,58,59^. WAGO-1 is implicated as a secondary AGO whose dysregulation causes transgenerational infertility due to erroneous mRNA silencing by the P-granule^37,40,54^. Loss of *wago-1* causes loss of 22G-RNAs against transposons and pseudogenes, thereby suggesting WAGO-1 is required for proper repression of deleterious elements^17,30^. Additionally, WAGO-1 demonstrates a sexually dimorphic targeting of transcripts within the piRNA pathway by primarily targeting genes required for proper progression through oogenesis^17,39^. WAGO-1 is also required for presevering fertility of *C. elegans* hermaphrodites. Tagging of the N-terminus of WAGO-1 causes a decrease in fertility, suggesting wild-type WAGO-1 structure is required for proper germ cell development^17^. The mechanisms leading to sex-specific usage of WAGO-1 remain largely unclear.

WAGO-1 and PGL-1 also interact with ZNFX-1, a well-conserved zinc finger helicase that marks the Z-granule^19,60^. Loss of ZNFX-1 causes transgenerational loss of fertility, as well as defects in both endogenous and exogenous targeting by sRNA pathways through interaction with secondary AGO, WAGO-4^17,19,49,61^. Both ZNFX-1 and PGL-1 are hypothesized to ensure proper gene regulation by sRNA pathways during germ cell development. During oogenesis, there is an interplay between ZNFX-1 and PGL-1 with WAGO-1 for monitoring transcript expression^17,19,30,37,54,61^. During spermatogenesis, ZNFX-1, PGL-1, and WAGO-1 co-localize within the germ granule^13,17,24,32,34,62^, but their role and sexual dimorphisms remain largely unknown.

Here we demonstrate that WAGO-1 is a sexually dimorphic Argonaute, with distinct dynamics and localization with germ granule components ZNFX-1, PGL-1, as well as with the PIWI AGO, PRG-1, during oogenesis and spermatogenesis. We find that the structural domains of WAGO-1 are sexually dimorphic for maintaining germ granule formation, fertility, and gene expression. Together, our findings indicate that small RNA pathways are sexually dimorphic in their modulation of germ cell development, and that these differences may be due to sexually dimorphic AGO localization and interactions within the germ granule.

## Results

### WAGO-1 abundance and accumulation within the germ granule is sexually dimorphic during oogenesis and spermatogenesis

The Argonaute WAGO-1 is a germ granule component in L4 hermaphrodites undergoing spermatogenesis^17^ and adult hermaphrodites undergoing oogenesis^17,30,54^. To examine the expression and localization of WAGO-1 in adult males undergoing spermatogenesis compared to adult hermaphrodites undergoing oogenesis, we raised an antibody against native WAGO-1(Fig. S2). Utilizing this antibody, we performed fixed immunofluorescence microscopy to examine WAGO-1 localization and morphology throughout germlines undergoing either oogenesis or spermatogenesis. We found WAGO-1 forms foci in the cytoplasm around the periphery of developing germ cell nuclei starting in the pre-meiotic tip(PMT) and persisting through late pachytene(LP) during oogenesis(Fig. 1A, top) and spermatogenesis(Fig. 1A, bottom). While WAGO-1 foci are present during both processes, males undergoing spermatogenesis have 36% of WAGO-1 protein levels compared to hermaphrodites undergoing oogenesis(Fig. 1B, C). WAGO-1 transcripts are similarly downregulated in adult males compared to adult hermaphrodites(Fig. S3). Despite lower WAGO-1 protein levels, relatively more WAGO-1 foci accumulate during spermatogenesis through mid-pachytene(MP) compared to oogenesis(Fig. 1D). This sexual dimorphism could be attributed to the higher proportion of WAGO-1 within foci instead of diffuse throughout the cytoplasm during spermatogenesis(Fig. S4). Taken together, our data suggest that while WAGO-1 transcript and protein levels are higher during oogenesis, WAGO-1 foci accumulate faster during early spermatogenesis.

**Figure 1:**
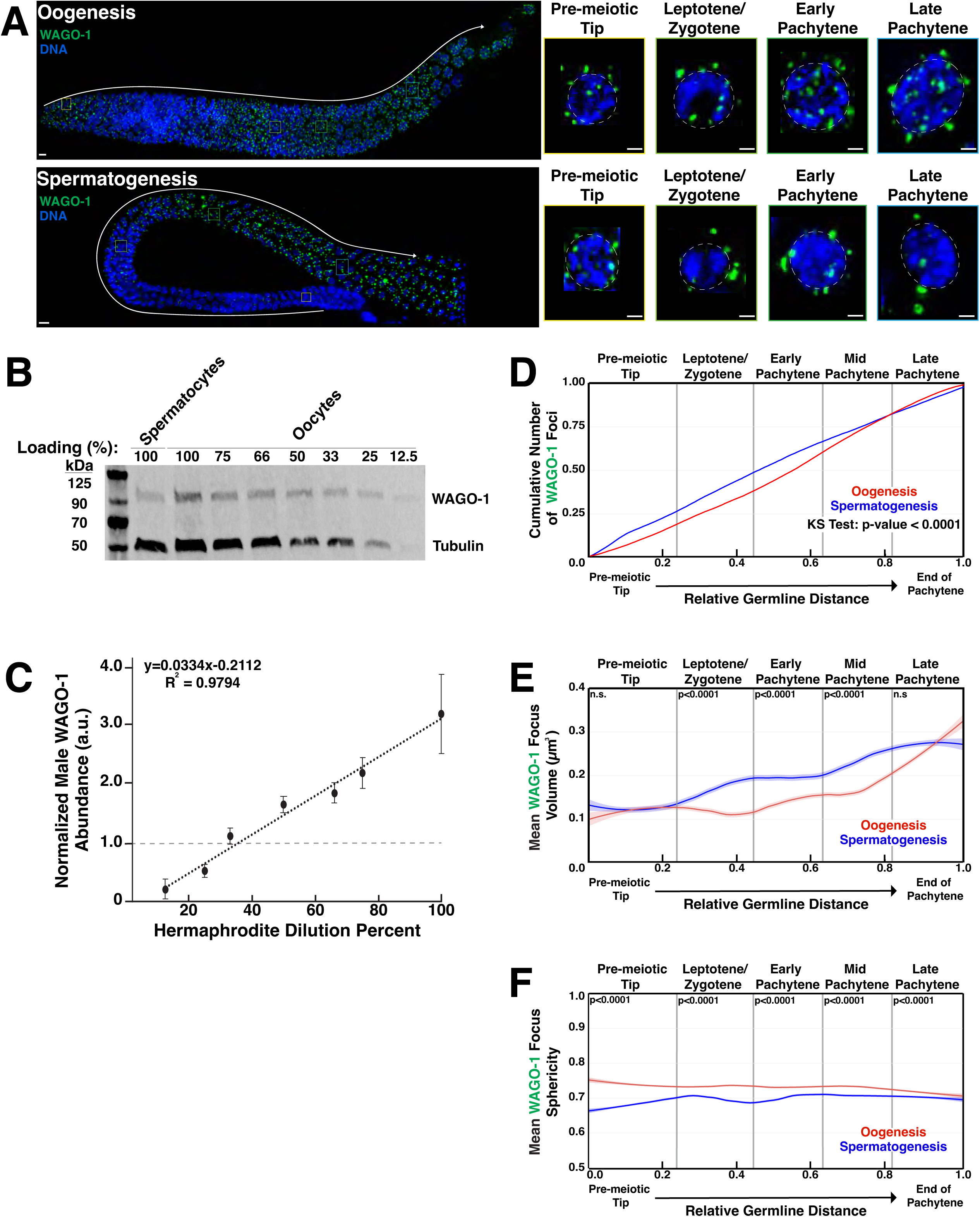
WAGO-1 localization and protein levels are sexually dimorphic within the germline. **(A)**Representative immunofluorescence images of WAGO-1(green) throughout meiosis of wild-type adult hermaphrodites(oogenesis) and adult males(spermatogenesis). Enlarged panels display individual nuclei from four distinct stages. Scale bars represent 1μm in insert panels and 10μm in full gonad images. **(B)**Dilution series Western blot with adult hermaphrodites to determine the dilution level necessary for hermaphrodite(oogenesis) WAGO-1 amounts to equal male(spermatogenesis) WAGO-1 amounts. The 100% loading is the undiluted sample for both hermaphrodites and males. **(C)**Quantification of WAGO-1 band normalized to the averaged undiluted male WAGO-1 amounts. **(D)**Empirical cumulative distribution function (ECDF) plot of WAGO-1 foci per germline through oogenesis(red) and spermatogenesis(blue) as a function of meiotic progression. **(E)**Quantification of mean WAGO-1 foci volume through oogenesis(red) and spermatogenesis(blue). **(F)**Quantification of mean WAGO-1 foci sphericity through oogenesis(red) and spermatogenesis(blue). p-values calculated using Mann-Whitney *U* tests.

### WAGO-1 displays sexual dimorphic biophysical properties during meiotic prophase I

The formation of phase separated condensates is often driven by interactions between multivalent proteins with intrinsically disordered regions(IDRs) and RNAs, and is fine-tuned by protein concentrations within the cytoplasm^63–68^. The volume of condensates can act as a read out on the overall concentration of phase-separated protein, with larger volume associated with higher protein concentrations within the cytoplasm^69–71^. To understand dynamics of WAGO-1 concentration and its ability to phase separate, we calculated the volume and sphericity of each WAGO-1 focus throughout germlines undergoing either oogenesis(Fig. 1A, top) or spermatogenesis(Fig. 1A, bottom). We found that WAGO-1 foci volume is different between sexes and depends on the meiotic stage(Fig. 1E; Table S1). In the PMT, WAGO-1 foci were larger during spermatogenesis compared to oogenesis, primarily driven by differences very early in the region(Fig. 1E, PMT oogenesis=0.124μm^2^, spermatogenesis=0.126μm^2^; Table S1). Upon entering meiosis I, as marked by the leptotene/zygotene(L/Z) zone, WAGO-1 foci rapidly gained more volume during spermatogenesis compared to oogenesis(Fig. 1E, oogenesis=0.117μm^2^, spermatogenesis=0.172 μm^2^; Table S1), and through MP remained relatively larger during spermatogenesis(Fig. 1E, EP oogenesis=0.143μm^2^, spermatogenesis=0.194μm^2^; MP oogenesis=0.168μm^2^, spermatogenesis=0.237μm^2^; Table S1). By LP, WAGO-1 foci volume plateaus during spermatogenesis while continuing to grow during oogenesis(Fig. 1E, LP). Overall, spermatogenetic WAGO-1 foci are larger during LP, despite oogenesis having larger WAGO-1 foci in the final third of the region(Fig. 1E, LP oogenesis=0.261μm^2^, spermatogenesis=0.274μm^2^; Table S1). These results suggest that WAGO-1 foci size is dynamic and sexually dimorphic through oogenesis and spermatogenesis.

To assess sexual dimorphisms in the liquid-like properties of these condensates during germ cell development, we interrogated the sphericity of WAGO-1 foci. Previous work on liquid-liquid phase separation(LLPS) of proteins correlated sphericity of foci formed with liquid-like nature of the protein and associated RNA^63,66,70,72,73^. Less spherical foci tend to contain proteins or RNAs that are more gel-like in nature, leading to less protein turnover with the cytoplasm compared to more spherical, and therefore liquid-like, foci^66,70,73–76^. Across all meiotic prophase I, our quantitative image analysis found that WAGO-1 foci are significantly more spherical during oogenesis compared to spermatogenesis(Fig. 1F; Table S3). Further, WAGO-1 foci in both sexes display opposite trends in sphericity during meiotic progression, with foci growing less spherical through oogenesis and more spherical through spermatogenesis(Fig. 1F; Table S3). Our data suggests that WAGO-1 foci are biophysically distinct between oogenesis and spermatogenesis and rapidly responds to meiotic progression.

### Sexually dimorphic colocalization of WAGO-1 with structural germ granule components ZNFX-1 and PGL-1

Prior studies in L4 hermaphrodites (spermatogenesis) and adult hermaphrodites (oogenesis) identified WAGO-1 as a germ granule component that co-localizes and directly interacts with structural germ granule component PGL-1, a marker of the P-granule^17,30,54^. WAGO-1 also interacts and modestly overlaps with the Z-granule component ZNFX-1 during oogenesis^17,60^. To determine if WAGO-1 localizes with PGL-1 and ZNFX-1 in adult males undergoing spermatogenesis, we analyzed the percent of WAGO-1 foci volume that overlaps with PGL-1 and ZNFX-1 during meiotic prophase I progression. During spermatogenesis and oogenesis, the majority of WAGO-1 foci overlaps with PGL-1(Fig. 2A-B). Furthermore, the percent of WAGO-1 foci that overlapped with PGL-1 increased from L/Z to LP, suggesting that WAGO-1 localization within the germ granule is dynamic through meiosis. Despite similar localization patterns with PGL-1 in both sexes, WAGO-1 localized significantly more with PGL-1 during spermatogenesis compared to oogenesis in L/Z and LP(Fig. 2B; Fig. S5A). WAGO-1 foci were more likely to have greater than 25% of their volume overlapping with PGL-1 during all stages of spermatogenesis compared to oogenesis(Fig. 2B L/Z oogenesis=21.9%, spermatogenesis=41.4% ; EP oogenesis=26.9%, spermatogenesis=39.0%; LP oogenesis=31.2%, spermatogenesis=50.7%; Table S5). Most WAGO-1 foci also co-localized with ZNFX-1 during oogenesis and spermatogenesis(Fig. 2C, D). Unlike with PGL-1, distinct patterns of WAGO-1 overlap with ZNFX-1 through meiotic progression depend on sex. During L/Z, significantly more WAGO-1 foci localized with ZNFX-1 during spermatogenesis compared to oogenesis(Fig. 2D). Oogenesis saw a significant increase in WAGO-1 foci overlapping with ZNFX-1 from L/Z to EP, leading to significantly more overlap during oogenesis compared to spermatogenesis(Fig. 2D, oogenesis L/Z vs EP p=0.004; EP spermatogenesis vs oogenesis p<0.0001). By LP, the percent of WAGO-1 foci overlapping with ZNFX-1 during spermatogenesis were similar to oogenesis(Fig. 2D). Together, these data suggest that oogenesis and spermatogenesis display similar trends of WAGO-1 overlap with other germ granule components. Most WAGO-1 foci have some volume overlapping with ZNFX-1 and/or PGL-1 foci throughout meiosis, with the percent of WAGO-1 overlap increasing through meiotic progression. Despite this overlap, the percent of WAGO-1 foci having any overlap with PGL-1 and ZNFX-1 was sexually dimorphic.

**Figure 2:**
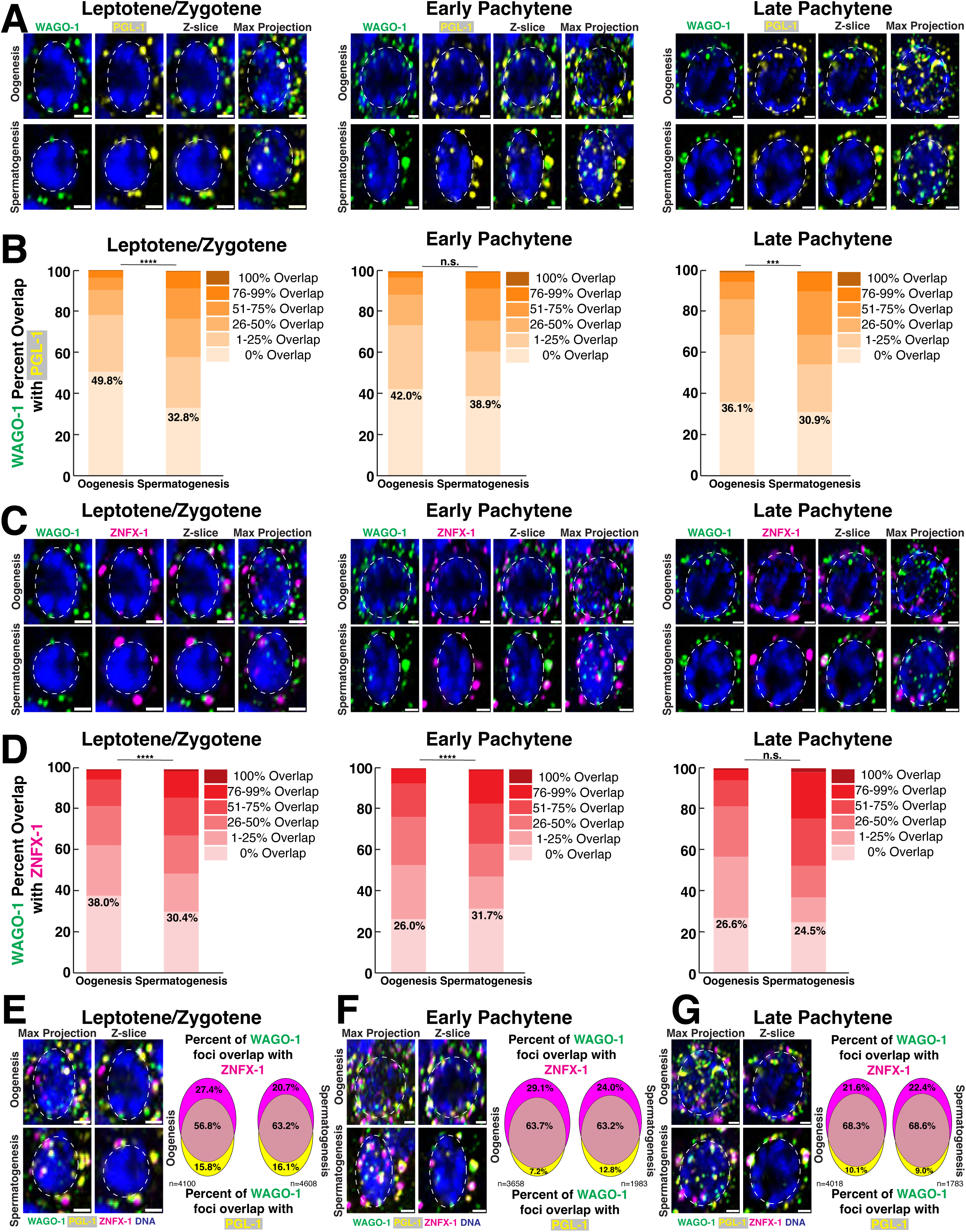
WAGO-1 co-localization with PGL-1 and ZNFX-1 is sexually dimorphic and dynamic through meiotic progression. **(A,C)**Representative images of **(A)**WAGO-1(green) and PGL-1(yellow) or **(C)**WAGO-1(green) and ZNFX-1(magenta) undergoing oogenesis(top) or spermatogenesis(bottom) during L/Z, EP, and LP. Scale bars represent 1μm. **(B,D)**Bar graphs indicating percentage of WAGO-1 foci volume overlapping with **(B)**PGL-1 foci and **(D)**ZNFX-1 foci between oogenesis and spermatogenesis from meiotic regions L/Z, EP, and LP. The numbers reported indicate the percentage of WAGO-1 foci that have 0% volume overlap with p-values of Chi-square test with Bonferroni correction. **(E, F, G)**Representative images of WAGO-1(green), PGL-1(yellow), and ZNFX-1(magenta) localization around single nuclei undergoing oogenesis(top) and spermatogenesis(bottom) during **(E)**L/Z, **(F)**EP, and **(G)**LP. Venn Diagrams show percent WAGO-1 foci overlapping with ZNFX-1 foci(top), PGL-1 foci(bottom), and with both PGL-1 and ZNFX-1 foci(middle).

### Dynamic co-localization of WAGO-1 with PGL-1 and ZNFX-1 during meiotic progression

PGL-1 and ZNFX-1 preserve the transgenerational integrity of the germline and ensure proper silencing by AGOs^19,56,60,77^. PGL-1 and ZNFX-1 also co-localize with distinct Argonautes and these localization patterns inform AGO function within sRNA pathways^17,19,54,60^. To understand WAGO-1 localization with PGL-1 and ZNFX-1, we compared the percent of WAGO-1 foci that overlapped with: only ZNFX-1, only PGL-1, or with both proteins. Throughout oogenesis and spermatogenesis, WAGO-1 was most likely to co-localize with both ZNFX-1 and PGL-1, rather than localize with a single structural component(Fig. 2E-G). When WAGO-1 localized with only a single component, it primarily co-localized with ZNFX-1 independent of meiotic stage and sex(Fig. 2E-G). These data suggest that WAGO-1 localization is not restricted to only the P-granule as marked by PGL-1. Rather, WAGO-1 displays versatile localization patterns within the germ granule, which may broaden possible WAGO-1 protein interactions.

### WAGO-1 and the PIWI Argonaute PRG-1 form dynamic toroidal rings only during spermatogenesis

Previous work established WAGO-1 as a secondary Argonaute that acts within multiple sRNA pathways including the piRNA pathway, which silences transposons and pseudogenes, in developing germ cells^17,23,30,37,38,40^. To determine how WAGO-1 and PRG-1, the primary AGO of the piRNA pathway, co-localize during germ cell development, we stained for WAGO-1, PRG-1, and ZNFX-1. During spermatogenesis, we found a subset of PRG-1 foci form toroidal rings which become more prevalent and defined in later meiosis I(Fig. 3A-C, white arrows; Fig S6). Ring-like PRG-1 structures first appear during the L/Z stage of spermatogenesis(Fig. 3A, white arrows). These rings was verified by line scans, where PRG-1 fluorescent intensity has two distinct peaks on either side of a local minima across an individual germ granule(Fig. 3B, black arrows; Fig. S7). In germ granules that contained ring-like PRG-1, we found that both WAGO-1 and ZNFX-1 foci were present but only contained a single intensity maximum between peaks in PGL-1 intensity(Fig. 3B, bottom). While we found very few ring-like structures of PRG-1 in oocytes during L/Z(Fig. S6A), numerous germ granules contained PRG-1, WAGO-1, and ZNFX-1, all with single maxima(Fig. 3B, top). In LP, we noted a germ granule containing at least one ring-like PRG-1 foci on nearly every developing spermatocyte(Fig. S6B). These PRG-1 rings were more defined compared to those observed in L/Z(Fig. 3C, bottom), with two sharp peaks of intensity around a local minimum(Fig. 3D, bottom, black arrows). WAGO-1 formed a half-moon structure with these PRG-1 rings, where line scans across the WAGO-1 foci found a single maximum(Fig. 3D, green arrow) and line scans perpendicular to the granule had two maxima around a local minimum that coincided with PRG-1 peaks(Fig. S7). In oocytes during LP, a subset of PRG-1 foci form ring structures, albeit less defined than in spermatocytes, primarily towards the end of the region(Fig. 3C-D, black arrows). These rings were independent of pocket-like germ granule structures that were also noted on developing oocytes, similar to previous observations^51^. In the condensation zone of spermatogenesis, PRG-1 rings convert into a “shell-like” casing structure that encompasses both PGL-1 and ZNFX-1 foci(Fig. 3E). Line scans reveal that PGL-1 specifically displays a local maximum where PRG-1 experiences a local minimum in intensity(Fig. 3F, top, green arrow). Unlike the rings formed by PRG-1 during earlier stages of meiotic prophase I(Fig. S8C-D), line scans across maximum projections of condensation zone rings lack local PRG-1 minimums suggesting PRG-1 forms a shell rather than a toroidal ring(Fig. 3F bottom: max projection). Overall, our data suggests a sexually dimorphic organization of a subset of PRG-1 foci, which can dynamically change from a toroidal ring-like structure localizing around WAGO-1 and ZNFX-1 into a shell-like complex that surrounds ZNFX-1 and PGL-1 only during spermatogenesis.

**Figure 3:**
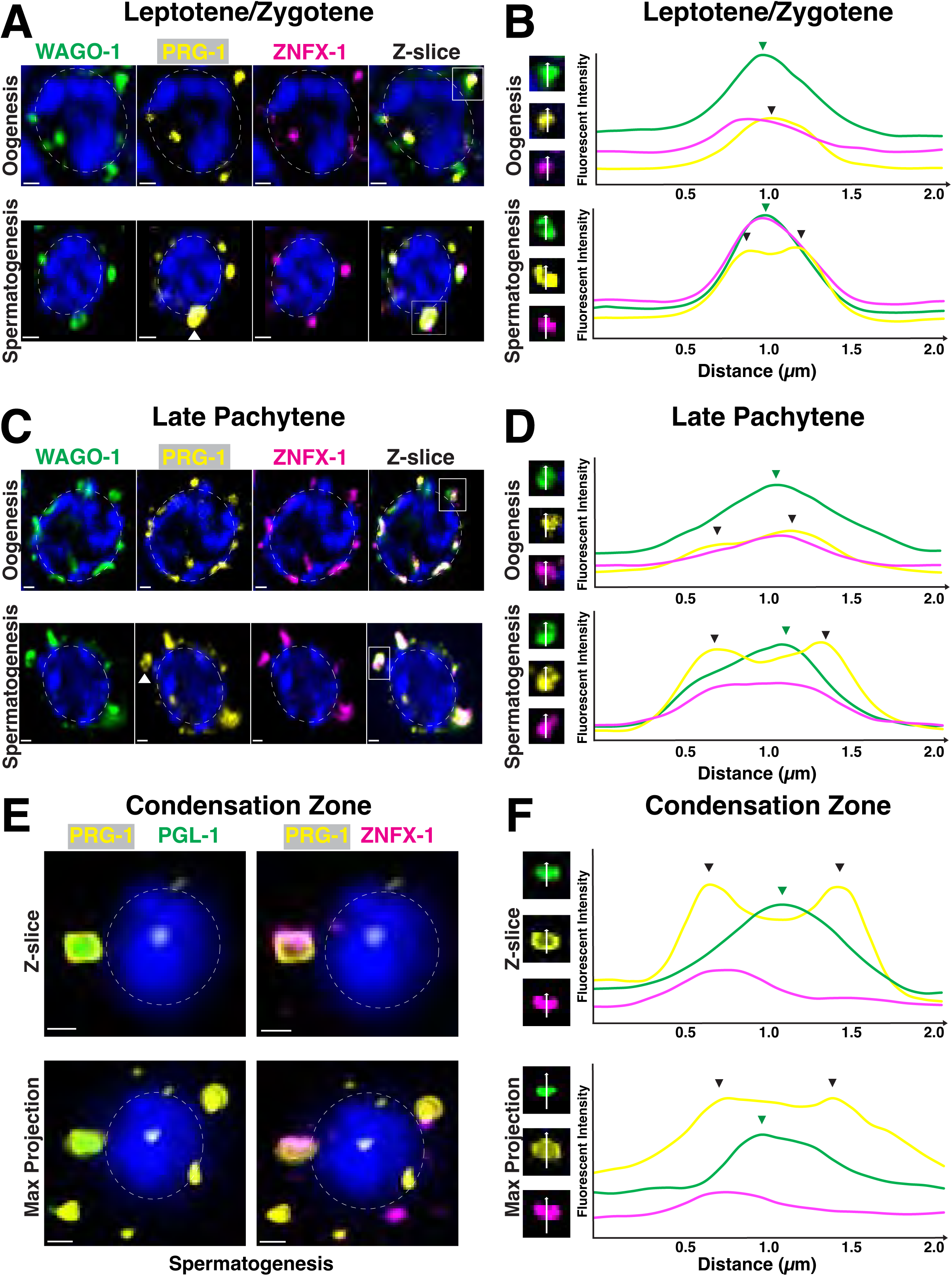
PRG-1 and WAGO-1 forms toroidal structure around ZNFX-1 during spermatogenesis. **(A,C)**Representative Z-slice of single nuclei undergoing oogenesis(top) and spermatogenesis(bottom) during **(A)**L/Z and **(C)**LP stained for WAGO-1(green), PRG-1(yellow), and ZNFX-1(magenta). Scale bars represent 1μm. **(B, D)**Left: Inset of single channels from representative germ granules boxed in **(B)**A and **(D)**C. Right: Averaged line scans of pixel intensity across germ granules during **(B)**L/Z and **(D)**LP. **(E)**Representative Z-slice(top) and max projection(bottom) of single condensing spermatid stained for PRG-1(yellow), PGL-1(green), and ZNFX-1(magenta). Scale bars represent 1μm. **(F)**Left: Inset of single channels from representative germ granules with toroidal PRG-1 structure. Right: Averaged line scans of pixel intensity across germ granules with toroidal PRG-1 structure in single Z-slice(top) and max projection(bottom). Line scan direction(white arrows). Local maxima of PRG-1 intensity(black arrows). Local maxima of WAGO-1 intensity(green arrows).

### Sexually dimorphic roles of the WAGO-1 N-terminus in PGL-1 phase separation

Previous work looking at WAGO-1 localization utilized N-terminus-tagging of both endogenous and transgenic copies of WAGO-1 with 3xFLAG or GFP^17,30,78^. Predictive structures of an N-terminal 3xFLAG tag on WAGO-1 by AlphaFold3^79^ suggest these tags may disrupt the intrinsically disordered region(IDR) of WAGO-1, but otherwise does not change the overall protein structure(Fig. 4A). To assess germ granule structure in tagged WAGO-1 strains, we stained for PGL-1. In all the tagged WAGO strains assessed, 30-50% of germlines undergoing spermatogenesis lose PGL-1 phase separation(Fig. 4B; Fig. S9). Independent of the tag used, tagging of WAGO-1 on the N-terminus caused a significant decrease in the brood size of both hermaphrodites and males in fertility assays(Fig. 4C; see Methods). For all genotypes, these drops in brood size are greater in adult males compared to adult hermaphrodites(Fig. 4C; S10). We also noted that endogenous tagging of WAGO-1 with 3xFLAG tag caused a transgenerational silencing of *wago-1* transcripts and WAGO-1 protein expression(Fig. S11). We hypothesized that these mutant phenotypes were due to the modified IDRs blocking sRNA and/or mRNA access to the RNA binding pocket of WAGO-1. WAGO-1 binds sRNAs and target mRNAs within its positively-charged binding pocket(Fig. S12)^10,11,17,30^. FLAG tags are primarily comprised of negatively-charged aspartic acid, and when added onto the IDR of WAGO-1, could allow the flexible IDR to dock within the RNA binding pocket and block any mRNA or sRNA interactions. To test this hypothesis, we utilized Molecular Dynamic(MD) simulations to assess accessibility of the RNA binding pocket with and without the 3xFLAG tag added onto the N-terminus of WAGO-1. We found that 3xFLAG::WAGO-1 had lower solvent accessible surface area (SASA) compared to wild-type WAGO-1, suggesting that when the IDR of WAGO-1 contains a 3xFLAG-tag, the binding pocket is less available to interface with target sRNAs or mRNAs(Fig. 4D; S13). By comparing the structures across multiple frames of the MD simulation, we observed the 3xFLAG::IDR within the binding pocket more often compared to the wild-type IDR(Fig. 4D; S14; S15). Collectively, our data suggests proper N-terminus structure of WAGO-1 is required for maintaining fertility and WAGO-1 expression in both hermaphrodites and males, as well as PGL-1 phase separation during spermatogenesis.

**Figure 4:**
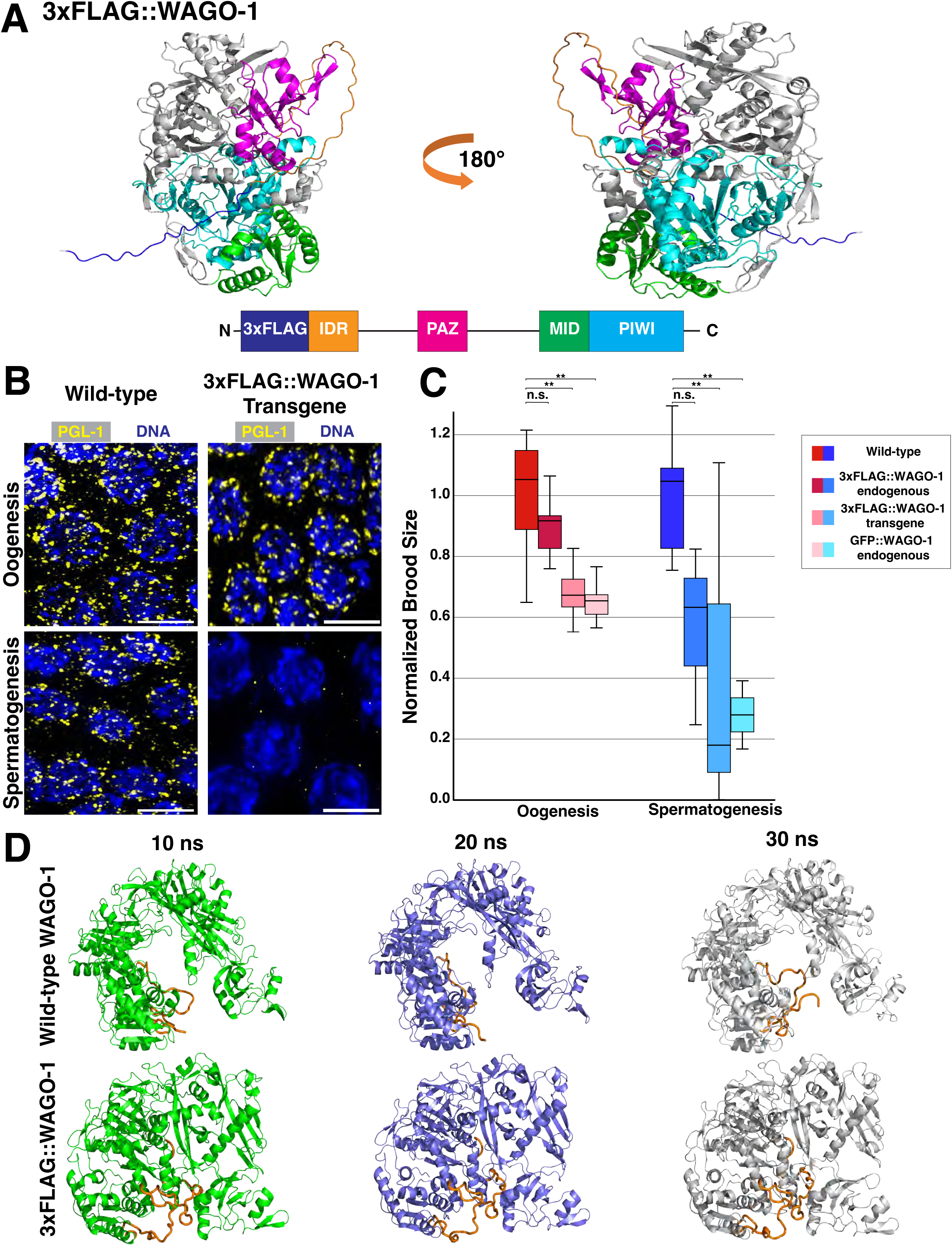
N-terminus tags of WAGO-1 cause sex-specific infertility and loss of PGL-1 phase separation. **(A)**Ribbon diagram of endogenous 3xFLAG::WAGO-1 AlphaFold3 structure(top) and gene structure(bottom). 3xFLAG tag(dark blue); N-terminus IDR(orange); PAZ domain(pink); MID domain(green); PIWI domain(light blue). **(B)**Representative immunofluorescence images of germ cells during LP undergoing oogenesis(top) and spermatogenesis(bottom) from wild-type(left) and a 3xFLAG::WAGO-1 transgene(right) strain stained for PGL-1(yellow). Scale bars represent 5μm. **(C)**Normalized brood size of adult hermaphrodites undergoing oogenesis(red; left) and adult males undergoing spermatogenesis(blue; right) of wild-type, endogenously tagged 3xFLAG::WAGO-1, 3xFLAG::WAGO-1 transgene, and endogenously tagged GFP::WAGO-1. *p<0.05, **p<0.001, Two-way ANOVA with Tukey’s multiple comparison test. **(D)**Stills from MD simulation of wild-type WAGO-1(top) and 3xFLAG::WAGO-1(bottom) taken 10 nanoseconds(ns) apart. IDR and 3xFLAG tag(orange), remaining protein in green(10ns), blue(20ns), and white(30ns).

### C-terminal truncation of WAGO-1 leads to complete infertility in males

WAGO-1 structure has a C-terminal lobe containing MID and PIWI domains(Fig. S1)^10^. Loss of the MID domain and truncation of the PIWI domain in the *wago-1(tm1414)* allele revealed that both domains are required for proper WAGO-1 function(Fig. 5A)^17,30^. To determine whether loss of these domains affects fertility, we compared the brood sizes of the *wago-1(tm1414)* allele and wild-type in both sexes(Fig. 5B). We found that, similar to tagging of the N-terminus of WAGO-1, truncation of the C-terminus causes a significant decrease in brood size for both adult hermaphrodites and males(Fig. 5B). *wago-1(tm1414)* males failed to produce any living progeny or dead eggs when mated with obligate females, suggesting that the C-terminus of WAGO-1 is required for fertility during spermatogenesis. Our analysis of the adult hermaphrodite germlines of the *wago-1* truncation mutants found reduced levels of germ cells in comparison to age-matched wild-type animals(Fig. S16). Despite decreased germ cells in adult hermaphrodite germlines, developing oocytes still properly progressed through all stages of prophase I(Fig. S16A). We next looked at localization of WAGO-1 and PGL-1 within *wago-1(tm1414)* adult hermaphrodites. Since the pachytene region of oogenesis was reduced in *wago-1(tm1414)* compared to wild-type, we analyzed all of pachytene as a singular region(Fig. 5C-D). We found that during both leptotene/zygotene and pachytene, WAGO-1 and PGL-1 are still recruited to the nuclear periphery of *wago-1(tm1414)* developing oocytes, but they rarely co-localize compared to wild-type germlines(Fig. 5C-F). These data suggest a sexually dimorphic role for the MID and PIWI domains of WAGO-1 in modulating fertility and germ granule structure. Proper MID and PIWI structure are required for male fertility, while loss of these WAGO-1 protein domains during oogenesis still allows some progeny to develop. Combined with the disruption of the germ granule observed with tagging of WAGO-1 IDR and the coinciding decreases in fertility, these data suggest that proper germ granule structure may be critical for preserving male and hermaphrodite fertility.

**Figure 5:**
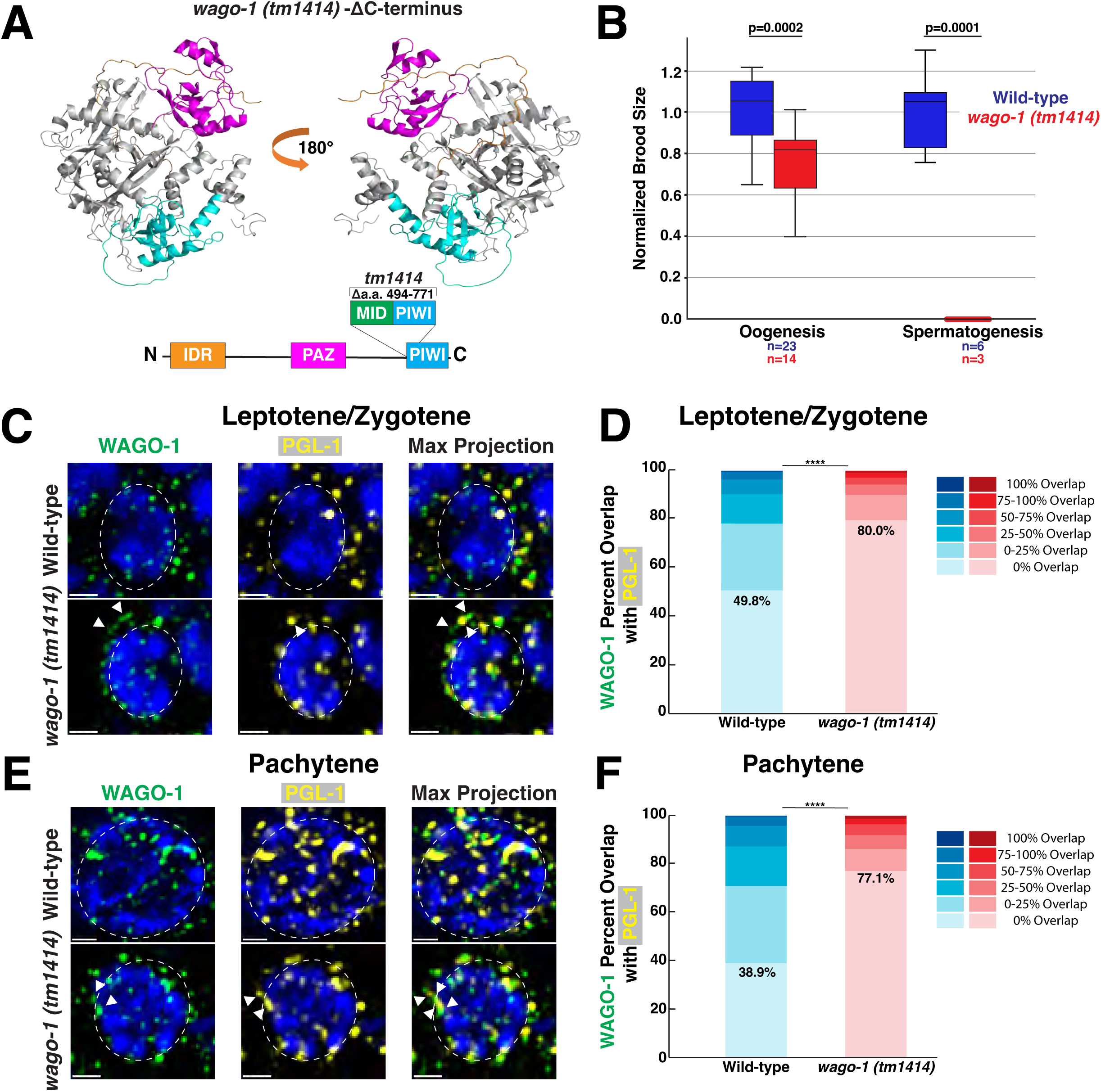
MID deletion and PIWI truncation of WAGO-1 causes male-specific infertility and WAGO-1 localization with PGL-1. **(A)**Ribbon diagram of *wago-1(tm1414)* allele AlphaFold3 structure(top) and gene structure(bottom). N-terminus IDR(orange); PAZ domain(pink); MID domain(green); PIWI domain(light blue). **(B)**Normalized brood size of adult hermaphrodites(left) and adult males(right) of wild-type and *wago-1(tm1414)*. p-values calculated using one-Way ANOVA with Bonferroni correction. **(C, E)**Representative immunofluorescence images of single nuclei undergoing oogenesis from wild-type (top) and *wago-1(tm1414)* during **(C)**L/Z and **(E)**pachytene stained for WAGO-1(green) and PGL-1(yellow). Scale bars represent 1μm. **(D, F)**Bar graphs indicating percentage of WAGO-1 foci volume overlap with PGL-1 foci from wild-type and *wago-1(tm1414)* during **(D)**L/Z and **(F)**pachytene. **** p<0.0001, Chi-square test with Bonferroni correction post-hoc.

### N-terminus tagging of WAGO-1 leads to misregulation of sperm-specific AGOs during oogenesis

To determine how disruption to native N-terminus WAGO-1 structure affects the transcriptional landscape, we performed mRNA-seq in *C. elegans* with an endogenous 3xFLAG::WAGO-1 tag, a transgene of 3xFLAG::WAGO-1, or wild-type WAGO-1. We found a significant number of misregulated genes between wild-type and WAGO-1-tagged strains(Fig. 6AB; S17). Both endogenous 3xFLAG::WAGO-1 and 3xFLAG::WAGO-1 transgene hermaphrodites showed >2x depletion of *alg-3*, *alg-4*, and *wago-10* compared to wild-type(Fig. 6AB; Supplemental Data 1). ALG-3 and ALG-4 are paralogs that bind 26G-RNAs and act as the primary Argonautes for licensing and silencing of transcripts during spermatogenesis utilizing secondary AGOs WAGO-10, CSR-1a, and RDE-1^13,17,24,32,34^. *alg-3/4* mutations cause downstream sperm-specific infertility tied to improper pseudopod formation and sperm motility, with complete sterility when *csr-1a* and *wago-10* are also knocked out^13,17,24,32^. When we examined the gene ontology(GO) terms of differentially expressed genes between wild-type and WAGO-1-tagged hermaphrodites, we found that both tagged strains were depleted of transcripts associated with Major Sperm Protein(MSP) compared to wild-type, with the transgene 3xFLAG::WAGO-1 showing a larger depletion compared to the endogenous 3xFLAG::WAGO-1 population(Fig. 6CD). This depletion in MSP-associated transcripts suggests that sperm present in adult hermaphrodites lack the proteins required for proper sperm activation, possibly driving the decrease in brood size observed in the hermaphrodites(Fig. 4B).

**Figure 6:**
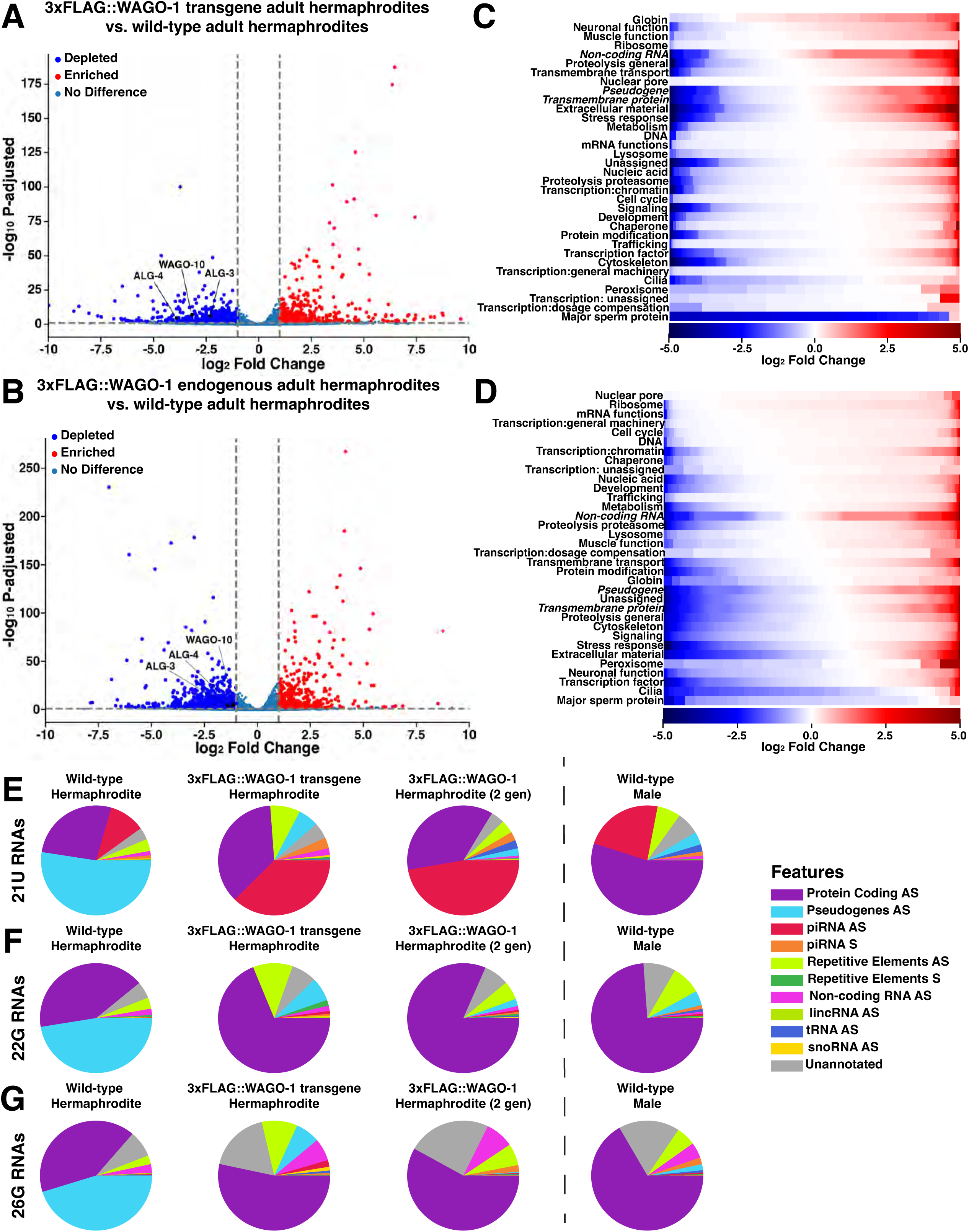
3xFLAG::WAGO-1 hermaphrodites downregulate genes required for spermatogenesis and proper sperm maturation. **(A-B)**Volcano plot of mRNA-sequencing data depicting genes up-regulated(red), downregulated(blue), or have no change(blue-grey) between wild-type and **(A)**3xFLAG::WAGO-1 transgene hermaphrodites or **(B)**endogenously tagged 3xFLAG::WAGO-1 hermaphrodites. **(C-D)**Heat map depicting individual genes, grouped by gene ontology(GO) terms, that are upregulated(red) or downregulated(blue) based on log_2_Fold Change in **(C)**3xFLAG::WAGO-1 transgene hermaphrodites or **(D)**3xFLAG::WAGO-1 endogenously tagged hermaphrodites compared to wild-type. GO groups are vertically sorted in descending order based on the proportion of genes that are upregulated. White indicates no significant change. **(E-G)**Pie charts depicting the proportion of **(E)**21U-, **(F)**22G-, and **(G)**26G-RNAs mapping to specific genetic elements in wild-type, 3xFLAG::WAGO-1 transgene, and 3xFLAG::WAGO-1 endogenously tagged hermaphrodites and wild-type males. AS=antisense, S=sense.

While we found no change in AGO expression levels between wild-type and endogenously tagged 3xFLAG::WAGO-1 males, numerous genes displayed changes in expression(Fig. S17). In both sexes, we found frequent misregulation of pseudogenes, a gene class regulated by WAGO-1(Fig. S18)^17^. Misexpression of pseudogenes, as well as genes from the other two highest classes of misregulated genes(non-coding RNAs and transmembrane proteins), was not directional(Fig. 6CD). We found that a subset of these misregulated genes significantly decreased in expression in 3xFLAG::WAGO-1 compared to wild-type, while a distinct subset saw a significant increase in gene expression in 3xFLAG::WAGO-1(Fig. 6CD). Overall, our data suggests that the addition of 3xFLAG tag on the N-terminus of WAGO-1 disrupts proper gene expression in adult hermaphrodites and males. Specifically, adult hermaphrodites with tagged WAGO-1 are depleted for sperm-specific transcripts, including *alg-3/4, wago-10,* and MSPs, which may drive the reduction in brood size. Numerous WAGO-1 targets, including pseudogenes, are misregulated in both sexes, possibly leading to observed decreases in brood size and disruption to germ granule structure.

### N-terminus tagging of WAGO-1 causes disruption to hermaphrodite and male small RNAs

Pseudogenes are primarily targeted by secondary AGOs within the WAGO Cluster of the piRNA pathway, which includes WAGO-1^17^. To determine if perturbation of WAGO-1 structure leads to dysregulation of pseudogenes due to sRNA profile changes, we performed small RNA-seq on the same samples utilized for mRNA-seq. We first looked at the global expression of different types of sRNAs based on length (18-30nt) and the 5’ nucleotide end of each sRNA(Fig. S19). In the 3xFLAG::WAGO-1 transgene adult hermaphrodites, the proportion of 22G-RNAs is depleted compared to the wild-type(Fig. S19AC). This lower share of 22G-RNAs resembles the proportions we observed in the wild-type male sRNA profile(Fig. S19D). In both adult male and adult hermaphrodite endogenously tagged 3xFLAG::WAGO-1 populations, there is a significant increase in the proportion of 26G-RNAs compared to wild-type(Fig. S19BE). Following passage of the endogenously tagged 3xFLAG::WAGO-1 strain for 100 generations, we found that the relative levels of 26G-RNAs return to baseline(Fig. S20).

Due to the misregulation of 22G- and 26G-RNAs and their role in sRNA pathways, we next wanted to look at sRNAs that associate with specific genetic elements. We also interrogated the genomic features present in the 21U-RNA populations, an sRNA class that associates with PRG-1 and templates WAGO-1 bound 22G-RNAs. We found that the overall sRNA profiles of each class are sexually dimorphic when comparing adult wild-type hermaphrodites to wild-type males(Fig. 6E). For all three classes of sRNAs in hermaphrodites, we found that the majority were antisense to pseudogenes, while in males they are primarily antisense to protein-coding genes. In the male 21U-RNA population, we see significantly larger proportion of antisense piRNAs in males compared to hermaphrodites(Fig. 6E).

We next compared the sRNA profiles of wild-type hermaphrodites to 3xFLAG::WAGO-1 hermaphrodites. In both 3xFLAG::WAGO-1 strains, the largest proportion of 21U-RNAs were antisense piRNAs, while 22G- and 26G-RNAs were primarily anti-sense to protein-coding genes(Fig. 6EG). In contrast, pseudogenes were much less prevalent in 21U-, 22G-, and 26G-RNA species for both 3xFLAG::WAGO-1 tagged hermaphrodite strains, suggesting a possible mechanism for the observed change in pseudogene expression(Fig. 6; S18). sRNAs antisense to repetitive elements, which includes transposons, were prevalent for all classes of sRNAs in the 3xFLAG::WAGO-1 transgene, suggesting that the mechanisms causing sRNA misregulation between the N-terminal tagged strains may differ(Fig. 6E-G, middle left). In endogenously tagged 3xFLAG::WAGO-1 males, we found that the proportions of sRNAs for each class were similar to endogenous 3xFLAG::WAGO-1 hermaphrodites(Fig. S21).

To determine how the transgenerational silencing of WAGO-1 changes the sRNA landscape, we compared the sRNA profiles of endogenously tagged 3xFLAG::WAGO-1 males and hermaphrodites following 100 generations of passaging(Fig. S11). Upon 100 generations of passaging, all three classes of sRNAs appeared to move towards wild-type proportions(Fig. S21). Overall, our data suggests that disruption to the IDR of WAGO-1 via a 3xFLAG tag changes the sRNA profiles of adult hermaphrodites but transgenerational silencing of endogenously tagged WAGO-1 shifts sRNA profiles back towards wild-type.

### *wago-1*(*tm1414*) hermaphrodites upregulate mRNAs enriched in adult males

Similar to N-terminus modified WAGO-1, truncation of the WAGO-1 protein with the *tm1414* allele causes male specific infertility(Fig. 5B) but preserves hermaphrodite fertility despite underproliferation of the germline and disruption to both WAGO-1 and PGL-1 localization(Fig. 5CD; S16). To determine how mRNA expression differs in *wago-1(tm1414*) compared to wild-type, we performed mRNA-seq on *wago-1(tm1414)*. We found transcript levels were perturbed in both *wago-1(tm1414)* adult males and adult hermaphrodites(Fig. 7AB). *wago-1* expression in both sexes was depleted but transcribed at significantly higher levels compared to the 3xFLAG::WAGO-1 strain after *wago-1* silencing(Fig. 7AB; S11; Supplemental Data 1; 3xFLAG::WAGO-1 endogenous hermaphrodite log_2_Fold Change=-7.18, *wago-1(tm1414*) hermaphrodite log_2_ Fold Change=-2.58; 3xFLAG::WAGO-1 endogenous male log_2_Fold Change=-6.37, *wago-1(tm1414*) male log_2_ Fold Change=-3.19).

**Figure 7:**
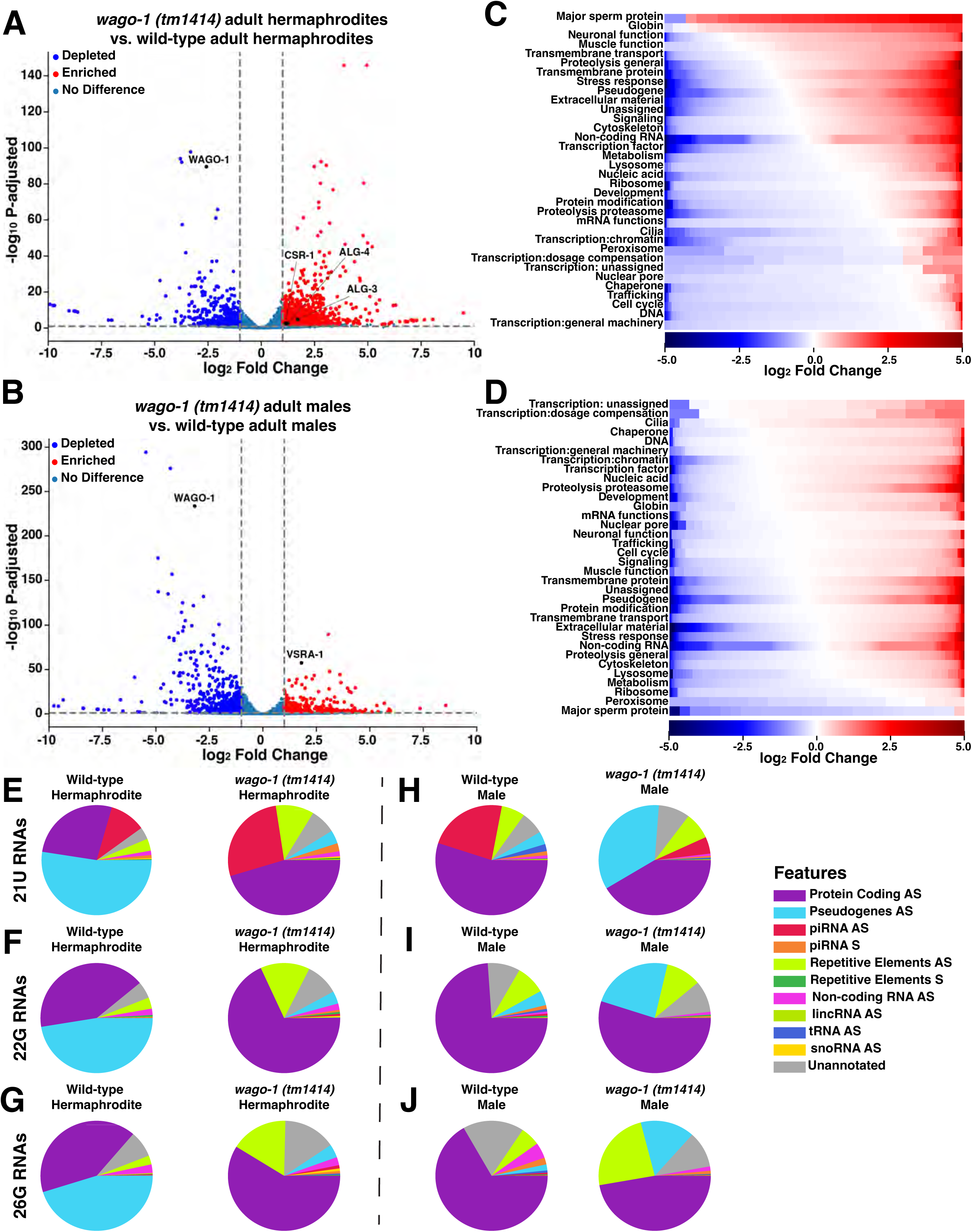
*wago-1(tm1414)* hermaphrodites and males upregulate AGOs specific to sperm and egg development. **(A-B)**Volcano plot of mRNA-sequencing data depicting genes upregulated(red), downregulated(blue), or have no change(blue-grey) between wild-type and **(A)***wago-1(tm1414)* hermaphrodites or **(B)***wago-1(tm1414)* males. **(C-D)**Heat map depicting individual genes, grouped by GO terms, that are up-regulated (red) or down-regulated (blue) based on log_2_ Fold Change in **(C)***wago-1(tm1414)* hermaphrodites or **(D)***wago-1(tm1414)* males compared to wild-type. GO groups are vertically sorted in descending order based on the proportion of genes that are upregulated. White indicates no significant change. **(E-J)**Pie charts depicting the proportion of **(E,H)**21U-, **(F, I)**22G-, and **(G, J)**26G-RNAs mapping to specific genetic elements in wild-type and *wago-1(tm1414)* **(E-G)**hermaphrodites and **(H-J)**males. AS=antisense, S=sense.

We found that *wago-1(tm1414)* hermaphrodites have increased expression of the Argonaute genes *csr-1*, *alg-3*, and *alg-4*(Fig. 7A). Paralogs ALG-3 and ALG-4, and the CSR-1a isoform are AGOs primarily expressed in germlines undergoing spermatogenesis^13,17,24,32,34^, and are upregulated in adult males compared to adult hermaphrodites(Fig. S3). All three play a role in repressing oogenesis genes and licensing the expression of genes required for proper sperm development and maturation^13,24,32,34,62^. Genes targeted for upregulation by ALG-3/4 include ones that are required for sperm mobility, such as those associated with MSP^13^. When we compared which genes classified by GO terms were up-versus down-regulated, we found that MSP genes were upregulated in *wago-1(tm1414)* hermaphrodites versus wild-type hermaphrodites(Fig. 7C).

In *wago-1(tm1414)* males, we observed increased expression of *vsra-1*, a secondary AGO primarily expressed during oogenesis that acts downstream of both ERGO-1 and CSR-1 to target protein coding genes and lincRNAs^17^. Increased expression of *vsra-1* suggests that *wago-1(tm1414)* males may repress genes required for proper sperm development despite wild-type levels of spermatogenic-specific AGOs, *alg-3/4* and *csr-1*. The majority of MSP genes were downregulated in *wago-1(tm1414)* males compared to wild-type males(Fig. 7D). Together, our data suggests that loss of the MID domain and truncation to the PIWI domain of WAGO-1 leads to a misregulation of sex-specific genes required for proper oogenesis and spermatogenesis. Specifically, adult *wago-1(tm1414)* males repress genes associated with spermatogenesis while adult *wago-1(tm1414)* hermaphrodites are enriched for spermatogenic genes.

### *wago-1(tm1414)* hermaphrodite small RNA profiles are reminiscent of wild-type males

To determine whether the small RNA landscape of *wago-1(tm1414)* is perturbed, we performed sRNA-seq on the same samples utilized for mRNA-seq. When looking at the global profile of sRNAs, we found that the proportion of 22G-RNAs compared to all 22-nucleotide sRNAs have the most striking changes between wild-type and *wago-1(tm1414)* (Fig. S22)*. wago-1(tm1414)* hermaphrodites have lower proportion of 22G-RNAs compared to wild-type hermaphrodites(Fig. S22A-B), reminiscent of the 22G-population observed in wild-type males(Fig. S22C). In contrast, the global sRNA profile of *wago-1(tm1414)* males have a higher proportion of 22G-RNAs compared to wild-type males(Fig. S22C-D).

Due to the significant changes in relative 22G-RNA levels between *wago-1(tm1414)* and wild-type, we next analyzed the proportion of sRNAs mapping to specific regions of the genome. We analyzed 22G-RNAs, as well as upstream 21U- and 26G-RNAs from which 22G-RNAs are derived. Differences in the profiles of all three sRNA classes in both adult male and adult hermaphrodite *wago-1(tm1414)* populations were identified. Many 21U-RNAs from *wago-1(tm1414)* hermaphrodites were antisense to piRNAs, a class not highly prevalent in wild-type hermaphrodites(Fig. 7E) but are in wild-type males(Fig. 7H, left). We similarly observed that *wago-1(tm1414)* hermaphrodites had 22G- and 26G-RNA profiles more similar to wild-type males compared to wild-type hermaphrodites, where the primary species are antisense to protein coding genes rather than pseudogenes(Fig. 7F-J). We also noted a significant increase of the proportion of 22G- and 26G-RNAs against repetitive elements, including transposons, in the *wago-1(tm1414)* hermaphrodites compared to wild-type(Fig. 7F-G).

We found that 21U-, 22G-, and 26G-RNAs present in *wago-1(tm1414)* males are different from wild-type males or *wago-1(tm1414)* hermaphrodites(Fig. 7H-J). The proportion of 21U-RNAs antisense to piRNAs(red) in *wago-1(tm1414)* males was significantly lower compared to wild-type males(Fig. 7H). A higher proportion of all three sRNA classes were antisense to pseudogenes compared to wild-type males(Fig. 7H-J) but still made up a lower proportion of the sRNAs than observed in wild-type hermaphrodites (Fig. 7E-G, left). Overall, our data indicates that truncation of WAGO-1 on the C-terminus leads to sRNA profiles that are more similar to wild-type males in *wago-1(tm1414)* hermaphrodites, while *wago-1(tm1414)* males display a unique sRNA expression profile suggesting sex-specific misregulation of small RNAs.

## Discussion

### Sexually dimorphic germ granule structure

Mechanisms underlying how specific sRNA pathways achieve sexually dimorphic gene regulation remains an open area of investigation. Here we find that the organization and interactions of WAGO-1 within the germ granule may play a role in sexually dimorphic gene regulation by sRNA pathways during germ cell development. There are currently several hypotheses regarding the function of germ granules. First, germ granules are hypothesized as a location for mRNA regulation by sRNA pathways due to their localization and protein composition^20,48,50,51^. mRNA is exported out of the nucleus through nuclear pore complexes(NPCs) to the cytoplasm. During germ cell development, nearly 75% of NPCs associate with a germ granule^80,81^. Actively transcribed genes required for meiosis and embryogenesis are primarily exported through NPCs directly associated with germ granules^81,82^. While germ granule localization may prime sRNA pathways by mediating direct interactions with nascent transcripts exiting the NPCs, another hypothesis indicates that this perinuclear localization may not be required. Recent works demonstrate that docking of the germ granule to the NPC is expendable for most transcript regulation if all functional sRNA and germ granule components are still properly expressed^83^. Further, the silencing of target mRNAs does not depend on localization to germ granules during embryogenesis, supporting the hypothesis that these phase-separated compartments may be incidental condensates, and gene regulation by sRNA pathways occurs in the dilute, cytoplasmic phase^84,85^. We observed that WAGO-1 expression is higher in the dilute phase compared to germ granules, suggesting that indeed WAGO-1 complexes, both with RNAs and other proteins, may likely form and function in the cytoplasm. Phase-separation, as well as localization of germ granules to NPCs, may aid in increasing the probability that recently transcribed mRNAs encounter sRNA machinery, including WAGO-1, for efficient gene regulation. Overall, sexually dimorphic structure of WAGO-1 within the germ granule suggests potential sex-specific roles of germ granule regulation in monitoring recently transcribed mRNAs and/or sex-specific protein and RNA interactions with WAGO-1 within the cytoplasm.

Our data suggest that the N- and C-terminus of WAGO-1 have sexually dimorphic roles during germ cell development. Proper IDR structure, possibly due to access to the RNA-binding pocket of WAGO-1, appears critical for proper spermatogenesis both in adult males and in L4 hermaphrodites. Changes to the IDR causes sex-specific changes to gene expression, possibly due to PGL-1 no longer forming functional germ granule compartments and/or functional cytoplasmic interactions with WAGO-1 during spermatogenesis. By contrast, the C-terminus of WAGO-1 likely aids in the ability of WAGO-1 to localize with PGL-1 during oogenesis. We found that these interactions are critical for proper gene regulation, with loss of this localization leading to similar increase in *alg-3/4* transcripts as seen when *pgl-1* is knocked down^62^. Males with C-terminus truncations are sterile, which is possibly due to the upregulation of *vsra-1*, an AGO that represses spermatogenetic genes during oogenesis. Future work conditionally expressing VRSA-1 or looking at *pgl-1* mutant males could reveal whether this *vrsa-1* upregulation is due to disruption or loss of WAGO-1 interaction with PGL-1.

### Function of the WAGO-1 IDR in PGL-1 phase separation

Our work uncovers roles of the IDR domain of WAGO-1 for maintaining germ granule stability specifically during spermatogenesis. Previous work found that WAGO-1 and PGL-1 recruitment and stability within the germ granule are independent of each other during oogenesis^54^. Through manipulation of the N-terminus of WAGO-1 by both: 1) extending the disordered region; and, 2) inserting negatively charged amino acids, we found that altered WAGO-1 structure affects PGL-1 localization within the germ granule. Specifically, proper IDR structure of WAGO-1 mediates PGL-1 phase separation during spermatogenesis. Notably, the loss of PGL-1 phase separation during spermatogenesis is limited to mid-pachytene through the condensation zone. This pattern of PGL-1 phase separation loss observed in tagged-WAGO-1 germlines matches the expression profile of the *C. elegans* extracellular-signal-regulated kinase ortholog, MPK-1, within spermatogenetic germlines^86^.

MPK-1, the *C. elegans* ortholog to human MAPK-1, regulates pachytene progression during oogenesis and spermatogenesis through phosphorylation of key proteins^86–88^. While MPK-1 expression is required for germ cell development in both sexes, MPK-1 levels are sexually dimorphic. During spermatogenesis, MPK-1 levels fall to nearly background levels following germ cell entry into early pachytene. During oogenesis, MPK-1 levels increase following meiotic entry and forms a peak at the onset of late-pachytene^86^. Low MPK-1 protein levels during spermatogenesis spatially aligns without our observation of spermatocyte-specific loss of PGL-1 phase separation, which may explain why PGL-1 remains within the germ granule through oogenesis. Phosphorylation is associated with regulating germ granule aggregation and dissolution^67,89–92^ and phosphorylation of PGL-1 during embryogenesis drives more of the protein into a phase-separated granule, even upon stressors that cause loss of phase-separation ^66,69,92^. Overall, the IDR of WAGO-1 could play a role in maintaining PGL-1 phosphorylation, and therefore phase separation, during germ cell development.

### Small RNA pathways, germ granules, and meiotic progression

Previous work found that multiple components of the germ granule are dynamic during germ cell progression, including the Mutator foci protein MUT-16, PGL-1, ZNFX-1, and PRG-1^8,51,52^. Our work builds upon these findings with WAGO-1 displaying distinct biophysical confirmations and localization patterns within the germ granule that changes through meiotic prophase I progression. WAGO-1 localization during meiotic progression suggests a dynamic role for sRNA pathways in regulating meiosis. Further, our work suggests that the role of WAGO-1 differs between the sexes.

Small RNA pathways are highly active during meiosis to: 1) repress deleterious elements that can induce genomic instabilities; and, 2) ensure proper meiosis progression^8,17,18,22^. The coordinated spatiotemporal regulation of transcripts and translation is required for the different stages of germ cell development^93,94^. Small RNA pathways provide one mechanism for modulating protein levels through post-transcriptional gene regulation. Germ granules also house recently transcribed mRNAs and translation initiation factors^81,82,95^. Specifically, two *C. elegans* isoforms of eIF4E localize to the germ granule and mediate sex-specific germ cell development^95^. In most cells, eIF4E is the limiting initiation factor, thereby altering translation rates^96^. The germ granule localization of eIF4E components could be a mechanism to modulate overall translation rates within the germline.

We find that the Argonaute WAGO-1 helps maintain proper germ granule structure throughout meiotic progression. Following IDR disruption of WAGO-1, PGL-1 foci become diffuse upon entering pachytene. These changes to germ granule structure caused by WAGO-1 perturbations may be explained by WAGO-1 dynamics throughout germline progression in wild-type strains. We found that WAGO-1 foci change in sphericity and volume during meiosis progression, which may reorganize granule structure and/or modulate gene regulation. These changes in the biophysical properties of WAGO-1 differs between the sexes(less spherical during oogenesis and more spherical during spermatogenesis) suggesting a possible change in liquid-like dynamics(Fig. 1E). Changes in the liquid-like nature of WAGO-1 could affect the protein-protein interfaces within the germ granule or the cytoplasm for proper transcriptional regulation. Future studies regarding the sexually dimorphic properties and localization patterns of other germ granule components may reveal additional mechanisms that enable sexual dimorphic gene regulation.

## Methods and Materials

### Strains

All strains were generated from the N2 background and maintained at 20°C under standard conditions of nematode growth media (NGM) with OP50 *Escherichia coli.* Strains were maintained as mating stocks unless otherwise mentioned. All spermatogenesis experiments were conducted in adult males to ensure spermatocyte-only producing populations for analysis. Mutant and tagged WAGO-1 strains were back crossed five times with N2 prior to work.

The following strains were using in this study:

N2: Bristol wildtype strain

WM616: *wago-1(ne4435[3xflag::wago-1]) I*.

WM192: *nels21 [wago-1::3xflag+ unc-119(+)] III*.

C184: *wago-1(tm1414) I*.

YY1325: *wago-4(gg620[3xflag::gfp::wago-4]) II*.

*WM205: unc-119(ed3) III; nels22[wago-1::GFP + unc-119(+)]*

YY916: *znfx-1(gg554[3xflag::gfp::znfx-1]) II*.

JMC250: *prg-1(tor149[prg1::gfp::flag]) I; znfx-1(gg634[ha::tagRFP::znfx-1]) II*.

CB4108: *fog-2(q71) V*.

### WAGO-1 Antibody Production

Peptides for the last fourteen amino acids (Cys-GYKQTDLNQKRVNA) of the C-terminus of WAGO-1 were produced by Biomatik. Antibodies against the WAGO-1 peptides were produced in chicken and affinity purified by Pocono Rabbit Farms. Antibody specificity was verified utilizing western blotting (Figure S2) and immunofluorescence.

### Immunofluorescence

Immunofluorescence was performed as described in Claycomb *et al.,* 2009 with the following adaptations. Briefly, gonads from adult male and hermaphrodite worms at 18-22 hour post-L4 stage were dissected together into 1x sperm salts (50 mM PIPES, pH 7.0, 25mM KCl, 1 mM MgSO4, 45 mM NaCl, 2 mM CaCl2) on the same VWR Superfrost Plus slides, frozen and cracked on dry ice for 10 minutes and then fixed at −20°C for 5 minutes in 100% methanol, 5 minutes in 50% methanol/50% acetone, and 5 minutes in 100% acetone. Samples were blocked in 1xPBST + 3% BSA at room temperature for 15 minutes. Primary antibody dilutions were made in 1xPBST + 3% BSA and added to the slides. Slides were covered with a parafilm cover slip and incubated overnight at 4°C in a humid chamber. Slides were washed 3 x 10 minutes in 1xPBST and then blocked in 1xPBST + 3% BSA for 15 minutes at room temperature. Secondary antibody dilutions were made at 1:200 in 1x PBST + 3% BSA using Invitrogen goat Alexa Fluro-labeled antibodies and were added to the slides. Secondary antibody incubation occurred for 1 hour at room temperature in a dark humid chamber. Slides were then washed 3×10 minutes in 1xPBS in the dark and then incubated with 2 μg/ mL DAPI for 5 minutes in a humid chamber in the dark. Slides were then washed with 1xPBS for 5 minutes in the dark and then mounted with Vectashield. Slides were sealed with nail polish immediately following mounting and then stored at 4°C prior to imaging. All slides were imaged within 2 weeks of preparation. The following primary antibody dilutions were used: WAGO-1 anti-chicken (1:1000, this study), monoclonal mouse aPGL-1 K76 (1:20, Developmental Studies Hybridoma Bank), rabbit anti-PGL-1 (1:1000, Susan Strome Lab), mouse anti-FLAG (1:1000, Millipore Sigma F1804-50UG), ChromoTek GFP-booster Alexa Fluor® 488 (1:200, ChromoTek gb2AF488), rabbit anti-HA (1:500, Abcam ab9110).

### Imaging

Immunofluorescence slides were imaged at 1024 x 1024 pixel dimensions on an Applied Precision DeltaVision microscope with a 63x lens and a 1.5x optivar. Images were acquired as Z stacks at 0.2μm intervals and deconvolved with Applied Precisions softWoRx deconvolution software.

### Foci quantification and Overlap

Foci quantification adapted from^97^. Briefly, germ granule foci were defined as Surface objects in Imaris (Bitplane) with the following settings: Smooth (not enabled), Background 0.513, and Seed Point Diameter (not enabled). Foci were analyzed if larger than 0.034 μm^2^ and smaller than 10 μm^2^ in volume. Counts, volume, and sphericity of granule structures were then aligned along an X-Y axis utilizing a Gonad Linearization Algorithm as described in Toraason *et al.,* 2021 and normalized from the pre-meiotic tip to the end of pachytene. Differences in counts were quantified using two sample Kolmogorov-Smirnov tests. Volume and sphericity were graphed using the plotnine.geom_smooth metric and the loess method. Counts were binned along meiotic progression utilizing well established nuclei morphology criteria to indicate meiotic stage: pre-meiotic tip, leptotene/zygotene, early pachytene, mid pachytene, late pachytene ^98^. Differences in distributions were calculated utilizing Mann-Whitney U test with a Bonferroni correction using SciPy stats.

To determine if foci colocalized, we applied the Surface-to-Surface Percent Overlap function in Imaris to identify and replicate surfaces with greater than 0.1% volume overlap. These overlapping surfaces were then given unique colocalization identity intensity channels. A focus was considered co-localized with another in all analyses if two or more foci of different types (*i.e.* PGL-1 and WAGO-1) with the same unique colocalization intensity value could be identified in the exported data. Protein foci were considered to have no overlap if the primary protein foci had no volume overlapping with a secondary protein focus. 50% overlap corresponds with half of a single primary foci’s volume being internal to a secondary protein’s foci, while 100% occurs when the entirety of a single primary protein focus resides within a secondary protein focus. The percent overlap of the primary foci, though, does not report the percent of the secondary foci volume associating with the primary. Co-localized foci were then aligned on an X-Y axis utilizing the Gonad Linearization Algorithm and binned based on normalized progression through the germline. Differences in the percent co-localization of foci within each bin was calculated utilizing Chi Squared Test with a Bonferroni Adjustment.

### Percent of WAGO-1 protein in germ granules

To quantify WAGO-1 protein amount in germ granules versus diffuse in the cytoplasm Surface objects were created for the entire germline in addition to the germ granule foci described above. The germline was then separated into five regions, pre-meiotic tip, leptotene/zygotene, early pachytene, mid pachytene, and late pachytene, for individual stage analysis. The total fluorescence sum intensity of the germ granules was divided by the total sum fluorescence intensity of the entire region to determine the percent of total WAGO-1 present in the germ granule. To normalize for variations between stains as well as the possibility of secondary antibodies forming granule-like structures, the ratio of total fluorescence sum intensity of granule-like structures to total sum fluorescence intensity of the entire region for samples only stained with secondary antibodies was subtracted from the calculated experimental ratios.

### Male and hermaphrodite fertility assays

For hermaphrodite fertility assays, hermaphrodites of each genotype were isolated at the L4 developmental stage and aged 18-22 hours to adult stage. Individual adult hermaphrodites were then singled on small plates (35 x 10 mm) with a small OP50 dot and allowed to lay for 24 hours. Following 24 hours, the adult hermaphrodite was moved to a fresh small plate (35 x 10 mm) with a small OP50 dot and allowed to lay for an additional 24 hours. This was repeated for a total of 5 plates. Following 24 hours on the fifth plate, the adult hermaphrodite was flamed off. 1-2 days following the removal of the adult hermaphrodite, plates were counted for number of living progeny, number of dead eggs, and number of unfertilized eggs. Plates where the hermaphrodite had run away or died were excluded. Differences in the ability of *wago-1* mutant hermaphrodites to produce viable progeny were analyzed using Two-way ANOVA with Tukey’s multiple comparison test.

For male fertility assays, males of each genotype and obligate females (CB4108: *fog2Δ*) were isolated at the L4 developmental stage and aged 18-22 hours to adult stage. Individual obligate females were then singled and paired with an individual male on small plates (35 x 10 mm) with a small OP50 dot ringed with garlic extract to keep males from leaving the plate for 24 hours. After 24 hours males were moved to a new plate with an additional 18-22 hours post-L4 stage obligate female and allowed to mate for another 24 hours. After removal of the male, obligate females were allowed to lay eggs for 24 additional hours and then scored for living progeny, dead eggs, and unfertilized eggs. Pairs with no unfertilized eggs or where either the male or obligate female died before egg counting were excluded. Differences in the ability of *wago-1* mutant males to produce viable progeny were analyzed using Two-way ANOVA with Tukey’s multiple comparison test.

### HIM-8 RNAi male production

Due to the low brood sizes produced by males following mating with hermaphrodites for tagged and *wago-1 (tm1414)* strains, to increase male populations for mRNA- and sRNA-sequencing we utilized HIM-8 RNAi. Briefly, bleached synchronized *C. elegans* were allowed to grow on NGM+AMP+IPTG plates seeded with *E. coli* expressing the pLT 653 plasmid starting from L1s. Worms were grown at 20°C on RNAi plates until 18-22 hours post-L4 before being placed onto NGM plates with OP50 and grown at 20°C. Plates were then screened daily for male progeny. During this RNAi treatment, we found that WM192 failed to produce males at higher rates than seen naturally. We therefore were unable to collect enough WM192 for downstream mRNA- and sRNA-sequencing experiments.

### Total RNA extraction and isolation for mRNA and sRNA-sequencing

300 adult hermaphrodites or 300 adult males 18-22 hours post-L4 stage were picked onto unseeded plates and washed off with 1 mL of cold M9. Worms were then spun down for 1 minute at 2000 x g and placed on ice for 3 minutes. Supernatant was then aspirated off to not disturb the worm pellet and washed with 200 μL of M9 and again pelleted. Worms were washed one final time with 200 μL of DNase and RNase free UltraPure distilled water (Invitrogen), spun for 1 minute at 2000 RPM and all but 50uL of liquid was removed. 500 μL of TRIzol (ThermoFisher) was added and tubes were flash frozen and placed in −80°C freezer for 15 minutes. Samples were then removed and vortexed for 15 minutes at room temperature with 100 μm acid washed, RNase free beads (Sigma-Aldrich, <106 μm). Freeze-thaw and vortexing was then repeated twice more for a total of 3 times. 100 μL of chloroform was then added to samples and tubes were shaken vigorously for 15 seconds and incubated at room temperature for 3 minutes. Tubes were then centrifuged at 12,000 x g for 15 minutes at 4°C. The top aqueous layer was then removed and transferred to a new tube. Phenol: chloroform: isoamyl alcohol was added 1:1 in the new tube and mixed for 15 seconds. Samples were then centrifuged at 12,000 x g for 15 minutes at 4°C. Top aqueous phase was again transferred to a new tube and 20 μg of GlycoBlue (15 mg/mL, ThermoFisher) and 1:1 ratio of isopropanol was added to new tube. Samples were then incubated at −20C overnight or −80C for 1 hour. Samples were then centrifuged at 16,000 x g for 30 minutes at 4°C and supernatant then removed. The pellet was then with washed with 900 μL of 70% ice-cold ethanol for 10 minutes and then centrifuged at 16,000 x g for 10 minutes at 4°C. Supernatant was again removed and washing and centrifugation was repeated. Following last centrifugation, as much ethanol as possible was removed and pelleted was left to airdry for 10 minutes at room temperature. Pellet was then resuspended in 12 μL of RNAse-free water and purity and amount was quantified using Nanodrop. Samples were then diluted to 1 μg of RNA per 50 μL and used for downstream preparations.

### Western Blotting

To generate protein lysates, 300 adult hermaphrodites or 300 adult males 18-22 hours post-L4 stage were picked onto unseeded plates and washed off with 1 mL of cold M9. Worms were then pelleted at 2000 x g for 1 minute and incubated on ice for 3 minutes. Supernatant was removed and worms were washed in 200 μL of M9 and again pelleted twice. Following the second pelleting step, all but 35 μL was removed, and worms were then mixed with NuPAGE LDS sample buffer (4x, Invitrogen) and DTT for a final concentration of 1xLDS and 100 mM DTT and boiled for 10 minutes at 95°C with intermittent vortexing. Samples were then run on SDS-PAGE gel (BioRad) at 100 V for 1:15 in 1x Laemelli Running Buffer. Samples were transferred at 100 V for 2 hours onto 0.45 μM nitrocellulose membrane (Thermo) in 1x Transfer Buffer (3.027 g Tris, 14.4 g Glycine, 200 mL MeOH). Membrane was then blocked for 1 hour at room temp in 5% milk powder + 1x PBST. Primary antibodies were then diluted in the milk powder solution and left at 4°C overnight with agitation. Membrane was then washed 3 x 5 minutes in 1x PBST and then incubated with Licor secondary antibodies in 1x PBST for 1 hour under agitation in the dark. Membrane was then washed in 1x PBST for 30 minutes in the dark before imaging using the Licor Imager. The following primary antibody dilutions were used: WAGO-1 anti-chicken (1:500, this study), alpha Tubulin anti-mouse (1:1000, Abcam ab7291).

### Structure predictions

Structures for WAGO-1, 3xFLAG::WAGO-1, and *wago-1 (tm1414)* were all created utilizing AlphaFold3^79^. Sequences for *wago-1* and *wago-1(tm1414)* were taken from WormBase and spliced and translated using aPe^99^. The sequence for 3xFLAG::WAGO-1 was derived from the WM616 strain from Dokshin *et al.* The electrostatics were predicted utilizing the APBS-PDB2PQR software suite and PyMol plugin^100^.

### Model construction for molecular dynamic simulations

For wildtype and 3xFLAG::WAGO-1 simulations, we started with the top three predicted AlphaFold3 models utilizing the sequence of WM616 from Dokshin *et al.,* 2018 with and without the 3xFLAG tag (Figure S1; Figure 4A). For all simulation parameters we utilized GROMACS 2023.4^101,102^ with the CHARMM36 2021 forcefield^103^ with TIP3P waters^104^. All simulations were done in the NPT ensemble. We ran each simulation for a minimum of 500ns. All scripts for running simulations are available on github (https://github.com/harmslab/setup_md). We did all calculations using the talapas high-performance computing cluster at the University of Oregon.

### Analysis of MD simulation trajectories

For analysis of MD simulations we visualized results using Visual Molecular Dynamics^105^, python scripts using the MDAnalysis library^106,107^, and PyMol^108^. We calculated the solvent-accessible surfaces areas using python scripts integrating the FreeSASA library^109^.

### mRNA library preparation and sequencing

Samples were prepared for sequencing using the KAPA mRNA HyperPrep Kit for Illumina Platforms (KAPA Biosystems) following the protocol provided by manufacturer. The resulting DNA library was visualized and checked for quality using the 5200 Fragment Analyzer System (Agilent). Samples were then sequenced on the NovaSeq 6000 with a coverage of 25-30 million reads per a sample.

### Small RNA library preparation and sequencing

Samples were prepared for sequencing using the Revvity small RNA kit following the protocol provided by manufacturer. The resulting DNA library was visualized and checked for quality using the 5200 Fragment Analyzer System (Agilent). Samples were then sequenced on the NovaSeq 6000 with a coverage of 10 million reads per a sample.

### mRNA-seq analysis

The mRNA sequences obtained from the sequencer were first assed for quality using MultiQC^110^. Adapter sequences were then removed using Trimmomatic for paired ends (version 0.36^111^) and run through MultiQC again to asses quality. The trimmed reads were then aligned to the *C.* elegans PRJNA13758 ce11 genome assembly (WormBase version WS280) using STAR (version 2.7.11b^112^). Reads were then counted using HTSeq (version 2.0.3^113^) and differential expression was determined utilizing DESeq2 (version 3.20^114^). Genes were considered differentially expressed if their Log_2_ Fold Change was great than 1 or less than −1 and had a p-adjusted value of less than 0.5. GO terms for each gene were assigned using WormCat 2.0^115^.

### sRNA-seq analysis

The sRNA sequences obtained from the sequencer were first assed for quality using MultiQC^110^. Adapter sequences were then removed using Trimmomatic for paired ends ends (version 0.36^111^) and run through MultiQC again to asses quality. The trimmed reads were then aligned to the *C.* elegans PRJNA13758 ce11 genome assembly (WormBase version WS280) using STAR (version 2.7.11b^112^). Reads were then counted using the custom R-script created by Seroussi *et al.,* 2023 aligned to the following genome annotations: WormBase version WS280 PRJNA13758 ce11, *C. elegans* miRNAs from miRbase (release 22.1), repats and transposons annotations from RepeatMasker+Dfam (ce10, October 2010, RepeatMasker open-4.0.6, Dfam 2.0). Counts for different gene biotypes were then utilized for total small RNA sequencing landscape. For specific enrichment of 21U, 22G, and 26G-RNAs, we removed all sense reads to non-coding RNAs, rRNAs, snoRNAs, snRNAs, tRNAs, lincRNAs, protein coding genes, and pseudogenes due to the likely event they were caused by degradation. We also removed all RNAs transcribed in high copy in *C. elegans* including miRNAs and anti-sense rRNA and snRNAs to enrich for small RNAs that would be bound by AGOs in the endogenous RNAi pathways ^116^.

## Data Availability

The mRNA-sequencing and small RNA-sequencing data sets generated in this study are available on the NCBI BioProject database (https://www.ncbi.nlm.nih.gov/bioproject/) under accession number PRJNA1252786. Strains and WAGO-1 antibodies are available upon request.

## Acknowledgments

We thank Z. Bush, J. Brown, C. Crahan, C. Cahoon, and J. Freedman for thoughtful discussion and comments on the manuscript. Special thanks to M. Harms for helping with MD simulations and SASA analysis. We also thank A. Barkan, B. Bowerman, D. Garcia, and S. Hansen for feedback on figures. We thank the Claycomb lab for PRG-1::GFP::FLAG; HA::tagRFP::ZNFX-1 strain, the Mello Lab for sharing the 3xFLAG::WAGO-1 endogenously tagged, 3xFLAG::WAGO-1 transgene, and *wago-1(tm1414)* strains, and the Strome Lab for sharing their PGL-1 α rabbit antibody. We are grateful to the University of Oregon’s Genomics and Cell Characterization Core Facility for small RNA and mRNA sample prep and sequencing. We thank the CGC for providing multiple strains for this study (funded by National Institutes of Health P40 OD010440).

## Supplemental Figure Titles and Legends

**Supplemental Figure 1: WAGO-1 protein structure**. **(A)** Cartoon schematic of WAGO-1 protein structure. Orange, IDR; pink, PAZ domain; green, MID domain; light blue, PIWI domain. **(B)** Ribbon diagram of WAGO-1 AlphaFold3 predicted structure. **(C)** Front views of WAGO-1 ribbon diagram of AlphaFold3 predicted structure.

**Supplemental Figure 2: Western blot confirmation of WAGO-1 anti-chicken antibody.** Representative western blot for WAGO-1 protein size in wild-type (N2) and endogenously tagged 3xFLAG::WAGO-1 (WM616) strains. Wild-type WAGO-1 size = 105kDa; endogenously tagged 3xFLAG::WAGO-1 size = 108kDa.

**Supplemental Figure 3: WAGO-1 mRNA upregulated in hermaphrodites compared to males.** Volcano plot of mRNA-seq between wild-type adult males and hermaphrodites. Genes upregulated in hermaphrodites are in red. Genes upregulated in males are in blue. Blue-grey dots represent genes with no statistical difference in regulation between the sexes. Black dots and labels denote differentially regulated AGOs between adult hermaphrodites and adult males.

**Supplemental Figure 4: Meiotic-stage specific effect on WAGO-1 phases during oogenesis and spermatogenesis.** Line graph depicting average WAGO-1 fluorescent intensity within germ granules as a percentage of total WAGO-1 fluorescent intensity for PMT, L/Z, EP, MP, and LP regions of the germline undergoing oogenesis (red) and spermatogenesis (blue). For each stage n=9 gonads were analyzed.

**Supplemental Figure 5: Kernal density estimate plots of percent of WAGO-1 foci volume overlap with PGL-1 and ZNFX-1. (A,B,C)** Kernal density estimate (KDE) plots of the percent of WAGO-1 foci volume overlapping with PGL-1 foci undergoing oogenesis (red) and spermatogenesis (blue) in the **(A)** L/Z, **(B)** EP, and **(C)** LP regions of meiosis. The p-values were calculated using two-sample Kolmogorov-Smirnov test. **(D,E,F)** KDE plots of the percent of WAGO-1 foci volume overlapping with ZNFX-1 foci undergoing oogenesis (red) and spermatogenesis (blue) in the **(D)** L/Z, **(E)** EP, and **(F)** LP stages of meiosis. The p-values were calculated using two-sample Kolmogorov-Smirnov test.

**Supplemental Figure 6: PRG-1 rings are most prevalent in spermatogenesis during Late Pachytene. (A)** Bar graphs indicating the percent of nuclei undergoing oogenesis (red) and spermatogenesis (blue) with 0, 1, 2, or 3 PRG-1 rings during L/Z. **(B)** Bar graphs indicating the percent of nuclei undergoing oogenesis (red) and spermatogenesis (blue) with 0, 1, 2, or 3 PRG-1 rings during LP. For each stage n=9 gonads were analyzed. For total number of nuclei see Table S7.

**Supplemental Figure 7: PRG-1 forms ring-like structure and WAGO-1 forms half-moon structure during spermatogenesis. (A)** Left: Zoom in panels of single channels from representative germ granules boxed in Fig. 3A. White arrow indicates line scan direction. Right: Averaged line scans of pixel intensity across germ granules from oogenesis (top) and spermatogenesis (bottom) during L/Z. **(B)** Left: Zoom in panels of single channels from representative germ granules boxed in Fig. 3C. White arrow indicates line scan direction. Right: Averaged line scans of pixel intensity across germ granules from oogenesis (top) and spermatogenesis (bottom) during late pachytene. Black arrows indicate local maxima of PRG-1 intensity, green arrows indicate local maxima of WAGO-1 intensity.

**Supplemental Figure 8: PRG-1 forms hollow rings during spermatogenesis. (A,C)** Representative Z-slice and max-projection of single nucleus either undergoing oogenesis (top) or spermatogenesis (bottom) during **(A)** L/Z and **(C)** LP stained for WAGO-1 (green), PRG-1 (yellow), and ZNFX-1 (magenta). Scale bars represent 1 μm. **(B,D)** Left: Zoom panels of single channels from representative germ granules boxed in **(B)** A and **(D)** C. White arrow indicates line scan direction Right: Averaged line scans of pixel intensity across max projection of germ granules from oogenesis (top) and spermatogenesis (bottom) during **(B)** L/Z and **(D)** LP. Black arrows indicate local maxima of PRG-1 intensity, green arrows indicate local maxima of WAGO-1 intensity.

**Supplemental Figure 9: Tagging WAGO-1 and WAGO-4 on N-terminus causes male specific loss of PGL-1 phase separation during late pachytene. (A)** Representative immunofluorescence images of nuclei (top) and PGL-1 (bottom) throughout spermatogenesis from an adult male with 3xFLAG::WAGO-1 transgene. Leptotene/zygotene (L/Z, white) and early pachytene (EP, light blue) zones are highlighted and identified based on nuclei morphology. Scale bars represent 10 μm. **(B,C)** Max projection images of L/Z (top) and LP (bottom) nuclei from **(B)** adult hermaphrodites undergoing oogenesis and **(C)** adult males undergoing spermatogenesis stained for PGL-1 (yellow) from the following strains (L to R): wild-type, 3xFLAG::WAGO-1 transgene, endogenously tagged 3xFLAG::WAGO-1, endogenously tagged GFP::WAGO-1, and endogenously tagged 3xFLAG::GFP::WAGO-4. Scale bars represent 5 μm.

**Supplemental Figure 10: Endogenous N-terminus tagging of WAGO-1 and WAGO-4 causes male specific decrease in brood size.** Normalized brood size of adult hermaphrodites undergoing oogenesis (red) and adult males undergoing spermatogenesis (blue) from wild-type, endogenously tagged 3xFLAG::WAGO-1, and endogenously tagged 3xFLAG::GFP::WAGO-4 strains. *p < 0.05, ***p <0.0001, Two-way ANOVA with Tukey’s multiple comparison test.

**Supplemental Figure 11: N-terminus tagging of endogenous WAGO-1 cause transgenerational loss of *wago-1* gene expression and WAGO-1 protein expression. (A,B)** Representative western blot showing WAGO-1 expression in wild-type (left) and 3xFLAG::WAGO-1 adult hermaphrodites (right) that have been passaged for **(A)** 2 generations and **(B)** 100 generations. **(C,D)** Volcano plot displaying differential expression from mRNA-sequencing of wild-type and endogenously tagged 3xFLAG::WAGO-1 adult hermaphrodites after **(C)** two generations and **(D)** 100 generations. Blue denotes genes significantly depleted in 3xFLAG::WAGO-1 hermaphrodites compared to wild-type. Red dots represent gene significantly enriched in 3xFLAG::WAGO-1 hermaphrodites compared to wild-type. Differentially expressed AGOs labeled based on decreased expression compared to wild-type labeled in black.

**Supplemental Figure 12: Local electrostatic potential of wild-type and 3xFLAG::WAGO-1. (A-B)** Electrostatic map of AlphaFold3 predicted **(A)** wild-type WAGO-1 and **(B)** 3xFLAG::WAGO-1 structure created utilizing APBS and PDB2PQR Pymol extensions^100^. Potentials are on a −3.0 to 3.0 red-while-blue color map with units kJ/mol/e, where blue represents regions with a more positive charge and red represents regions with a more negative charge.

**Supplemental Figure 13: Histogram of average solvent accessible surface area (SASA) of wild-type and 3xFLAG::WAGO-1. (A)** Histogram depicting the average number of frames from three molecular dynamic simulations where the N-terminal lobe wild-type WAGO-1 (blue) and 3xFLAG::WAGO-1 (green) have a specific SASA. **(B)** Histogram depicting the average number of frames from three molecular dynamic simulations where the C-terminal lobe wild-type WAGO-1 (blue) and 3xFLAG::WAGO-1 (green) have a specific SASA.

**Supplemental Figure 14: Stills from three wild-type WAGO-1 molecular dynamic simulations.** Stills from three molecular dynamic simulations of wild-type WAGO-1 taken 50 nanoseconds apart. The IDR is depicted as ball-and-sphere model while the core of the protein is depicted as ribbon model. See Methods and Materials for determining starting configuration.

**Supplemental Figure 15: Stills from three 3xFLAG::WAGO-1 molecular dynamic simulations.** Stills from three molecular dynamic simulations of 3xFLAG::WAGO-1 taken 50 nanoseconds apart. The IDR is depicted as ball-and-sphere model while the core of the protein is depicted as ribbon model. See Methods and Materials for determining starting configuration.

**Supplemental Figure 16: *wago-1* (*tm1414*) causes a decrease in germ cell number in germlines undergoing oogenesis. (A)** Representative immunofluorescence images of nuclei in germlines undergoing oogenesis from wild-type and *wago-1(tm1414)* adult hermaphrodites. Scale bars represent 10 μm. **(B)** Bar graph showing average number of nuclei counts from halved wild-type *wago-1* and *wago-1(tm1414)* adult hermaphrodite germlines undergoing oogenesis. For each genotype n=9 gonads were analyzed. ****p<0.00001, one-way Anova.

**Supplemental Figure 17: Endogenously tagged 3xFLAG::WAGO-1 males misregulates gene expression compared to wildtype males (A)** Volcano plot of mRNA-sequencing data between wild-type and endogenously tagged 3xFLAG::WAGO-1 males. Genes depleted in 3xFLAG::WAGO-1 males are in blue. Genes enriched in 3xFLAG::WAGO-1 males are in red. Genes with less than a log_2_ Fold Change of 1 and a p-adjusted value less than 0.05 are in blue-gray. **(B)** Heat map depicting individual genes that are up (red) or down (blue) regulated based on log_2_ Fold Change in 3xFLAG::WAGO-1 compared to wild-type WAGO-1 grouped by gene ontology (GO) terms. GO groups are vertically sorted in descending order based on the proportion of genes that are upregulated. GO terms were assigned using WormCat 2.0^115^. White indicates no significant change in expression.

**Supplemental Figure 18: Top 15 GO terms that are misregulated in 3xFLAG::WAGO-1 strains. (A-C)** Top 15 gene ontology (GO) terms of genes misregulated in **(A)** adult hermaphrodites with 3xFLAG::WAGO-1 transgene, **(B)** adult hermaphrodites with 3xFLAG::WAGO-1 endogenously tagged, **(C)** adult males with 3xFLAG::WAGO-1 endogenously compared to wild-type adult hermaphrodites or males. GO terms are vertically sorted in descending order based on the number of genes misregulated. GO terms were assigned using WormCat 2.0 ^115^.

**Supplemental Figure 19: Global small RNA profile of wild-type and 3xFLAG::WAGO-1 tagged strains. (A-E)** Small RNAs grouped by nucleotide length from 18 to 30, with percent sRNAs with an A (purple), T/U (light blue), C (coral), or G (teal) on the 5’ end for **(A)** wild-type adult hermaphrodites, **(B)** 3xFLAG::WAGO-1 endogenously tagged adult hermaphrodites, **(C)** adult hermaphrodites with 3xFLAG::WAGO-1 transgene, **(D)** wild-type adult males, and **(E)** 3xFLAG::WAGO-endogenously tagged adult males.

**Supplemental Figure 20: Global small RNA profile of wild-type and silenced *wago-1* 3xFLAG::WAGO-1 tagged strains. (A-F)** Small RNAs grouped by nucleotide length from 18 to 30, with percent sRNAs with an A (purple), T/U (light blue), C (coral), or G (teal) on the 5’ end for **(A)** wild-type adult hermaphrodites, **(B)** 3xFLAG::WAGO-1 endogenously tagged adult hermaphrodites passaged for two generations, **(C)** 3xFLAG::WAGO-1 endogenously tagged adult hermaphrodites passaged for 100 generations, **(D)** wild-type adult males, **(E)** 3xFLAG::WAGO-1 endogenously tagged adult males passaged for two generations, and **(F)** 3xFLAG::WAGO-1 endogenously tagged adult males passaged for 100 generations.

**Supplemental Figure 21: Proportion of small RNAs mapping to specific genetic elements in silenced *wago-1* 3xFLAG::WAGO-1 strains. (A-F)** Pie charts depicting the proportion of **(A-B)** 21U-, **(C-D)** 22G-, and **(E-F)** 26G-RNAs mapping to specific genetic elements in wild-type, 3xFLAG::WAGO-1 passaged for two generations, and 3xFLAG::WAGO-1 passaged for 100 generations **(A,C,E)** adult hermaphrodites and **(B,D,F)** adult males. Small RNAs derived from highly translated RNAs, including AS rRNA and miRNA, as well as sRNAs sense to protein coding, pseudogenes, non-coding RNA, snoRNA, tRNA, and lincRNA were excluded (see Materials and Methods). AS = antisense, S=sense.

**Supplemental Figure 22: Global small RNA profile of wild-type and *wago-1 (tm1414)* hermaphrodites and males. (A-D)** Small RNAs grouped by nucleotide length from 18 to 30, with percent sRNAs with an A (purple), T/U (light blue), C (coral), or G (teal) on the 5’ end for **(A)** wild-type adult hermaphrodites, **(B)** *wago-1 (tm1414)* adult hermaphrodites, **(C)** wild-type adult males, and **(D)** *wago-1 (tm1414)* adult males.

## Supplemental Tables

**Table S1:** Means and standard deviations of WAGO-1 foci volumes throughout meiotic progression. Foci are counted if equal to or greater than 0.034 μm^2^ and less than 10 μm^2^. Hermaphrodite germlines n=9, male germlines n=9.

| Sex | Stage | Mean Volume (μm<sup>2</sup>) | STD | Number of foci |
| --- | --- | --- | --- | --- |
| Hermaphrodite | PMT | 0.124 | 0.17 | 5722 |
| Male | PMT | 0.126 | 0.12 | 7116 |
| Hermaphrodite | L/Z | 0.117 | 0.15 | 6448 |
| Male | L/Z | 0.172 | 0.17 | 5959 |
| Hermaphrodite | EP | 0.143 | 0.15 | 7014 |
| Male | EP | 0.194 | 0.20 | 4449 |
| Hermaphrodite | MP | 0.168 | 0.23 | 7114 |
| Male | MP | 0.237 | 0.31 | 3913 |
| Hermaphrodite | LP | 0.261 | 0.37 | 4552 |
| Male | LP | 0.274 | 0.36 | 3775 |

**Table S2:** Mann-Whitney U test comparing WAGO-1 foci volume distribution between hermaphrodites and males throughout meiosis.

| Stage | Statistic | p-value |
| --- | --- | --- |
| PMT | 18166008.5 | 4.04E-26 |
| L/Z | 14258864 | 1.33E-136 |
| EP | 13047742.5 | 7.62E-50 |
| MP | 12160562 | 2.11E-28 |
| LP | 8221293.0 | 0.00034 |

**Table S3:** Means and standard deviations of WAGO-1 foci sphericity throughout meiotic progression. Foci are counted if volumes are equal to or greater than 0.034 μm^2^ and less than 10 μm^2^.

| Sex | Stage | Mean ($\mu\text{m}^3$) | STD |
| --- | --- | --- | --- |
| Hermaphrodite | PMT | 0.738 | 0.101 |
| Male | PMT | 0.680 | 0.101 |
| Hermaphrodite | L/Z | 0.734 | 0.101 |
| Male | L/Z | 0.698 | 0.103 |
| Hermaphrodite | EP | 0.732 | 0.093 |
| Male | EP | 0.704 | 0.101 |
| Hermaphrodite | MP | 0.731 | 0.096 |
| Male | MP | 0.706 | 0.101 |
| Hermaphrodite | LP | 0.715 | 0.100 |
| Male | LP | 0.702 | 0.105 |

**Table S4:** Mann-Whitney U test comparing WAGO-1 foci sphericity between hermaphrodites and males throughout meiosis.

| Stage | Statistic | p-value |
| --- | --- | --- |
| PMT | 13774769 | 1.05E-218 |
| L/Z | 15217242.0 | 1.22E-89 |
| EP | 12992480.5 | 6.22E-52 |
| MP | 11715351.0 | 1.81E-43 |
| LP | 7883386.0 | 4.35E-11 |

**Table S5:** Binning of WAGO-1 overlap with PGL-1 and ZNFX-1 during meiotic progression.

| **Process** | **Classification** | **Protein** | **Zone** | **Foci Counts** | **Percent** |
| --- | --- | --- | --- | --- | --- |
| Oogenesis | 0% | PGL-1 | L/Z | 3128 | 0.498 |
| Oogenesis | 1-25% | PGL-1 | L/Z | 1775 | 0.283 |
| Oogenesis | 26-50% | PGL-1 | L/Z | 780 | 0.124 |
| Oogenesis | 51-75% | PGL-1 | L/Z | 396 | 0.063 |
| Oogenesis | 76-99% | PGL-1 | L/Z | 190 | 0.030 |
| Oogenesis | 100% | PGL-1 | L/Z | 10 | 0.002 |
| Oogenesis | Total | PGL-1 | L/Z | 6279 | 1.000 |
| Spermatogenesis | 0% | PGL-1 | L/Z | 1787 | 0.328 |
| Spermatogenesis | 1-25% | PGL-1 | L/Z | 1373 | 0.252 |
| Spermatogenesis | 26-50% | PGL-1 | L/Z | 1035 | 0.190 |
| Spermatogenesis | 51-75% | PGL-1 | L/Z | 808 | 0.148 |
| Spermatogenesis | 76-99% | PGL-1 | L/Z | 422 | 0.078 |
| Spermatogenesis | 100% | PGL-1 | L/Z | 18 | 0.003 |
| Spermatogenesis | Total | PGL-1 | L/Z | 5443 | 1.000 |
| Oogenesis | 0% | PGL-1 | EP | 2120 | 0.420 |
| Oogenesis | 1-25% | PGL-1 | EP | 1574 | 0.312 |
| Oogenesis | 26-50% | PGL-1 | EP | 768 | 0.152 |
| Oogenesis | 51-75% | PGL-1 | EP | 436 | 0.086 |
| Oogenesis | 76-99% | PGL-1 | EP | 144 | 0.029 |
| Oogenesis | 100% | PGL-1 | EP | 8 | 0.002 |
| Oogenesis | Total | PGL-1 | EP | 5050 | 1.000 |
| Spermatogenesis | 0% | PGL-1 | EP | 960 | 0.389 |
| Spermatogenesis | 1-25% | PGL-1 | EP | 545 | 0.221 |
| Spermatogenesis | 26-50% | PGL-1 | EP | 372 | 0.151 |
| Spermatogenesis | 51-75% | PGL-1 | EP | 386 | 0.156 |
| Spermatogenesis | 76-99% | PGL-1 | EP | 198 | 0.080 |
| Spermatogenesis | 100% | PGL-1 | EP | 7 | 0.003 |
| Spermatogenesis | Total | PGL-1 | EP | 2468 | 1.000 |
| Oogenesis | 0% | PGL-1 | LP | 1966 | 0.361 |
| Oogenesis | 1-25% | PGL-1 | LP | 1778 | 0.327 |
| Oogenesis | 26-50% | PGL-1 | LP | 963 | 0.177 |
| Oogenesis | 51-75% | PGL-1 | LP | 458 | 0.084 |
| Oogenesis | 76-99% | PGL-1 | LP | 264 | 0.048 |
| Oogenesis | 100% | PGL-1 | LP | 15 | 0.003 |
| Oogenesis | Total | PGL-1 | LP | 5444 | 1.000 |
| Spermatogenesis | 0% | PGL-1 | LP | 617 | 0.309 |
| Spermatogenesis | 1-25% | PGL-1 | LP | 459 | 0.230 |
| Spermatogenesis | 26-50% | PGL-1 | LP | 292 | 0.146 |
| Spermatogenesis | 51-75% | PGL-1 | LP | 422 | 0.211 |
| Spermatogenesis | 76-99% | PGL-1 | LP | 200 | 0.100 |
| Spermatogenesis | 100% | PGL-1 | LP | 10 | 0.005 |
| Spermatogenesis | Total | PGL-1 | LP | 2000 | 1.000 |
| Oogenesis | 0% | ZNFX-1 | L/Z | 2691 | 0.380 |
| Oogenesis | 1-25% | ZNFX-1 | L/Z | 1766 | 0.249 |
| Oogenesis | 26-50% | ZNFX-1 | L/Z | 1350 | 0.191 |
| Oogenesis | 51-75% | ZNFX-1 | L/Z | 918 | 0.130 |
| Oogenesis | 76-99% | ZNFX-1 | L/Z | 349 | 0.049 |
| Oogenesis | 100% | ZNFX-1 | L/Z | 11 | 0.002 |
| Oogenesis | Total | ZNFX-1 | L/Z | 7085 | 1.000 |
| Spermatogenesis | 0% | ZNFX-1 | L/Z | 1784 | 0.304 |
| Spermatogenesis | 1-25% | ZNFX-1 | L/Z | 1092 | 0.186 |
| Spermatogenesis | 26-50% | ZNFX-1 | L/Z | 1088 | 0.185 |
| Spermatogenesis | 51-75% | ZNFX-1 | L/Z | 1089 | 0.186 |
| Spermatogenesis | 76-99% | ZNFX-1 | L/Z | 772 | 0.132 |
| Spermatogenesis | 100% | ZNFX-1 | L/Z | 45 | 0.008 |
| Spermatogenesis | Total | ZNFX-1 | L/Z | 5870 | 1.000 |
| Oogenesis | 0% | ZNFX-1 | EP | 1401 | 0.260 |
| Oogenesis | 1-25% | ZNFX-1 | EP | 1425 | 0.265 |
| Oogenesis | 26-50% | ZNFX-1 | EP | 1269 | 0.236 |
| Oogenesis | 51-75% | ZNFX-1 | EP | 876 | 0.163 |
| Oogenesis | 76-99% | ZNFX-1 | EP | 396 | 0.074 |
| Oogenesis | 100% | ZNFX-1 | EP | 14 | 0.003 |
| Oogenesis | Total | ZNFX-1 | EP | 5381 | 1.000 |
| Spermatogenesis | 0% | ZNFX-1 | EP | 895 | 0.317 |
| Spermatogenesis | 1-25% | ZNFX-1 | EP | 450 | 0.159 |
| Spermatogenesis | 26-50% | ZNFX-1 | EP | 459 | 0.162 |
| Spermatogenesis | 51-75% | ZNFX-1 | EP | 552 | 0.195 |
| Spermatogenesis | 76-99% | ZNFX-1 | EP | 453 | 0.160 |
| Spermatogenesis | 100% | ZNFX-1 | EP | 18 | 0.006 |
| Spermatogenesis | Total | ZNFX-1 | EP | 2827 | 1.000 |
| Oogenesis | 0% | ZNFX-1 | LP | 1550 | 0.266 |
| Oogenesis | 1-25% | ZNFX-1 | LP | 1741 | 0.299 |
| Oogenesis | 26-50% | ZNFX-1 | LP | 1442 | 0.248 |
| Oogenesis | 51-75% | ZNFX-1 | LP | 752 | 0.129 |
| Oogenesis | 76-99% | ZNFX-1 | LP | 306 | 0.053 |
| Oogenesis | 100% | ZNFX-1 | LP | 33 | 0.006 |
| Oogenesis | Total | ZNFX-1 | LP | 5824 | 1.000 |
| Spermatogenesis | 0% | ZNFX-1 | LP | 570 | 0.245 |
| Spermatogenesis | 1-25% | ZNFX-1 | LP | 290 | 0.125 |
| Spermatogenesis | 26-50% | ZNFX-1 | LP | 359 | 0.154 |
| Spermatogenesis | 51-75% | ZNFX-1 | LP | 545 | 0.234 |
| Spermatogenesis | 76-99% | ZNFX-1 | LP | 528 | 0.227 |
| Spermatogenesis | 100% | ZNFX-1 | LP | 37 | 0.016 |
| Spermatogenesis | Total | ZNFX-1 | LP | 2329 | 1.000 |

**Table S6:** Sex-specific comparisons of WAGO-1 overlap with PGL-1 or ZNFX-1 through meiotic progression.

| **Comparison** | **Classification** | **Protein** | **Zone** | **Chi-squared statistic** | **p-value** |
| --- | --- | --- | --- | --- | --- |
| Oogenesis vs. Spermatogenesis | 0% | PGL-1 | L/Z | 344.78 | 2.91E-76 |
| Oogenesis vs. Spermatogenesis | 1-25% | PGL-1 | L/Z | 13.6 | 1.15E-03 |
| Oogenesis vs. Spermatogenesis | 26-50% | PGL-1 | L/Z | 96.34 | 4.84E-22 |
| Oogenesis vs. Spermatogenesis | 51-75% | PGL-1 | L/Z | 229.69 | 3.49E-51 |
| Oogenesis vs. Spermatogenesis | 76-99% | PGL-1 | L/Z | 130.71 | 1.44E-29 |
| Oogenesis vs. Spermatogenesis | 100% | PGL-1 | L/Z | 2.91 | 4.40E-01 |
| Oogenesis vs. Spermatogenesis | 0% | PGL-1 | EP | 6.39 | 5.75E-02 |
| Oogenesis vs. Spermatogenesis | 1-25% | PGL-1 | EP | 67.16 | 1.25E-15 |
| Oogenesis vs. Spermatogenesis | 26-50% | PGL-1 | EP | 0.014 | 4.53E+00 |
| Oogenesis vs. Spermatogenesis | 51-75% | PGL-1 | EP | 83.36 | 3.43E-19 |
| Oogenesis vs. Spermatogenesis | 76-99% | PGL-1 | EP | 100.91 | 4.82E-23 |
| Oogenesis vs. Spermatogenesis | 100% | PGL-1 | EP | 0.75 | 1.98E+00 |
| Oogenesis vs. Spermatogenesis | 0% | PGL-1 | LP | 17.64 | 1.33E-04 |
| Oogenesis vs. Spermatogenesis | 1-25% | PGL-1 | LP | 65.14 | 3.49E-15 |
| Oogenesis vs. Spermatogenesis | 26-50% | PGL-1 | LP | 9.74 | 9.00E-03 |
| Oogenesis vs. Spermatogenesis | 51-75% | PGL-1 | LP | 224.63 | 4.41E-50 |
| Oogenesis vs. Spermatogenesis | 76-99% | PGL-1 | LP | 65.51 | 2.89E-15 |
| Oogenesis vs. Spermatogenesis | 100% | PGL-1 | LP | 1.58 | 1.05E+00 |
| Oogenesis vs. Spermatogenesis | 0% | ZNFX-1 | L/Z | 81.33 | 9.55E-19 |
| Oogenesis vs. Spermatogenesis | 1-25% | ZNFX-1 | L/Z | 74.19 | 3.55E-17 |
| Oogenesis vs. Spermatogenesis | 26-50% | ZNFX-1 | L/Z | 0.53 | 2.34E+00 |
| Oogenesis vs. Spermatogenesis | 51-75% | ZNFX-1 | L/Z | 76.41 | 1.16E-17 |
| Oogenesis vs. Spermatogenesis | 76-99% | ZNFX-1 | L/Z | 274.12 | 7.20E-61 |
| Oogenesis vs. Spermatogenesis | 100% | ZNFX-1 | L/Z | 20 | 3.87E-05 |
| Oogenesis vs. Spermatogenesis | 0% | ZNFX-1 | EP | 28.37 | 5.00E-07 |
| Oogenesis vs. Spermatogenesis | 1-25% | ZNFX-1 | EP | 116.75 | 1.63E-26 |
| Oogenesis vs. Spermatogenesis | 26-50% | ZNFX-1 | EP | 59.74 | 5.40E-14 |
| Oogenesis vs. Spermatogenesis | 51-75% | ZNFX-1 | EP | 13.38 | 1.30E-03 |
| Oogenesis vs. Spermatogenesis | 76-99% | ZNFX-1 | EP | 149.11 | 1.36E-33 |
| Oogenesis vs. Spermatogenesis | 100% | ZNFX-1 | EP | 5.83 | 8.00E-02 |
| Oogenesis vs. Spermatogenesis | 0% | ZNFX-1 | LP | 3.85 | 2.50E-01 |
| Oogenesis vs. Spermatogenesis | 1-25% | ZNFX-1 | LP | 296.64 | 6.80E-60 |
| Oogenesis vs. Spermatogenesis | 26-50% | ZNFX-1 | LP | 83.88 | 2.63E-19 |
| Oogenesis vs. Spermatogenesis | 51-75% | ZNFX-1 | LP | 136.03 | 9.85E-31 |
| Oogenesis vs. Spermatogenesis | 76-99% | ZNFX-1 | LP | 547.66 | 2.03E-120 |
| Oogenesis vs. Spermatogenesis | 100% | ZNFX-1 | LP | 19.23 | 5.80E-05 |

**Table S7:** WAGO-1 foci overlap with ZNFX-1, PGL-1, or both germ granule components.

| Stage | Sex | ZNFX-1 | PGL-1 | Both | Total | % ZNFX-1 | % PGL-1 | % Both |
| --- | --- | --- | --- | --- | --- | --- | --- | --- |
| L/Z | Hermaphrodite | 1123 | 648 | 2329 | 4100 | 27.4 | 15.8 | 56.8 |
| L/Z | Male | 842 | 655 | 2571 | 4068 | 20.7 | 16.1 | 63.2 |
| EP | Hermaphrodite | 1065 | 263 | 2330 | 3658 | 29.1 | 7.2 | 63.7 |
| EP | Male | 476 | 254 | 1253 | 1983 | 24.0 | 12.8 | 63.2 |
| LP | Hermaphrodite | 868 | 406 | 2744 | 4018 | 21.6 | 10.1 | 68.3 |
| LP | Male | 400 | 160 | 1223 | 1783 | 22.4 | 9.0 | 68.6 |

**Table S8:** Sex-specific comparisons of percent WAGO-1 foci that overlap with PGL-1, ZNFX-1, or both foci.

| Comparison | Protein | Zone | Chi-squared statistic | p-value |
| --- | --- | --- | --- | --- |
| Oogenesis vs. Spermatogenesis | Both | L/Z | 34.53 | 1.2564E-08 |
| Oogenesis vs. Spermatogenesis | PGL-1 | L/Z | 59.17 | 4.32E-14 |
| Oogenesis vs. Spermatogenesis | ZNFX-1 | L/Z | 0.113 | 2.211 |
| Oogenesis vs. Spermatogenesis | Both | EP | 0.123 | 2.178 |
| Oogenesis vs. Spermatogenesis | PGL-1 | EP | 16.66 | 0.0001344 |
| Oogenesis vs. Spermatogenesis | ZNFX-1 | EP | 48.1 | 1.2159E-11 |
| Oogenesis vs. Spermatogenesis | Both | LP | 0.038 | 2.535 |
| Oogenesis vs. Spermatogenesis | PGL-1 | LP | 0.452 | 1.503 |
| Oogenesis vs. Spermatogenesis | ZNFX-1 | LP | 1.668 | 0.588 |

**Table S9:**
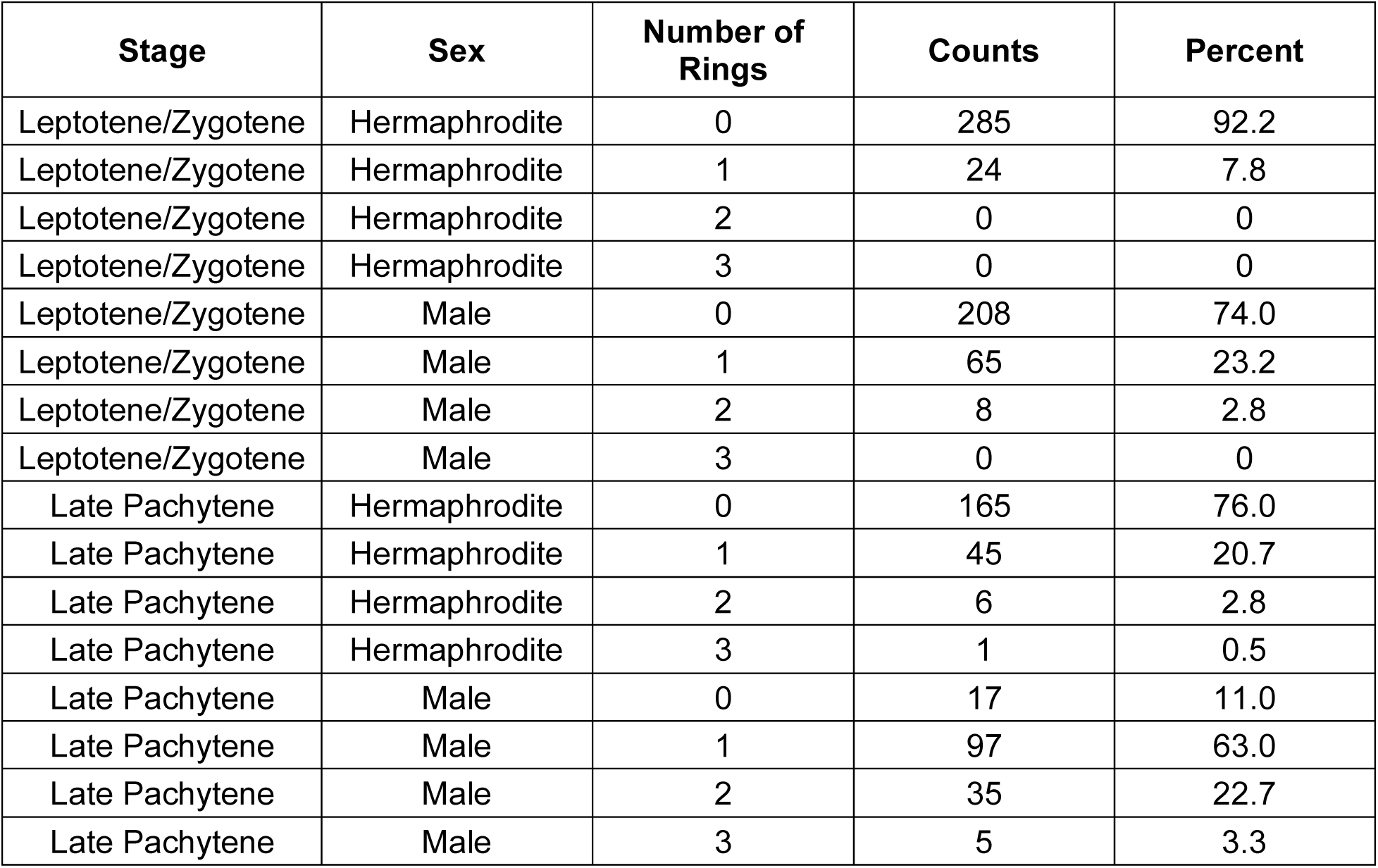
PRG-1 ring counts. Germlines n=6 per a sex taken for at least three distinct technical replicates.

| Stage | Sex | Number of Rings | Counts | Percent |
| --- | --- | --- | --- | --- |
| Leptotene/Zygotene | Hermaphrodite | 0 | 285 | 92.2 |
| Leptotene/Zygotene | Hermaphrodite | 1 | 24 | 7.8 |
| Leptotene/Zygotene | Hermaphrodite | 2 | 0 | 0 |
| Leptotene/Zygotene | Hermaphrodite | 3 | 0 | 0 |
| Leptotene/Zygotene | Male | 0 | 208 | 74.0 |
| Leptotene/Zygotene | Male | 1 | 65 | 23.2 |
| Leptotene/Zygotene | Male | 2 | 8 | 2.8 |
| Leptotene/Zygotene | Male | 3 | 0 | 0 |
| Late Pachytene | Hermaphrodite | 0 | 165 | 76.0 |
| Late Pachytene | Hermaphrodite | 1 | 45 | 20.7 |
| Late Pachytene | Hermaphrodite | 2 | 6 | 2.8 |
| Late Pachytene | Hermaphrodite | 3 | 1 | 0.5 |
| Late Pachytene | Male | 0 | 17 | 11.0 |
| Late Pachytene | Male | 1 | 97 | 63.0 |
| Late Pachytene | Male | 2 | 35 | 22.7 |
| Late Pachytene | Male | 3 | 5 | 3.3 |

**Table S10:** Sex specific normalized fertility of tagged WAGO-1 and WAGO-4 strains.

| Strain | Counts | Sex | Genotype | Average | Normalized Count |
| --- | --- | --- | --- | --- | --- |
| N2 | 318 | Hermaphrodite | Wild-type | 298.77 | 1.06 |
| N2 | 353 | Hermaphrodite | Wild-type |  | 1.18 |
| N2 | 310 | Hermaphrodite | Wild-type |  | 1.03 |
| N2 | 304 | Hermaphrodite | Wild-type |  | 1.02 |
| N2 | 346 | Hermaphrodite | Wild-type |  | 1.16 |
| N2 | 315 | Hermaphrodite | Wild-type |  | 1.05 |
| N2 | 345 | Hermaphrodite | Wild-type |  | 1.15 |
| N2 | 346 | Hermaphrodite | Wild-type |  | 1.16 |
| N2 | 346 | Hermaphrodite | Wild-type |  | 1.16 |
| N2 | 345 | Hermaphrodite | Wild-type |  | 1.15 |
| N2 | 337 | Hermaphrodite | Wild-type |  | 1.13 |
| N2 | 250 | Hermaphrodite | Wild-type |  | 0.84 |
| N2 | 314 | Hermaphrodite | Wild-type |  | 1.05 |
| N2 | 363 | Hermaphrodite | Wild-type |  | 1.21 |
| N2 | 321 | Hermaphrodite | Wild-type |  | 1.07 |
| N2 | 335 | Hermaphrodite | Wild-type |  | 1.12 |
| N2 | 293 | Hermaphrodite | Wild-type |  | 0.98 |
| N2 | 286 | Hermaphrodite | Wild-type |  | 0.96 |
| N2 | 218 | Hermaphrodite | Wild-type |  | 0.73 |
| N2 | 194 | Hermaphrodite | Wild-type |  | 0.65 |
| N2 | 246 | Hermaphrodite | Wild-type |  | 0.82 |
| N2 | 319 | Hermaphrodite | Wild-type |  | 1.07 |
| N2 | 226 | Hermaphrodite | Wild-type |  | 0.76 |
| N2 | 282 | Hermaphrodite | Wild-type |  | 0.94 |
| N2 | 196 | Hermaphrodite | Wild-type |  | 0.66 |
| N2 | 260 | Hermaphrodite | Wild-type |  | 0.87 |
| WM205 | 196 | Hermaphrodite | GFP::WAGO-1<br>endogenous | 199.69 | 0.66 |
| WM205 | 153 | Hermaphrodite | GFP::WAGO-1<br>endogenous |  | 0.51 |
| WM205 | 195 | Hermaphrodite | GFP::WAGO-1<br>endogenous |  | 0.65 |
| WM205 | 293 | Hermaphrodite | GFP::WAGO-1<br>endogenous |  | 0.98 |
| WM205 | 241 | Hermaphrodite | GFP::WAGO-1<br>endogenous |  | 0.81 |
| WM205 | 200 | Hermaphrodite | GFP::WAGO-1<br>endogenous |  | 0.67 |
| WM205 | 229 | Hermaphrodite | GFP::WAGO-1<br>endogenous |  | 0.77 |
| WM205 | 198 | Hermaphrodite | GFP::WAGO-1<br>endogenous |  | 0.66 |
| WM205 | 188 | Hermaphrodite | GFP::WAGO-1<br>endogenous |  | 0.63 |
| WM205 | 188 | Hermaphrodite | GFP::WAGO-1<br>endogenous |  | 0.63 |
| WM205 | 169 | Hermaphrodite | GFP::WAGO-1<br>endogenous |  | 0.57 |
| WM205 | 203 | Hermaphrodite | GFP::WAGO-1<br>endogenous |  | 0.68 |
| WM205 | 180 | Hermaphrodite | GFP::WAGO-1<br>endogenous |  | 0.60 |
| WM205 | 183 | Hermaphrodite | GFP::WAGO-1<br>endogenous |  | 0.61 |
| WM205 | 201 | Hermaphrodite | GFP::WAGO-1<br>endogenous |  | 0.67 |
| WM205 | 178 | Hermaphrodite | GFP::WAGO-1<br>endogenous |  | 0.60 |
| WM616 | 265 | Hermaphrodite | 3xFLAG::WAGO-1<br>endogenous | 264.06 | 0.89 |
| WM616 | 55 | Hermaphrodite | 3xFLAG::WAGO-1<br>endogenous |  | 0.18 |
| WM616 | 243 | Hermaphrodite | 3xFLAG::WAGO-1<br>endogenous |  | 0.81 |
| WM616 | 247 | Hermaphrodite | 3xFLAG::WAGO-1<br>endogenous |  | 0.83 |
| WM616 | 269 | Hermaphrodite | 3xFLAG::WAGO-1<br>endogenous |  | 0.90 |
| WM616 | 275 | Hermaphrodite | 3xFLAG::WAGO-1<br>endogenous |  | 0.92 |
| WM616 | 318 | Hermaphrodite | 3xFLAG::WAGO-1<br>endogenous |  | 1.06 |
| WM616 | 305 | Hermaphrodite | 3xFLAG::WAGO-1<br>endogenous |  | 1.02 |
| WM616 | 277 | Hermaphrodite | 3xFLAG::WAGO-1<br>endogenous |  | 0.93 |
| WM616 | 346 | Hermaphrodite | 3xFLAG::WAGO-1<br>endogenous |  | 1.16 |
| WM616 | 246 | Hermaphrodite | 3xFLAG::WAGO-1<br>endogenous |  | 0.82 |
| WM616 | 227 | Hermaphrodite | 3xFLAG::WAGO-1<br>endogenous |  | 0.76 |
| WM616 | 274 | Hermaphrodite | 3xFLAG::WAGO-1<br>endogenous |  | 0.92 |
| WM616 | 332 | Hermaphrodite | 3xFLAG::WAGO-1<br>endogenous |  | 1.11 |
| WM616 | 276 | Hermaphrodite | 3xFLAG::WAGO-1<br>endogenous |  | 0.92 |
| WM616 | 279 | Hermaphrodite | 3xFLAG::WAGO-1<br>endogenous |  | 0.93 |
| WM616 | 255 | Hermaphrodite | 3xFLAG::WAGO-1<br>endogenous |  | 0.85 |
| WM192 | 214 | Hermaphrodite | 3xFLAG::WAGO-1<br>transgene | 208.20 | 0.72 |
| WM192 | 165 | Hermaphrodite | 3xFLAG::WAGO-1<br>transgene |  | 0.55 |
| WM192 | 183 | Hermaphrodite | 3xFLAG::WAGO-1<br>transgene |  | 0.61 |
| WM192 | 305 | Hermaphrodite | 3xFLAG::WAGO-1<br>transgene |  | 1.02 |
| WM192 | 247 | Hermaphrodite | 3xFLAG::WAGO-1<br>transgene |  | 0.83 |
| WM192 | 219 | Hermaphrodite | 3xFLAG::WAGO-1<br>transgene |  | 0.73 |
| WM192 | 211 | Hermaphrodite | 3xFLAG::WAGO-1<br>transgene |  | 0.71 |
| WM192 | 199 | Hermaphrodite | 3xFLAG::WAGO-1<br>transgene |  | 0.67 |
| WM192 | 190 | Hermaphrodite | 3xFLAG::WAGO-1<br>transgene |  | 0.64 |
| WM192 | 214 | Hermaphrodite | 3xFLAG::WAGO-1<br>transgene |  | 0.72 |
| WM192 | 194 | Hermaphrodite | 3xFLAG::WAGO-1<br>transgene |  | 0.65 |
| WM192 | 175 | Hermaphrodite | 3xFLAG::WAGO-1<br>transgene |  | 0.59 |
| WM192 | 240 | Hermaphrodite | 3xFLAG::WAGO-1<br>transgene |  | 0.80 |
| WM192 | 188 | Hermaphrodite | 3xFLAG::WAGO-1<br>transgene |  | 0.63 |
| WM192 | 203 | Hermaphrodite | 3xFLAG::WAGO-1<br>transgene |  | 0.68 |
| WM192 | 180 | Hermaphrodite | 3xFLAG::WAGO-1<br>transgene |  | 0.60 |
| WM192 | 234 | Hermaphrodite | 3xFLAG::WAGO-1 transgene |  | 0.78 |
| WM192 | 191 | Hermaphrodite | 3xFLAG::WAGO-1 transgene |  | 0.64 |
| WM192 | 216 | Hermaphrodite | 3xFLAG::WAGO-1 transgene |  | 0.72 |
| WM192 | 196 | Hermaphrodite | 3xFLAG::WAGO-1 transgene |  | 0.66 |
| YY1325 | 322 | Hermaphrodite | GFP::WAGO-4 endogenous | 276.31 | 1.08 |
| YY1325 | 295 | Hermaphrodite | GFP::WAGO-4 endogenous |  | 0.99 |
| YY1325 | 277 | Hermaphrodite | GFP::WAGO-4 endogenous |  | 0.93 |
| YY1325 | 239 | Hermaphrodite | GFP::WAGO-4 endogenous |  | 0.80 |
| YY1325 | 172 | Hermaphrodite | GFP::WAGO-4 endogenous |  | 0.56 |
| YY1325 | 279 | Hermaphrodite | GFP::WAGO-4 endogenous |  | 0.93 |
| YY1325 | 292 | Hermaphrodite | GFP::WAGO-4 endogenous |  | 0.98 |
| YY1325 | 350 | Hermaphrodite | GFP::WAGO-4 endogenous |  | 1.17 |
| YY1325 | 282 | Hermaphrodite | GFP::WAGO-4 endogenous |  | 0.94 |
| YY1325 | 325 | Hermaphrodite | GFP::WAGO-4 endogenous |  | 1.09 |
| YY1325 | 279 | Hermaphrodite | GFP::WAGO-4 endogenous |  | 0.93 |
| YY1325 | 219 | Hermaphrodite | GFP::WAGO-4 endogenous |  | 0.73 |
| YY1325 | 327 | Hermaphrodite | GFP::WAGO-4<br>endogenous |  | 1.09 |
| YY1325 | 326 | Hermaphrodite | GFP::WAGO-4<br>endogenous |  | 1.09 |
| YY1325 | 259 | Hermaphrodite | GFP::WAGO-4<br>endogenous |  | 0.87 |
| YY1325 | 178 | Hermaphrodite | GFP::WAGO-4<br>endogenous |  | 0.60 |
| N2 | 520 | Male | Wild-type | 401.67 | 1.29 |
| N2 | 413 | Male | Wild-type |  | 1.03 |
| N2 | 428 | Male | Wild-type |  | 1.07 |
| N2 | 303 | Male | Wild-type |  | 0.75 |
| N2 | 305 | Male | Wild-type |  | 0.76 |
| N2 | 441 | Male | Wild-type |  | 1.10 |
| WM205 | 67 | Male | GFP::WAGO-1<br>endogenous | 112.00 | 0.17 |
| WM205 | 157 | Male | GFP::WAGO-1<br>endogenous |  | 0.39 |
| WM616 | 254 | Male | 3xFLAG::WAGO-1<br>endogenous | 228.00 | 0.63 |
| WM616 | 331 | Male | 3xFLAG::WAGO-1<br>endogenous |  | 0.82 |
| WM616 | 99 | Male | 3xFLAG::WAGO-1<br>endogenous |  | 0.25 |
| WM192 | 72 | Male | 3xFLAG::WAGO-1<br>transgene | 172.33 | 0.18 |
| WM192 | 445 | Male | 3xFLAG::WAGO-1<br>transgene |  | 1.11 |
| WM192 | 0 | Male | 3xFLAG::WAGO-1<br>transgene |  | 0 |
| YY1325 | 0 | Male | GFP::WAGO-4<br>endogenous | 41.00 | 0 |
| YY1325 | 2 | Male | GFP::WAGO-4<br>endogenous |  | 0.005 |
| YY1325 | 121 | Male | GFP::WAGO-4<br>endogenous |  | 0.30 |

**Table S11:** Sex specific normalized fertility of wild-type versus C-terminal truncated WAGO-1.

| Strain | Counts | Sex | Genotype | Average | Normalized Count |
| --- | --- | --- | --- | --- | --- |
| N2 | 318 | Hermaphrodite | Wild-type | 298.77 | 1.06 |
| N2 | 353 | Hermaphrodite | Wild-type |  | 1.18 |
| N2 | 310 | Hermaphrodite | Wild-type |  | 1.04 |
| N2 | 304 | Hermaphrodite | Wild-type |  | 1.02 |
| N2 | 346 | Hermaphrodite | Wild-type |  | 1.16 |
| N2 | 315 | Hermaphrodite | Wild-type |  | 1.05 |
| N2 | 345 | Hermaphrodite | Wild-type |  | 1.15 |
| N2 | 346 | Hermaphrodite | Wild-type |  | 1.16 |
| N2 | 346 | Hermaphrodite | Wild-type |  | 1.16 |
| N2 | 345 | Hermaphrodite | Wild-type |  | 1.15 |
| N2 | 337 | Hermaphrodite | Wild-type |  | 1.13 |
| N2 | 250 | Hermaphrodite | Wild-type |  | 0.84 |
| N2 | 314 | Hermaphrodite | Wild-type |  | 1.05 |
| N2 | 363 | Hermaphrodite | Wild-type |  | 1.21 |
| N2 | 321 | Hermaphrodite | Wild-type |  | 1.07 |
| N2 | 335 | Hermaphrodite | Wild-type |  | 1.12 |
| N2 | 293 | Hermaphrodite | Wild-type |  | 0.98 |
| N2 | 286 | Hermaphrodite | Wild-type |  | 0.96 |
| N2 | 218 | Hermaphrodite | Wild-type |  | 0.73 |
| N2 | 194 | Hermaphrodite | Wild-type |  | 0.65 |
| N2 | 246 | Hermaphrodite | Wild-type |  | 0.82 |
| N2 | 319 | Hermaphrodite | Wild-type |  | 1.07 |
| N2 | 226 | Hermaphrodite | Wild-type |  | 0.76 |
| N2 | 282 | Hermaphrodite | Wild-type |  | 0.94 |
| N2 | 196 | Hermaphrodite | Wild-type |  | 0.66 |
| N2 | 260 | Hermaphrodite | Wild-type |  | 0.87 |
| C184 | 193 | Hermaphrodite | *wago-1 (tm1414)* | 216.29 | 0.65 |
| C184 | 0 | Hermaphrodite | *wago-1 (tm1414)* |  | 0.00 |
| C184 | 119 | Hermaphrodite | *wago-1 (tm1414)* |  | 0.40 |
| C184 | 302 | Hermaphrodite | *wago-1 (tm1414)* |  | 1.01 |
| C184 | 144 | Hermaphrodite | *wago-1 (tm1414)* |  | 0.48 |
| C184 | 189 | Hermaphrodite | *wago-1 (tm1414)* |  | 0.63 |
| C184 | 279 | Hermaphrodite | *wago-1 (tm1414)* |  | 0.93 |
| C184 | 292 | Hermaphrodite | *wago-1 (tm1414)* |  | 0.98 |
| C184 | 273 | Hermaphrodite | *wago-1 (tm1414)* |  | 0.91 |
| C184 | 258 | Hermaphrodite | *wago-1 (tm1414)* |  | 0.86 |
| C184 | 258 | Hermaphrodite | *wago-1 (tm1414)* |  | 0.86 |
| C184 | 210 | Hermaphrodite | *wago-1 (tm1414)* |  | 0.70 |
| C184 | 244 | Hermaphrodite | *wago-1 (tm1414)* |  | 0.82 |
| C184 | 254 | Hermaphrodite | *wago-1 (tm1414)* |  | 0.85 |
| C184 | 249 | Hermaphrodite | *wago-1 (tm1414)* |  | 0.83 |
| C184 | 170 | Hermaphrodite | *wago-1 (tm1414)* |  | 0.57 |
| C184 | 243 | Hermaphrodite | *wago-1 (tm1414)* |  | 0.81 |
| N2 | 520 | Male | Wild-type | 401.67 | 1.29 |
| N2 | 413 | Male | Wild-type |  | 1.03 |
| N2 | 428 | Male | Wild-type |  | 1.07 |
| N2 | 303 | Male | Wild-type |  | 0.75 |
| N2 | 305 | Male | Wild-type |  | 0.76 |
| N2 | 441 | Male | Wild-type |  | 1.10 |
| C184 | 0 | Male | *wago-1 (tm1414)* | 0.00 | 0.00 |
| C184 | 0 | Male | *wago-1 (tm1414)* |  | 0.00 |
| C184 | 0 | Male | *wago-1 (tm1414)* |  | 0.00 |

**Table S12:** Binning of WAGO-1 overlap with PGL-1 during meiotic progression in wildtype and truncated *wago-1(tm1414)*

| Genotype | Classification | Protein | Zone | Foci Counts | Percent |
| --- | --- | --- | --- | --- | --- |
| Wild-type | 0% | PGL-1 | L/Z | 3128 | 0.498 |
| Wild-type | 1-25% | PGL-1 | L/Z | 1775 | 0.283 |
| Wild-type | 26-50% | PGL-1 | L/Z | 780 | 0.124 |
| Wild-type | 51-75% | PGL-1 | L/Z | 396 | 0.063 |
| Wild-type | 76-99% | PGL-1 | L/Z | 190 | 0.030 |
| Wild-type | 100% | PGL-1 | L/Z | 10 | 0.002 |
| Wild-type | Total | PGL-1 | L/Z | 6279 | 1.000 |
| *wago-1 (tm1414)* | 0% | PGL-1 | L/Z | 1721 | 0.800 |
| *wago-1 (tm1414)* | 1-25% | PGL-1 | L/Z | 222 | 0.103 |
| *wago-1 (tm1414)* | 26-50% | PGL-1 | L/Z | 92 | 0.043 |
| *wago-1 (tm1414)* | 51-75% | PGL-1 | L/Z | 62 | 0.029 |
| *wago-1 (tm1414)* | 76-99% | PGL-1 | L/Z | 40 | 0.019 |
| *wago-1 (tm1414)* | 100% | PGL-1 | L/Z | 15 | 0.007 |
| *wago-1 (tm1414)* | Total | PGL-1 | L/Z | 2152 | 1.000 |
| Wild-type | 0% | PGL-1 | Pachytene | 4086 | 0.389 |
| Wild-type | 1-25% | PGL-1 | Pachytene | 3352 | 0.319 |
| Wild-type | 26-50% | PGL-1 | Pachytene | 1731 | 0.165 |
| Wild-type | 51-75% | PGL-1 | Pachytene | 894 | 0.085 |
| Wild-type | 76-99% | PGL-1 | Pachytene | 408 | 0.039 |
| Wild-type | 100% | PGL-1 | Pachytene | 23 | 0.002 |
| Wild-type | Total | PGL-1 | Pachytene | 10494 | 1.000 |
| *wago-1 (tm1414)* | 0% | PGL-1 | Pachytene | 2690 | 0.771 |
| *wago-1 (tm1414)* | 1-25% | PGL-1 | Pachytene | 318 | 0.091 |
| *wago-1 (tm1414)* | 26-50% | PGL-1 | Pachytene | 199 | 0.057 |
| *wago-1 (tm1414)* | 51-75% | PGL-1 | Pachytene | 154 | 0.044 |
| *wago-1 (tm1414)* | 76-99% | PGL-1 | Pachytene | 80 | 0.023 |
| *wago-1 (tm1414)* | 100% | PGL-1 | Pachytene | 47 | 0.013 |
| *wago-1 (tm1414)* | Total | PGL-1 | Pachytene | 3488 | 1.000 |

**Table S13:** Wild-type versus truncated *wago-1 (tm1414)* comparisons of WAGO-1 overlap with PGL-1 through meiotic progression.

| Comparison | Classification | Protein | Zone | Chi-squared statistic | p-Value |
| --- | --- | --- | --- | --- | --- |
| Wild-type vs. *wago-1(tm1414)* | 0% | PGL-1 | L/Z | 59.556 | 1.18E-14 |
| Wild-type vs. *wago-1(tm1414)* | 1-25% | PGL-1 | L/Z | 36.49 | 1.53E-09 |
| Wild-type vs. *wago-1(tm1414)* | 26-50% | PGL-1 | L/Z | 5.85 | 0.0156 |
| Wild-type vs. *wago-1(tm1414)* | 51-75% | PGL-1 | L/Z | 9.81 | 0.0017 |
| Wild-type vs. *wago-1(tm1414)* | 76-99% | PGL-1 | L/Z | 6.46 | 0.011 |
| Wild-type vs. *wago-1(tm1414)* | 100% | PGL-1 | L/Z | 3.2 | 0.074 |
| Wild-type vs. *wago-1(tm1414)* | 0% | PGL-1 | Pachytene | 460.07 | 4.63E-102 |
| Wild-type vs. *wago-1(tm1414)* | 1-25% | PGL-1 | Pachytene | 240.43 | 3.17E-54 |
| Wild-type vs. *wago-1(tm1414)* | 26-50% | PGL-1 | Pachytene | 72.14 | 2.00E-17 |
| Wild-type vs. *wago-1(tm1414)* | 51-75% | PGL-1 | Pachytene | 40.71 | 1.77E-10 |
| Wild-type vs. *wago-1(tm1414)* | 76-99% | PGL-1 | Pachytene | 22.26 | 2.38E-06 |
| Wild-type vs. *wago-1(tm1414)* | 100% | PGL-1 | Pachytene | 21.28 | 3.96E-06 |

**Table S14:**
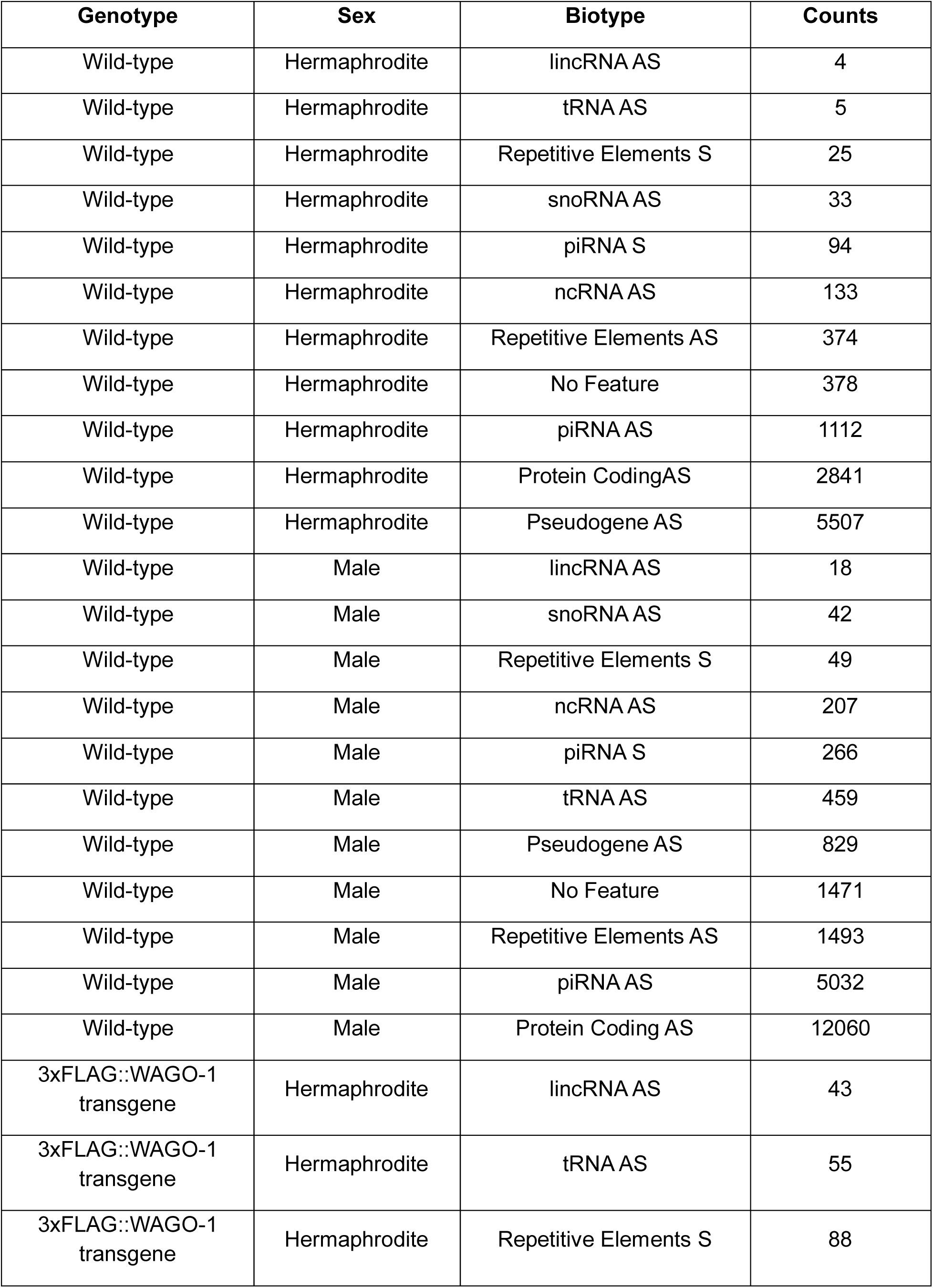
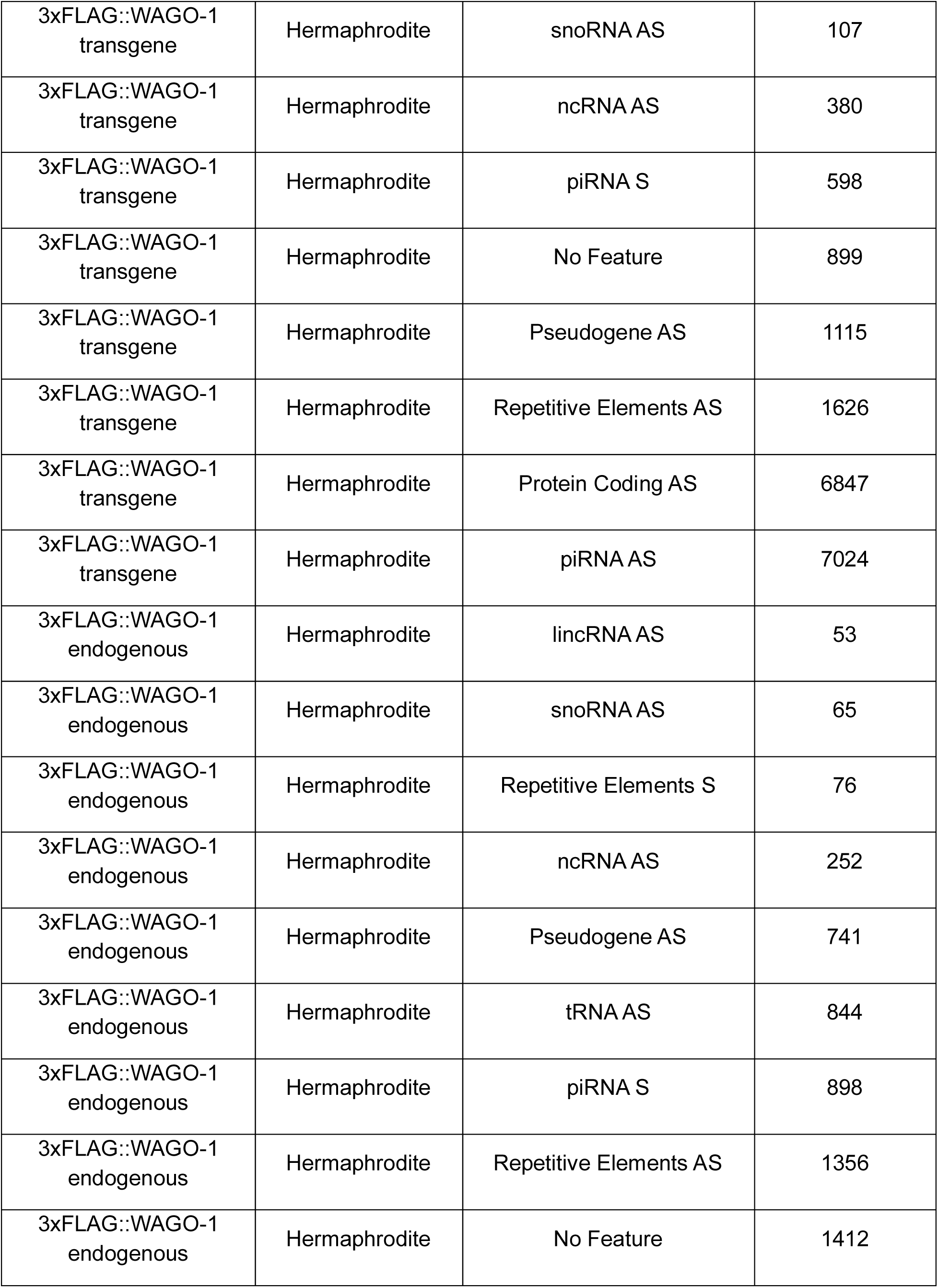
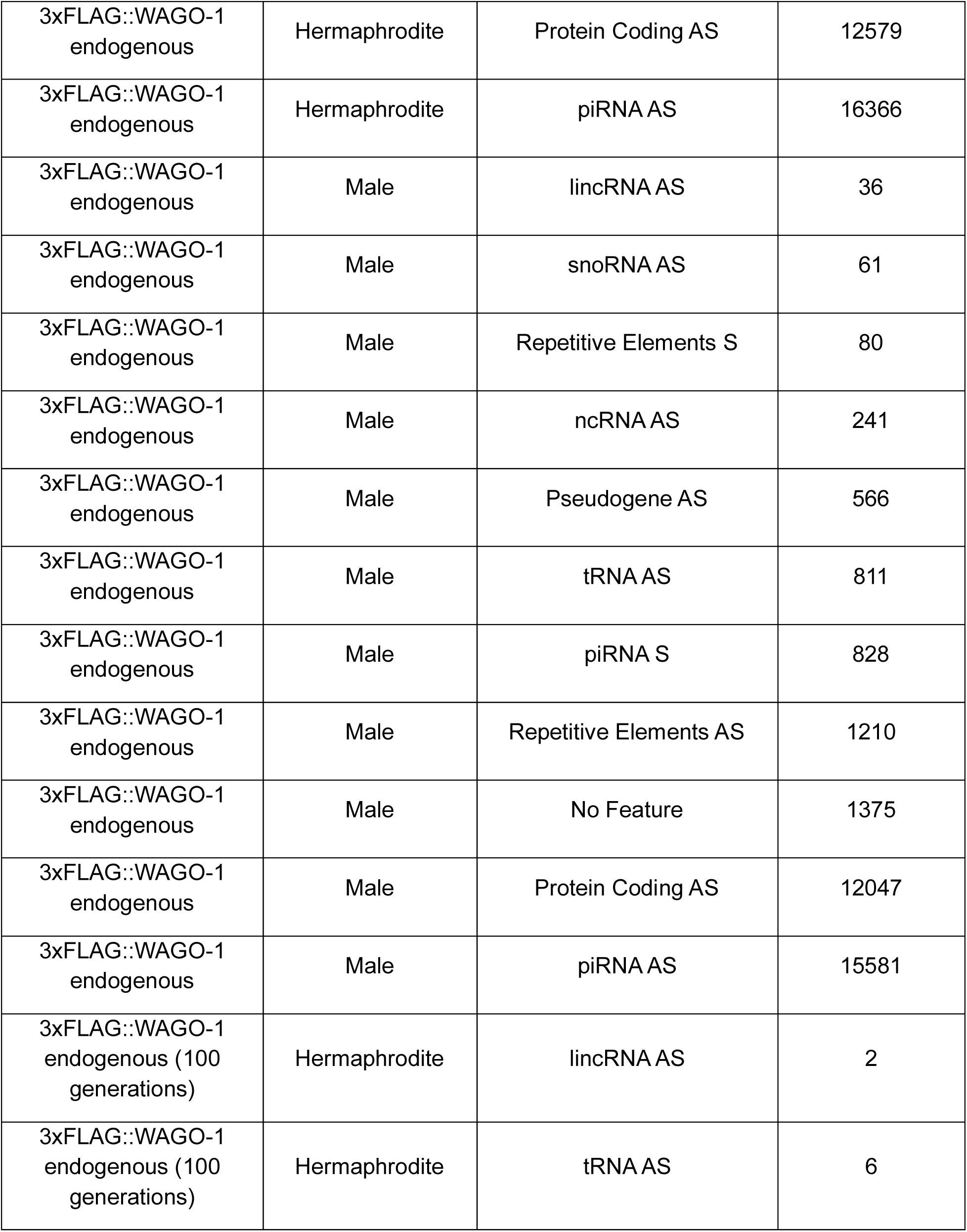
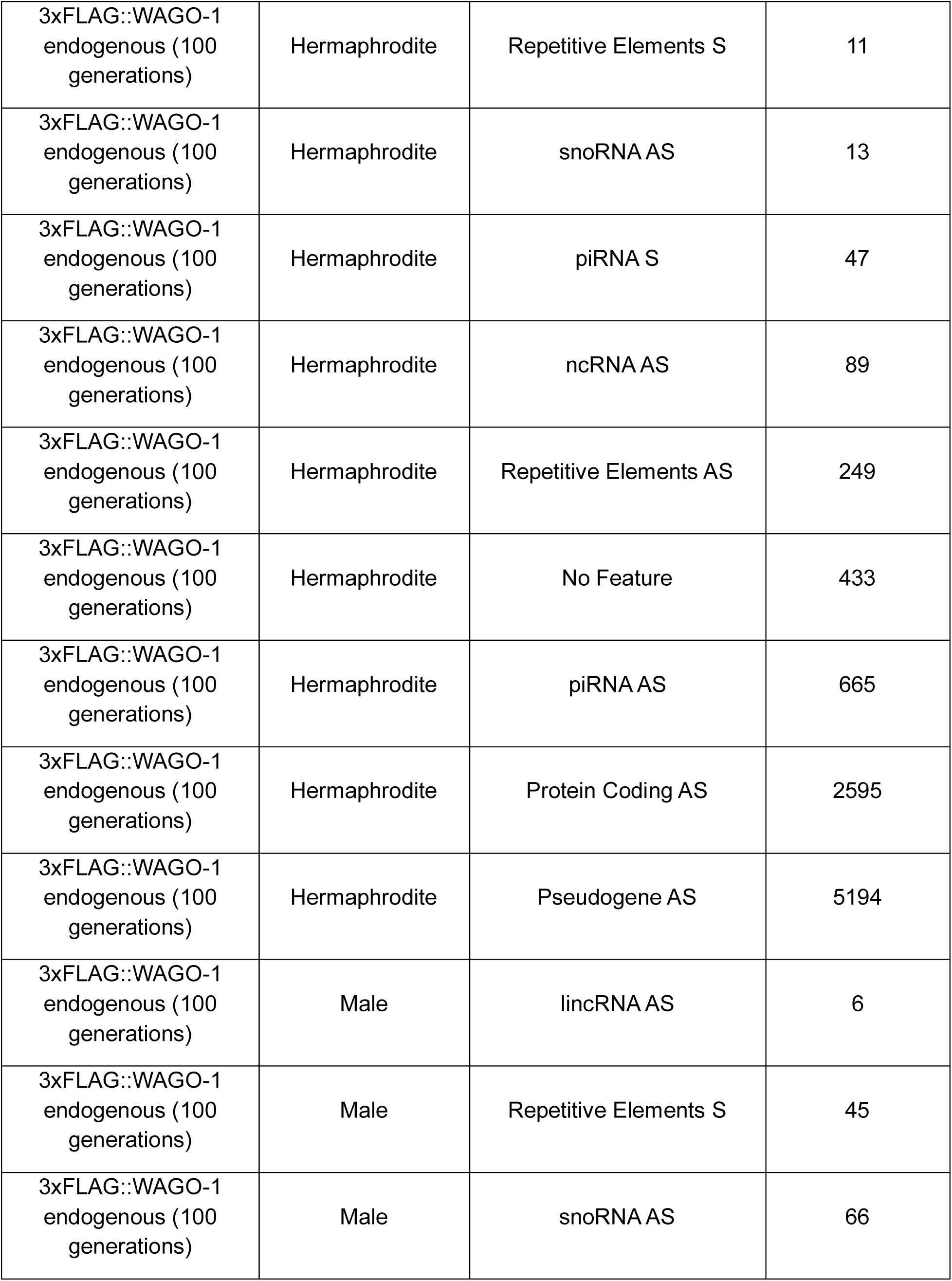
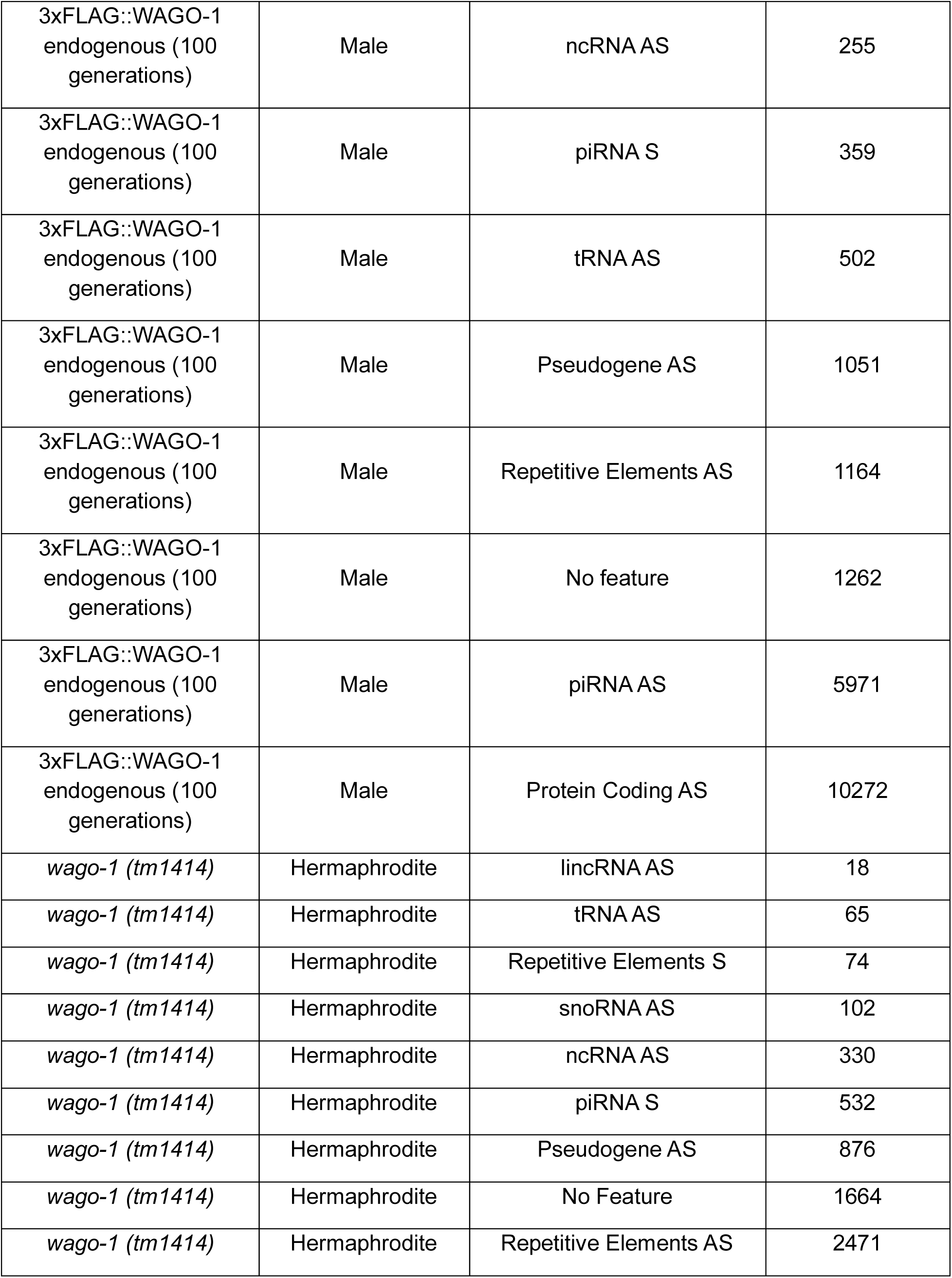
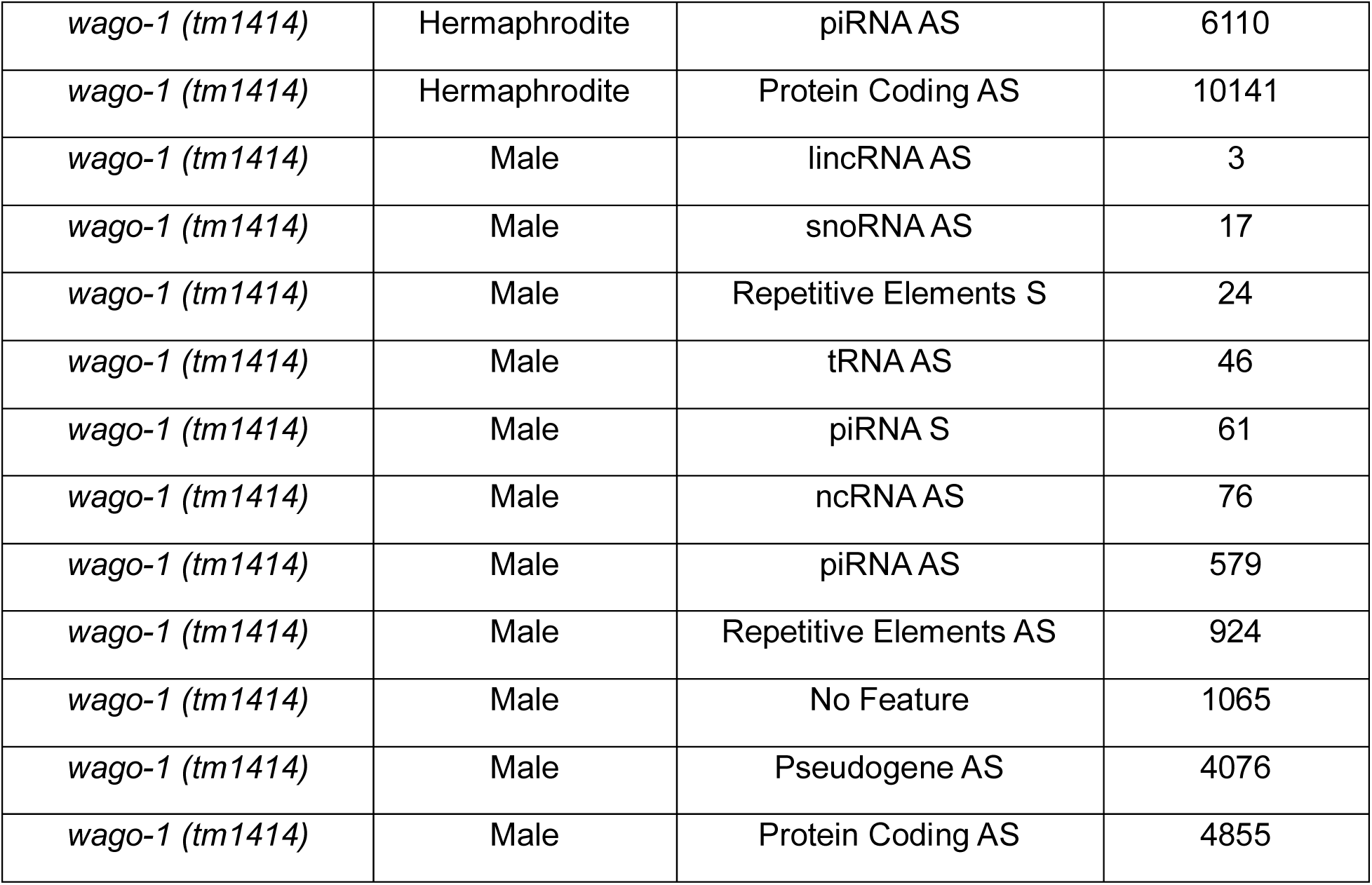
21U-RNA biotypes for wildtype and mutant WAGO-1 males and hermaphrodites.

| **Genotype** | **Sex** | **Biotype** | **Counts** |
| --- | --- | --- | --- |
| Wild-type | Hermaphrodite | lincRNA AS | 4 |
| Wild-type | Hermaphrodite | tRNA AS | 5 |
| Wild-type | Hermaphrodite | Repetitive Elements S | 25 |
| Wild-type | Hermaphrodite | snoRNA AS | 33 |
| Wild-type | Hermaphrodite | piRNA S | 94 |
| Wild-type | Hermaphrodite | ncRNA AS | 133 |
| Wild-type | Hermaphrodite | Repetitive Elements AS | 374 |
| Wild-type | Hermaphrodite | No Feature | 378 |
| Wild-type | Hermaphrodite | piRNA AS | 1112 |
| Wild-type | Hermaphrodite | Protein CodingAS | 2841 |
| Wild-type | Hermaphrodite | Pseudogene AS | 5507 |
| Wild-type | Male | lincRNA AS | 18 |
| Wild-type | Male | snoRNA AS | 42 |
| Wild-type | Male | Repetitive Elements S | 49 |
| Wild-type | Male | ncRNA AS | 207 |
| Wild-type | Male | piRNA S | 266 |
| Wild-type | Male | tRNA AS | 459 |
| Wild-type | Male | Pseudogene AS | 829 |
| Wild-type | Male | No Feature | 1471 |
| Wild-type | Male | Repetitive Elements AS | 1493 |
| Wild-type | Male | piRNA AS | 5032 |
| Wild-type | Male | Protein Coding AS | 12060 |
| 3xFLAG::WAGO-1<br>transgene | Hermaphrodite | lincRNA AS | 43 |
| 3xFLAG::WAGO-1<br>transgene | Hermaphrodite | tRNA AS | 55 |
| 3xFLAG::WAGO-1<br>transgene | Hermaphrodite | Repetitive Elements S | 88 |
| 3xFLAG::WAGO-1<br>transgene | Hermaphrodite | snoRNA AS | 107 |
| 3xFLAG::WAGO-1<br>transgene | Hermaphrodite | ncRNA AS | 380 |
| 3xFLAG::WAGO-1<br>transgene | Hermaphrodite | piRNA S | 598 |
| 3xFLAG::WAGO-1<br>transgene | Hermaphrodite | No Feature | 899 |
| 3xFLAG::WAGO-1<br>transgene | Hermaphrodite | Pseudogene AS | 1115 |
| 3xFLAG::WAGO-1<br>transgene | Hermaphrodite | Repetitive Elements AS | 1626 |
| 3xFLAG::WAGO-1<br>transgene | Hermaphrodite | Protein Coding AS | 6847 |
| 3xFLAG::WAGO-1<br>transgene | Hermaphrodite | piRNA AS | 7024 |
| 3xFLAG::WAGO-1<br>endogenous | Hermaphrodite | lincRNA AS | 53 |
| 3xFLAG::WAGO-1<br>endogenous | Hermaphrodite | snoRNA AS | 65 |
| 3xFLAG::WAGO-1<br>endogenous | Hermaphrodite | Repetitive Elements S | 76 |
| 3xFLAG::WAGO-1<br>endogenous | Hermaphrodite | ncRNA AS | 252 |
| 3xFLAG::WAGO-1<br>endogenous | Hermaphrodite | Pseudogene AS | 741 |
| 3xFLAG::WAGO-1<br>endogenous | Hermaphrodite | tRNA AS | 844 |
| 3xFLAG::WAGO-1<br>endogenous | Hermaphrodite | piRNA S | 898 |
| 3xFLAG::WAGO-1<br>endogenous | Hermaphrodite | Repetitive Elements AS | 1356 |
| 3xFLAG::WAGO-1<br>endogenous | Hermaphrodite | No Feature | 1412 |
| 3xFLAG::WAGO-1<br>endogenous | Hermaphrodite | Protein Coding AS | 12579 |
| 3xFLAG::WAGO-1<br>endogenous | Hermaphrodite | piRNA AS | 16366 |
| 3xFLAG::WAGO-1<br>endogenous | Male | lincRNA AS | 36 |
| 3xFLAG::WAGO-1<br>endogenous | Male | snoRNA AS | 61 |
| 3xFLAG::WAGO-1<br>endogenous | Male | Repetitive Elements S | 80 |
| 3xFLAG::WAGO-1<br>endogenous | Male | ncRNA AS | 241 |
| 3xFLAG::WAGO-1<br>endogenous | Male | Pseudogene AS | 566 |
| 3xFLAG::WAGO-1<br>endogenous | Male | tRNA AS | 811 |
| 3xFLAG::WAGO-1<br>endogenous | Male | piRNA S | 828 |
| 3xFLAG::WAGO-1<br>endogenous | Male | Repetitive Elements AS | 1210 |
| 3xFLAG::WAGO-1<br>endogenous | Male | No Feature | 1375 |
| 3xFLAG::WAGO-1<br>endogenous | Male | Protein Coding AS | 12047 |
| 3xFLAG::WAGO-1<br>endogenous | Male | piRNA AS | 15581 |
| 3xFLAG::WAGO-1<br>endogenous (100<br>generations) | Hermaphrodite | lincRNA AS | 2 |
| 3xFLAG::WAGO-1<br>endogenous (100<br>generations) | Hermaphrodite | tRNA AS | 6 |
| 3xFLAG::WAGO-1<br>endogenous (100<br>generations) | Hermaphrodite | Repetitive Elements S | 11 |
| 3xFLAG::WAGO-1<br>endogenous (100<br>generations) | Hermaphrodite | snoRNA AS | 13 |
| 3xFLAG::WAGO-1<br>endogenous (100<br>generations) | Hermaphrodite | piRNA S | 47 |
| 3xFLAG::WAGO-1<br>endogenous (100<br>generations) | Hermaphrodite | ncRNA AS | 89 |
| 3xFLAG::WAGO-1<br>endogenous (100<br>generations) | Hermaphrodite | Repetitive Elements AS | 249 |
| 3xFLAG::WAGO-1<br>endogenous (100<br>generations) | Hermaphrodite | No Feature | 433 |
| 3xFLAG::WAGO-1<br>endogenous (100<br>generations) | Hermaphrodite | piRNA AS | 665 |
| 3xFLAG::WAGO-1<br>endogenous (100<br>generations) | Hermaphrodite | Protein Coding AS | 2595 |
| 3xFLAG::WAGO-1<br>endogenous (100<br>generations) | Hermaphrodite | Pseudogene AS | 5194 |
| 3xFLAG::WAGO-1<br>endogenous (100<br>generations) | Male | lincRNA AS | 6 |
| 3xFLAG::WAGO-1<br>endogenous (100<br>generations) | Male | Repetitive Elements S | 45 |
| 3xFLAG::WAGO-1<br>endogenous (100<br>generations) | Male | snoRNA AS | 66 |
| 3xFLAG::WAGO-1<br>endogenous (100<br>generations) | Male | ncRNA AS | 255 |
| 3xFLAG::WAGO-1<br>endogenous (100<br>generations) | Male | piRNA S | 359 |
| 3xFLAG::WAGO-1<br>endogenous (100<br>generations) | Male | tRNA AS | 502 |
| 3xFLAG::WAGO-1<br>endogenous (100<br>generations) | Male | Pseudogene AS | 1051 |
| 3xFLAG::WAGO-1<br>endogenous (100<br>generations) | Male | Repetitive Elements AS | 1164 |
| 3xFLAG::WAGO-1<br>endogenous (100<br>generations) | Male | No feature | 1262 |
| 3xFLAG::WAGO-1<br>endogenous (100<br>generations) | Male | piRNA AS | 5971 |
| 3xFLAG::WAGO-1<br>endogenous (100<br>generations) | Male | Protein Coding AS | 10272 |
| *wago-1 (tm1414)* | Hermaphrodite | lincRNA AS | 18 |
| *wago-1 (tm1414)* | Hermaphrodite | tRNA AS | 65 |
| *wago-1 (tm1414)* | Hermaphrodite | Repetitive Elements S | 74 |
| *wago-1 (tm1414)* | Hermaphrodite | snoRNA AS | 102 |
| *wago-1 (tm1414)* | Hermaphrodite | ncRNA AS | 330 |
| *wago-1 (tm1414)* | Hermaphrodite | piRNA S | 532 |
| *wago-1 (tm1414)* | Hermaphrodite | Pseudogene AS | 876 |
| *wago-1 (tm1414)* | Hermaphrodite | No Feature | 1664 |
| *wago-1 (tm1414)* | Hermaphrodite | Repetitive Elements AS | 2471 |
| *wago-1 (tm1414)* | Hermaphrodite | piRNA AS | 6110 |
| *wago-1 (tm1414)* | Hermaphrodite | Protein Coding AS | 10141 |
| *wago-1 (tm1414)* | Male | lincRNA AS | 3 |
| *wago-1 (tm1414)* | Male | snoRNA AS | 17 |
| *wago-1 (tm1414)* | Male | Repetitive Elements S | 24 |
| *wago-1 (tm1414)* | Male | tRNA AS | 46 |
| *wago-1 (tm1414)* | Male | piRNA S | 61 |
| *wago-1 (tm1414)* | Male | ncRNA AS | 76 |
| *wago-1 (tm1414)* | Male | piRNA AS | 579 |
| *wago-1 (tm1414)* | Male | Repetitive Elements AS | 924 |
| *wago-1 (tm1414)* | Male | No Feature | 1065 |
| *wago-1 (tm1414)* | Male | Pseudogene AS | 4076 |
| *wago-1 (tm1414)* | Male | Protein Coding AS | 4855 |

**Table S15:** 22G small RNA biotypes for wildtype and mutant WAGO-1 males and hermaphrodites.

| **Genotype** | **Sex** | **Biotype** | **Counts** |
| --- | --- | --- | --- |
| Wild-type | Hermaphrodite | tRNA AS | 1 |
| Wild-type | Hermaphrodite | lincRNA AS | 3 |
| Wild-type | Hermaphrodite | piRNA S | 4 |
| Wild-type | Hermaphrodite | piRNA AS | 8 |
| Wild-type | Hermaphrodite | snoRNA AS | 10 |
| Wild-type | Hermaphrodite | Repetitive Elements S | 38 |
| Wild-type | Hermaphrodite | ncRNA AS | 125 |
| Wild-type | Hermaphrodite | Repetitive Elements AS | 223 |
| Wild-type | Hermaphrodite | No feature | 333 |
| Wild-type | Hermaphrodite | Protein Coding AS | 2798 |
| Wild-type | Hermaphrodite | Pseudogene AS | 3195 |
| Wild-type | Male | lincRNA AS | 6 |
| Wild-type | Male | snoRNA AS | 20 |
| Wild-type | Male | Repetitive Elements S | 65 |
| Wild-type | Male | piRNA AS | 70 |
| Wild-type | Male | ncRNA AS | 78 |
| Wild-type | Male | tRNA AS | 98 |
| Wild-type | Male | piRNA S | 139 |
| Wild-type | Male | Pseudogene AS | 538 |
| Wild-type | Male | Repetitive Elements AS | 1073 |
| Wild-type | Male | No Feature | 1167 |
| Wild-type | Male | Protein Coding AS | 9208 |
| 3xFLAG::WAGO-1<br>transgene | Hermaphrodite | tRNA AS | 9 |
| 3xFLAG::WAGO-1<br>transgene | Hermaphrodite | lincRNA AS | 19 |
| 3xFLAG::WAGO-1<br>transgene | Hermaphrodite | snoRNA AS | 28 |
| 3xFLAG::WAGO-1<br>transgene | Hermaphrodite | piRNA S | 43 |
| 3xFLAG::WAGO-1<br>transgene | Hermaphrodite | piRNA AS | 65 |
| 3xFLAG::WAGO-1<br>transgene | Hermaphrodite | ncRNA AS | 123 |
| 3xFLAG::WAGO-1<br>transgene | Hermaphrodite | Repetitive Elements S | 134 |
| 3xFLAG::WAGO-1<br>transgene | Hermaphrodite | Pseudogene AS | 546 |
| 3xFLAG::WAGO-1<br>transgene | Hermaphrodite | No Feature | 571 |
| 3xFLAG::WAGO-1<br>transgene | Hermaphrodite | Repetitive Elements AS | 896 |
| 3xFLAG::WAGO-1<br>transgene | Hermaphrodite | Protein Coding AS | 5378 |
| 3xFLAG::WAGO-1<br>endogenous | Hermaphrodite | lincRNA AS | 20 |
| 3xFLAG::WAGO-1<br>endogenous | Hermaphrodite | snoRNA AS | 30 |
| 3xFLAG::WAGO-1<br>endogenous | Hermaphrodite | tRNA_AS | 74 |
| 3xFLAG::WAGO-1<br>endogenous | Hermaphrodite | Repetitive Elements S | 80 |
| 3xFLAG::WAGO-1<br>endogenous | Hermaphrodite | piRNA S | 91 |
| 3xFLAG::WAGO-1<br>endogenous | Hermaphrodite | piRNA AS | 96 |
| 3xFLAG::WAGO-1<br>endogenous | Hermaphrodite | ncRNA AS | 186 |
| 3xFLAG::WAGO-1<br>endogenous | Hermaphrodite | Pseudogene AS | 348 |
| 3xFLAG::WAGO-1<br>endogenous | Hermaphrodite | Repetitive Elements AS | 892 |
| 3xFLAG::WAGO-1<br>endogenous | Hermaphrodite | No Feature | 1189 |
| 3xFLAG::WAGO-1<br>endogenous | Hermaphrodite | Protein Coding AS | 13201 |
| 3xFLAG::WAGO-1<br>endogenous | Male | lincRNA AS | 18 |
| 3xFLAG::WAGO-1<br>endogenous | Male | snoRNA AS | 22 |
| 3xFLAG::WAGO-1<br>endogenous | Male | tRNA AS | 67 |
| 3xFLAG::WAGO-1<br>endogenous | Male | Repetitive Elements S | 77 |
| 3xFLAG::WAGO-1<br>endogenous | Male | piRNA S | 88 |
| 3xFLAG::WAGO-1<br>endogenous | Male | piRNA AS | 96 |
| 3xFLAG::WAGO-1<br>endogenous | Male | ncRNA AS | 171 |
| 3xFLAG::WAGO-1<br>endogenous | Male | Pseudogene AS | 251 |
| 3xFLAG::WAGO-1<br>endogenous | Male | Repetitive Elements AS | 874 |
| 3xFLAG::WAGO-1<br>endogenous | Male | No Feature | 1189 |
| 3xFLAG::WAGO-1<br>endogenous | Male | Protein Coding AS | 12491 |
| 3xFLAG::WAGO-1<br>endogenous (100<br>generations) | Hermaphrodite | tRNA AS | 1 |
| 3xFLAG::WAGO-1<br>endogenous (100<br>generations) | Hermaphrodite | lincRNA AS | 2 |
| 3xFLAG::WAGO-1<br>endogenous (100<br>generations) | Hermaphrodite | snoRNA AS | 4 |
| 3xFLAG::WAGO-1<br>endogenous (100<br>generations) | Hermaphrodite | piRNA S | 6 |
| 3xFLAG::WAGO-1<br>endogenous (100<br>generations) | Hermaphrodite | piRNA AS | 9 |
| 3xFLAG::WAGO-1<br>endogenous (100<br>generations) | Hermaphrodite | Repetitive Elements S | 15 |
| 3xFLAG::WAGO-1<br>endogenous (100<br>generations) | Hermaphrodite | ncRNA AS | 81 |
| 3xFLAG::WAGO-1<br>endogenous (100<br>generations) | Hermaphrodite | Repetitive Elements AS | 195 |
| 3xFLAG::WAGO-1<br>endogenous (100<br>generations) | Hermaphrodite | No Feature | 397 |
| 3xFLAG::WAGO-1<br>endogenous (100<br>generations) | Hermaphrodite | Protein Coding AS | 2670 |
| 3xFLAG::WAGO-1<br>endogenous (100<br>generations) | Hermaphrodite | Pseudogene AS | 2913 |
| 3xFLAG::WAGO-1<br>endogenous (100<br>generations) | Male | lincRNA AS | 8 |
| 3xFLAG::WAGO-1<br>endogenous (100<br>generations) | Male | snoRNA AS | 20 |
| 3xFLAG::WAGO-1<br>endogenous (100<br>generations) | Male | piRNA AS | 76 |
| 3xFLAG::WAGO-1<br>endogenous (100<br>generations) | Male | Repetitive Elements S | 79 |
| 3xFLAG::WAGO-1<br>endogenous (100<br>generations) | Male | tRNA AS | 89 |
| 3xFLAG::WAGO-1<br>endogenous (100<br>generations) | Male | piRNA S | 112 |
| 3xFLAG::WAGO-1<br>endogenous (100<br>generations) | Male | ncRNA AS | 119 |
| 3xFLAG::WAGO-1<br>endogenous (100<br>generations) | Male | Pseudogene AS | 728 |
| 3xFLAG::WAGO-1<br>endogenous (100<br>generations) | Male | No Feature | 1303 |
| 3xFLAG::WAGO-1<br>endogenous (100<br>generations) | Male | Repetitive Elements AS | 1520 |
| 3xFLAG::WAGO-1<br>endogenous (100<br>generations) | Male | Protein Coding AS | 10192 |
| *wago-1 (tm1414)* | Hermaphrodite | tRNA AS | 14 |
| *wago-1 (tm1414)* | Hermaphrodite | lincRNA AS | 16 |
| *wago-1 (tm1414)* | Hermaphrodite | snoRNA AS | 42 |
| *wago-1 (tm1414)* | Hermaphrodite | piRNA AS | 64 |
| *wago-1 (tm1414)* | Hermaphrodite | Repetitive Elements S | 65 |
| *wago-1 (tm1414)* | Hermaphrodite | piRNA S | 65 |
| *wago-1 (tm1414)* | Hermaphrodite | ncRNA AS | 206 |
| *wago-1 (tm1414)* | Hermaphrodite | Pseudogene AS | 408 |
| *wago-1 (tm1414)* | Hermaphrodite | No Feature | 1064 |
| *wago-1 (tm1414)* | Hermaphrodite | Repetitive Elements AS | 1570 |
| *wago-1 (tm1414)* | Hermaphrodite | Protein Coding AS | 7525 |
| *wago-1 (tm1414)* | Male | lincRNA AS | 3 |
| *wago-1 (tm1414)* | Male | piRNA AS | 5 |
| *wago-1 (tm1414)* | Male | tRNA AS | 6 |
| *wago-1 (tm1414)* | Male | snoRNA AS | 7 |
| *wago-1 (tm1414)* | Male | piRNA S | 38 |
| *wago-1 (tm1414)* | Male | Repetitive Elements S | 40 |
| *wago-1 (tm1414)* | Male | ncRNA AS | 85 |
| *wago-1 (tm1414)* | Male | No Feature | 928 |
| *wago-1 (tm1414)* | Male | Repetitive Elements AS | 1016 |
| *wago-1 (tm1414)* | Male | Pseudogene AS | 2362 |
| *wago-1 (tm1414)* | Male | Protein Coding AS | 5471 |

**Table S16:** 26G small RNA biotypes for wildtype and mutant WAGO-1 males and hermaphrodites.

| Strain | Sex | Biotype | Count |
| --- | --- | --- | --- |
| Wild-type | Hermaphrodite | tRNA AS | 6 |
| Wild-type | Hermaphrodite | piRNA S | 6 |
| Wild-type | Hermaphrodite | piRNA AS | 8 |
| Wild-type | Hermaphrodite | Repetitive Elements S | 9 |
| Wild-type | Hermaphrodite | snoRNA AS | 9 |
| Wild-type | Hermaphrodite | ncRNA AS | 105 |
| Wild-type | Hermaphrodite | Repetitive Elements AS | 110 |
| Wild-type | Hermaphrodite | No Feature | 325 |
| Wild-type | Hermaphrodite | Protein Coding AS | 1722 |
| Wild-type | Hermaphrodite | Pseudogene AS | 1895 |
| Wild-type | Male | lincRNA AS | 2 |
| Wild-type | Male | snoRNA AS | 19 |
| Wild-type | Male | Repetitive Repeats | 51 |
| Wild-type | Male | piRNA AS | 99 |
| Wild-type | Male | tRNA AS | 132 |
| Wild-type | Male | Pseudogene AS | 362 |
| Wild-type | Male | piRNA S | 409 |
| Wild-type | Male | ncRNA AS | 936 |
| Wild-type | Male | Repetitive Elements AS | 1163 |
| Wild-type | Male | No Feature | 3617 |
| Wild-type | Male | Protein Coding AS | 13612 |
| 3xFLAG::WAGO-1 transgene | Hermaphrodite | lincRNA AS | 3 |
| 3xFLAG::WAGO-1 transgene | Hermaphrodite | Repetitive Elements S | 9 |
| 3xFLAG::WAGO-1 transgene | Hermaphrodite | piRNA S | 22 |
| 3xFLAG::WAGO-1 transgene | Hermaphrodite | tRNA AS | 30 |
| 3xFLAG::WAGO-1 transgene | Hermaphrodite | snoRNA AS | 47 |
| 3xFLAG::WAGO-1 transgene | Hermaphrodite | piRNA AS | 83 |
| 3xFLAG::WAGO-1 transgene | Hermaphrodite | ncRNA AS | 280 |
| 3xFLAG::WAGO-1 transgene | Hermaphrodite | Pseudogene AS | 301 |
| 3xFLAG::WAGO-1 transgene | Hermaphrodite | Repetitive Elements AS | 436 |
| 3xFLAG::WAGO-1 transgene | Hermaphrodite | No Feature | 764 |
| 3xFLAG::WAGO-1 transgene | Hermaphrodite | Protein Coding AS | 2259 |
| 3xFLAG::WAGO-1 endogenous | Hermaphrodite | lincRNA AS | 8 |
| 3xFLAG::WAGO-1 endogenous | Hermaphrodite | snoRNA AS | 32 |
| 3xFLAG::WAGO-1 endogenous | Hermaphrodite | tRNA AS | 81 |
| 3xFLAG::WAGO-1 endogenous | Hermaphrodite | Repetitive Elements | 119 |
| 3xFLAG::WAGO-1 endogenous | Hermaphrodite | piRNA AS | 178 |
| 3xFLAG::WAGO-1 endogenous | Hermaphrodite | Pseudogene AS | 181 |
| 3xFLAG::WAGO-1 endogenous | Hermaphrodite | piRNA | 1203 |
| 3xFLAG::WAGO-1 endogenous | Hermaphrodite | Repetitive Elements AS | 3864 |
| 3xFLAG::WAGO-1 endogenous | Hermaphrodite | ncRNA AS | 4992 |
| 3xFLAG::WAGO-1<br>endogenous | Hermaphrodite | No Feature | 14629 |
| 3xFLAG::WAGO-1<br>endogenous | Hermaphrodite | Protein Coding AS | 35084 |
| 3xFLAG::WAGO-1<br>endogenous | Male | lincRNA AS | 8 |
| 3xFLAG::WAGO-1<br>endogenous | Male | snoRNA AS | 33 |
| 3xFLAG::WAGO-1<br>endogenous | Male | tRNA AS | 90 |
| 3xFLAG::WAGO-1<br>endogenous | Male | Repetitive Elements S | 113 |
| 3xFLAG::WAGO-1<br>endogenous | Male | Pseudogene AS | 141 |
| 3xFLAG::WAGO-1<br>endogenous | Male | piRNA AS | 168 |
| 3xFLAG::WAGO-1<br>endogenous | Male | piRNA S | 1199 |
| 3xFLAG::WAGO-1<br>endogenous | Male | Repetitive Elements AS | 3822 |
| 3xFLAG::WAGO-1<br>endogenous | Male | ncRNA AS | 4885 |
| 3xFLAG::WAGO-1<br>endogenous | Male | No Feature | 14486 |
| 3xFLAG::WAGO-1<br>endogenous | Male | Protein Coding AS | 34412 |
| 3xFLAG::WAGO-1<br>endogenous (100<br>generations) | Hermaphrodite | lincRNA AS | 1 |
| 3xFLAG::WAGO-1<br>endogenous (100<br>generations) | Hermaphrodite | piRNA S | 2 |
| 3xFLAG::WAGO-1<br>endogenous (100<br>generations) | Hermaphrodite | tRNA AS | 3 |
| 3xFLAG::WAGO-1<br>endogenous (100<br>generations) | Hermaphrodite | Repetitive Elements S | 5 |
| 3xFLAG::WAGO-1<br>endogenous (100<br>generations) | Hermaphrodite | piRNA AS | 7 |
| 3xFLAG::WAGO-1<br>endogenous (100<br>generations) | Hermaphrodite | snoRNA AS | 8 |
| 3xFLAG::WAGO-1<br>endogenous (100<br>generations) | Hermaphrodite | ncRNA AS | 57 |
| 3xFLAG::WAGO-1<br>endogenous (100<br>generations) | Hermaphrodite | Repetitive Elements AS | 94 |
| 3xFLAG::WAGO-1<br>endogenous (100<br>generations) | Hermaphrodite | No Feature | 291 |
| 3xFLAG::WAGO-1<br>endogenous (100<br>generations) | Hermaphrodite | Pseudogene AS | 1596 |
| 3xFLAG::WAGO-1<br>endogenous (100<br>generations) | Hermaphrodite | Protein Coding AS | 1621 |
| 3xFLAG::WAGO-1<br>endogenous (100<br>generations) | Male | lincRNA AS | 9 |
| 3xFLAG::WAGO-1<br>endogenous (100<br>generations) | Male | snoRNA AS | 35 |
| 3xFLAG::WAGO-1<br>endogenous (100<br>generations) | Male | Repetitive Elements S | 45 |
| 3xFLAG::WAGO-1<br>endogenous (100<br>generations) | Male | piRNA AS | 86 |
| 3xFLAG::WAGO-1<br>endogenous (100<br>generations) | Male | tRNA AS | 151 |
| 3xFLAG::WAGO-1<br>endogenous (100<br>generations) | Male | piRNA S | 433 |
| 3xFLAG::WAGO-1<br>endogenous (100<br>generations) | Male | Pseudogene AS | 898 |
| 3xFLAG::WAGO-1<br>endogenous (100<br>generations) | Male | ncRNA AS | 1013 |
| 3xFLAG::WAGO-1<br>endogenous (100<br>generations) | Male | Repetitive Elements AS | 1275 |
| 3xFLAG::WAGO-1<br>endogenous (100<br>generations) | Male | No Feature | 3492 |
| 3xFLAG::WAGO-1<br>endogenous (100<br>generations) | Male | Protein Coding AS | 16172 |
| *wago-1 (tm1414)* | Hermaphrodite | lincRNA AS | 2 |
| *wago-1 (tm1414)* | Hermaphrodite | Repetitive Elements S | 18 |
| *wago-1 (tm1414)* | Hermaphrodite | tRNA AS | 25 |
| *wago-1 (tm1414)* | Hermaphrodite | piRNA S | 45 |
| *wago-1 (tm1414)* | Hermaphrodite | snoRNA AS | 50 |
| *wago-1 (tm1414)* | Hermaphrodite | piRNA AS | 61 |
| *wago-1 (tm1414)* | Hermaphrodite | ncRNA AS | 154 |
| *wago-1 (tm1414)* | Hermaphrodite | Pseudogene AS | 251 |
| *wago-1 (tm1414)* | Hermaphrodite | No Feature | 971 |
| *wago-1 (tm1414)* | Hermaphrodite | Repetitive Elements AS | 1056 |
| *wago-1 (tm1414)* | Hermaphrodite | Protein Coding AS | 3758 |
| *wago-1 (tm1414)* | Male | lincRNA AS | 2 |
| *wago-1 (tm1414)* | Male | snoRNA AS | 7 |
| *wago-1 (tm1414)* | Male | piRNA AS | 8 |
| *wago-1 (tm1414)* | Male | Repetitive Elements S | 12 |
| *wago-1 (tm1414)* | Male | tRNA AS | 16 |
| *wago-1 (tm1414)* | Male | piRNA S | 78 |
| *wago-1 (tm1414)* | Male | ncRNA AS | 107 |
| *wago-1 (tm1414)* | Male | No Feature | 897 |
| *wago-1 (tm1414)* | Male | Pseudogene AS | 1314 |
| *wago-1 (tm1414)* | Male | Repetitive Elements AS | 1994 |
| *wago-1 (tm1414)* | Male | Protein Coding AS | 3972 |

