## Supplementary figures and images for "WAGO-1 is a sexually dimorphic Argonaute protein required for proper germ granule structure and gametogenesis"

### Supplemental Figure S1

Figure S1

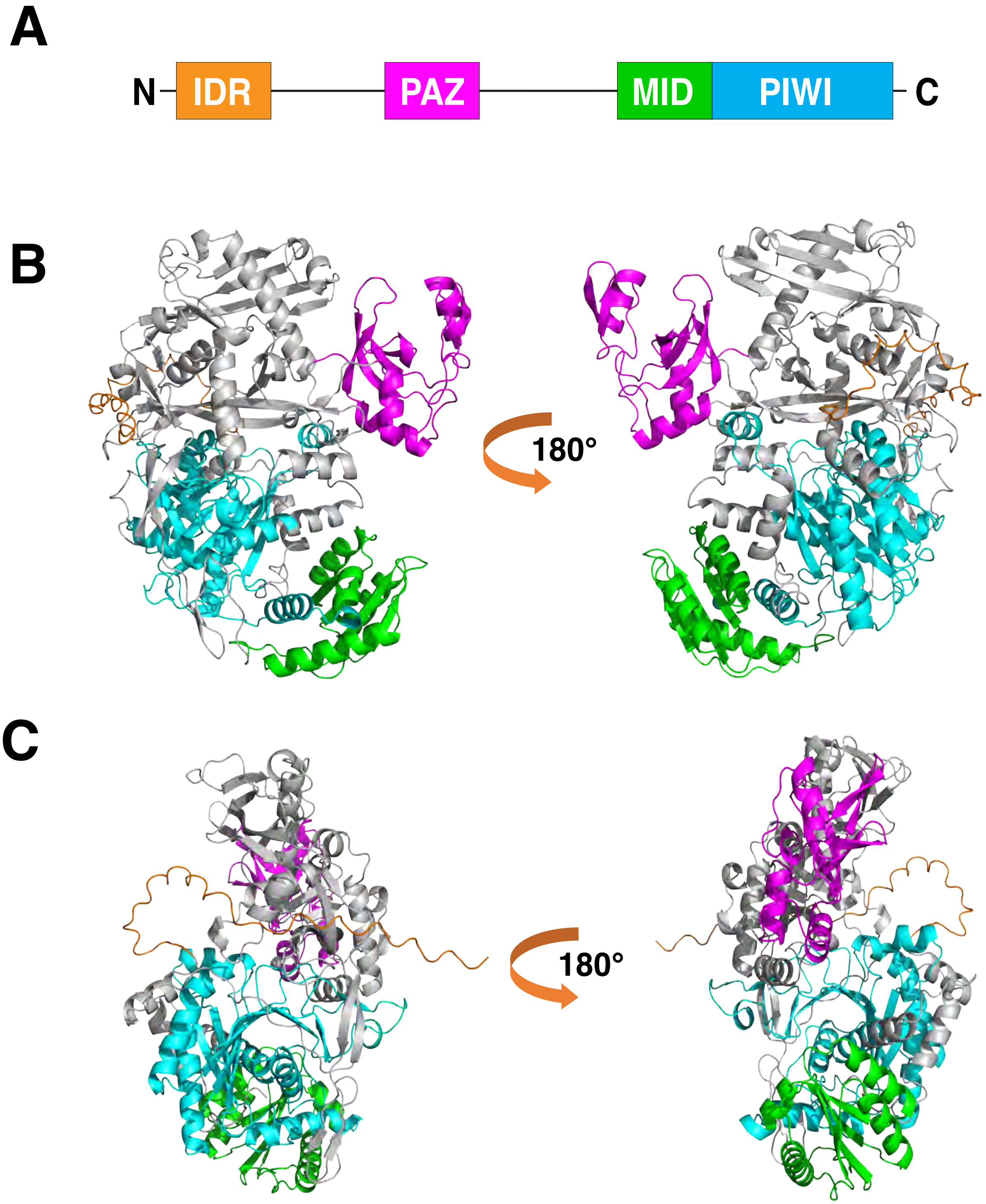

### Supplemental Figure S2

Figure S2

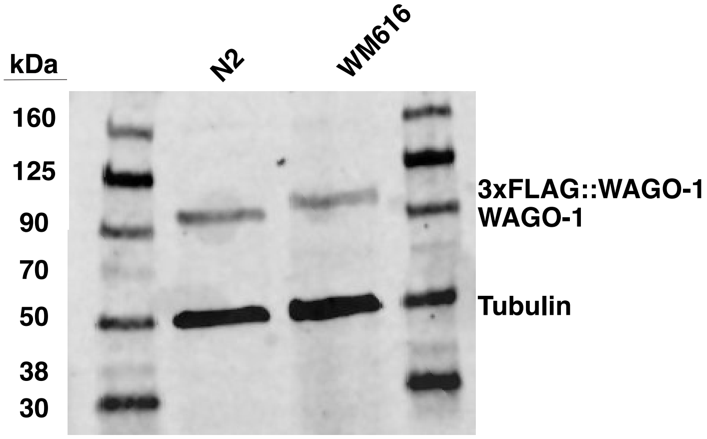

### Supplemental Figure S3

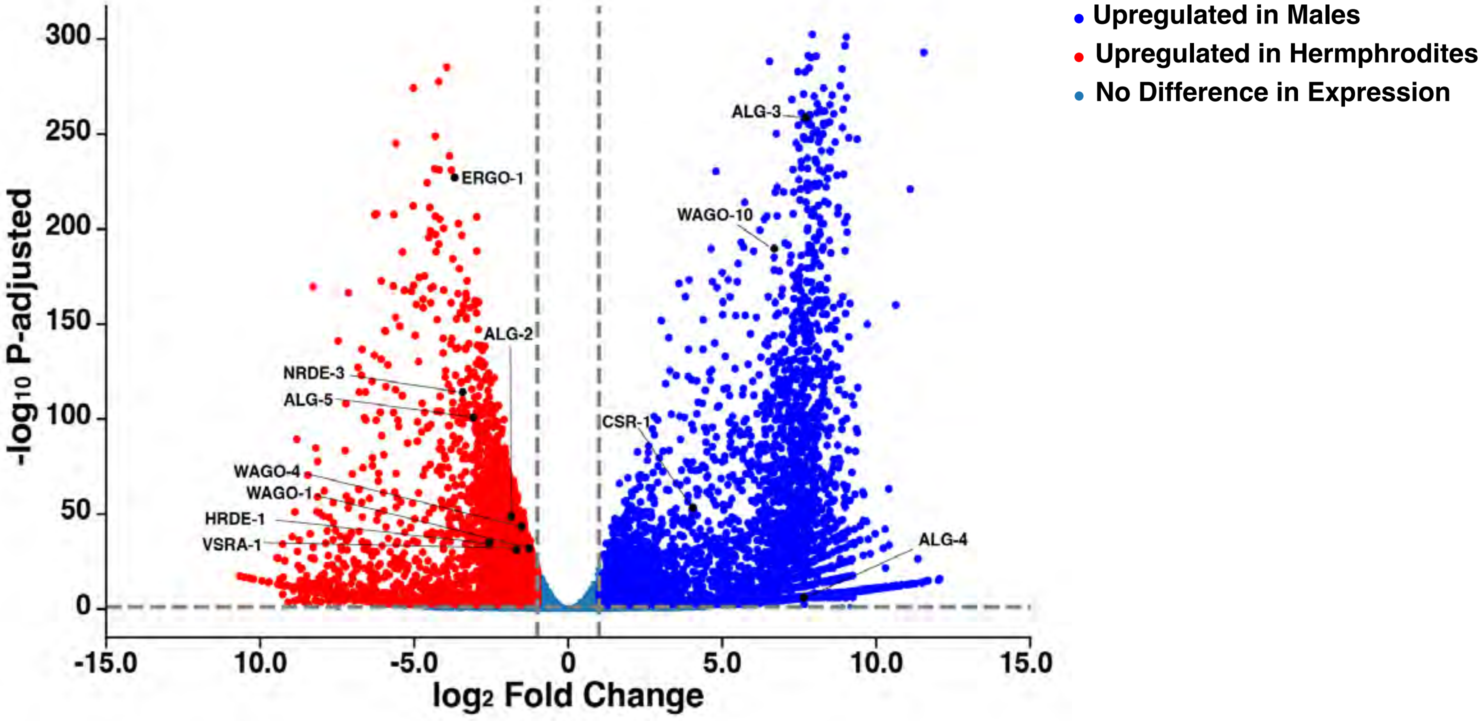

### Supplemental Figure S4

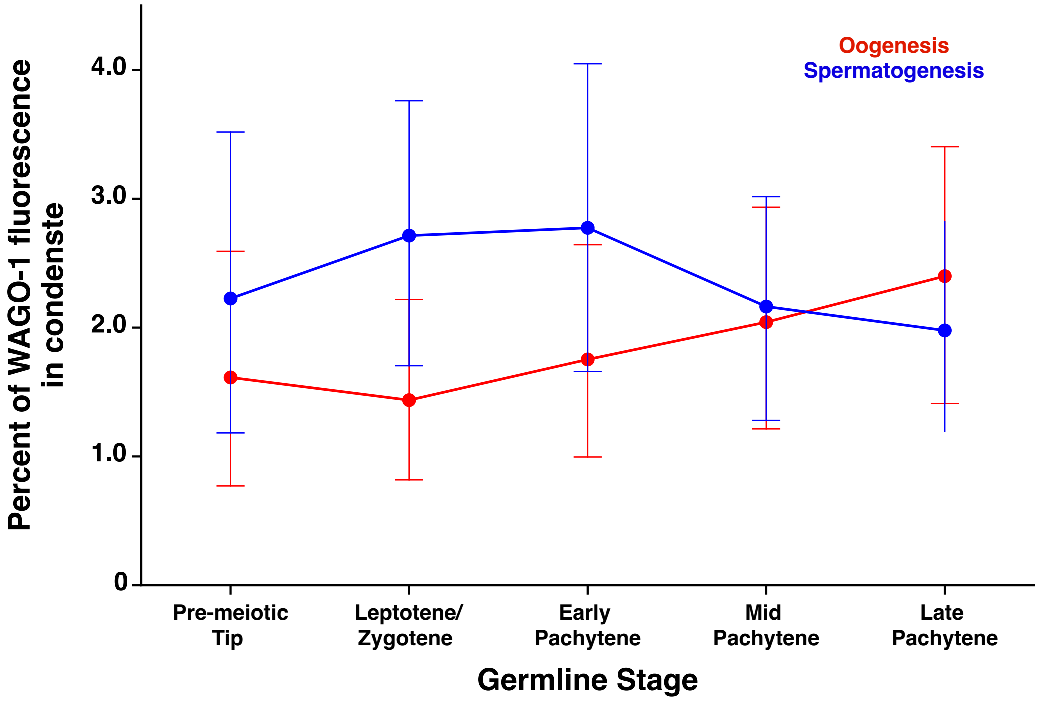

### Supplemental Figure S5

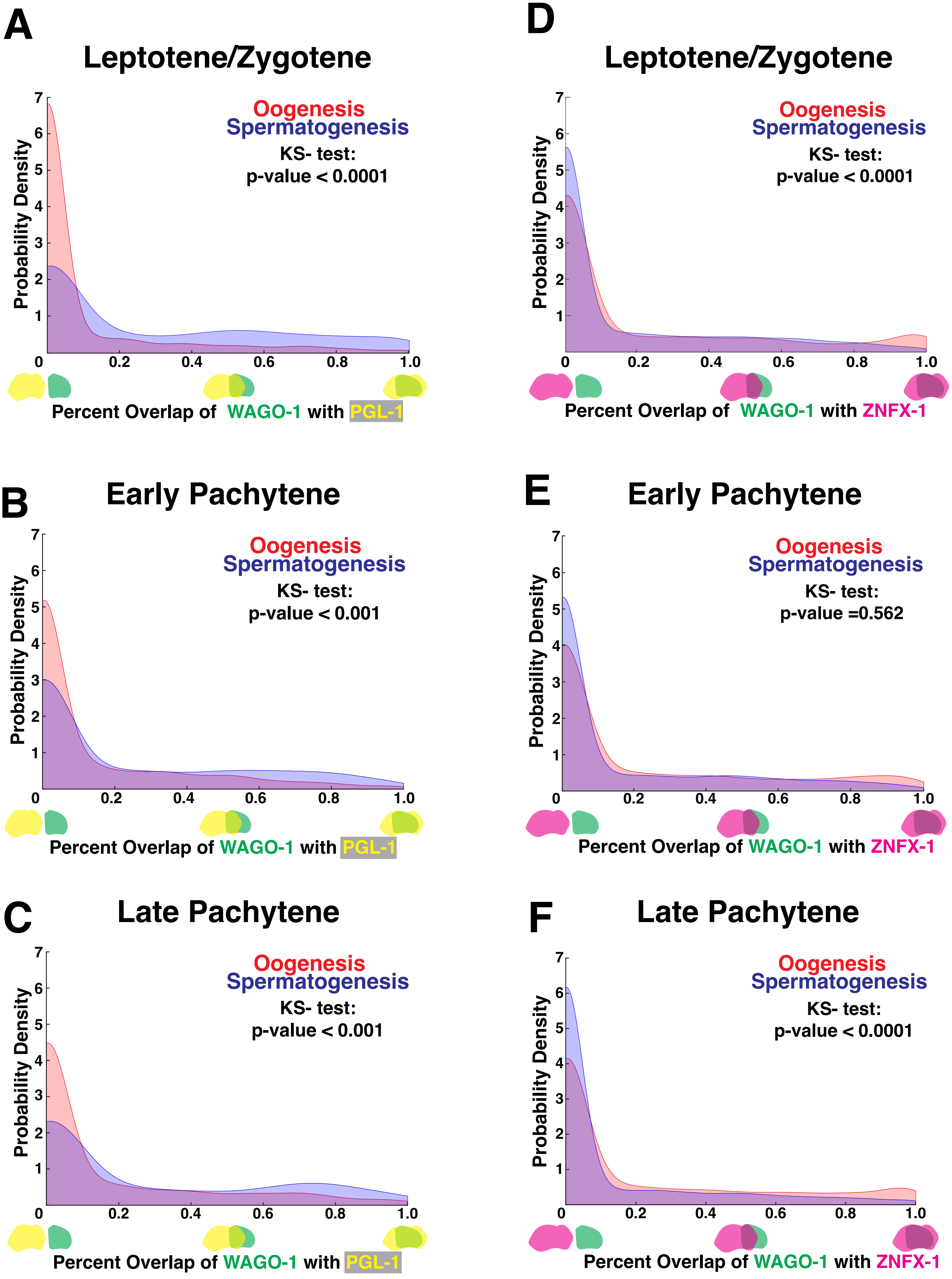

### Supplemental Figure S6

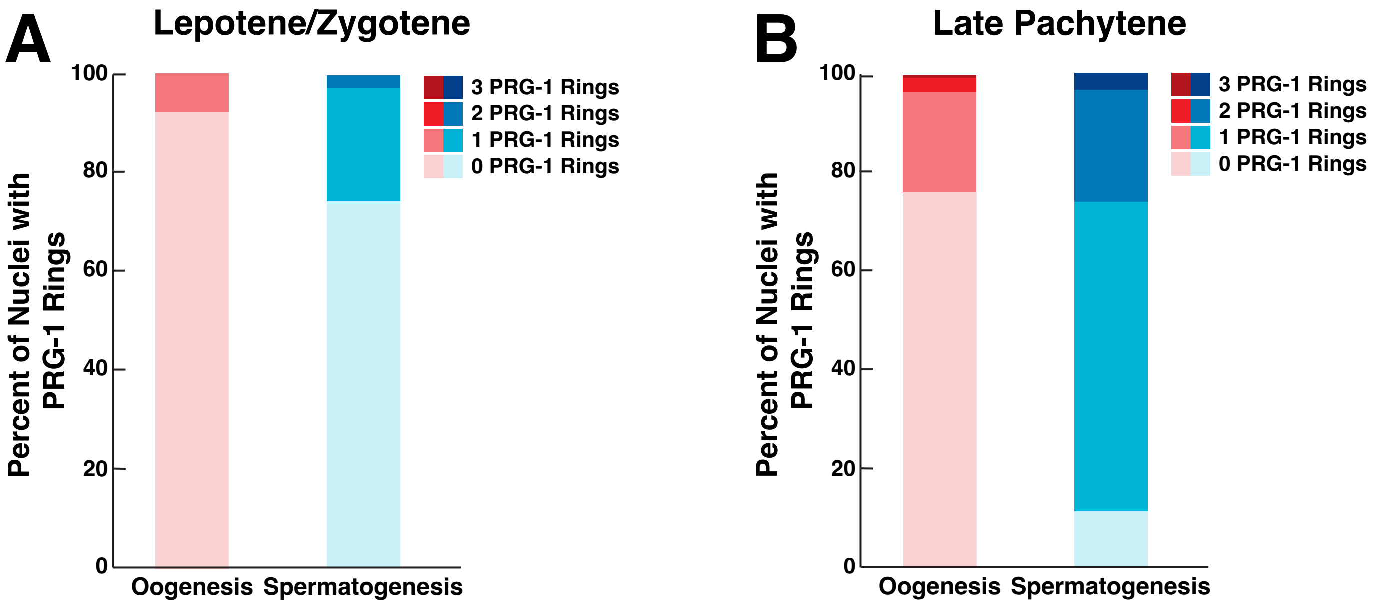

### Supplemental Figure S7

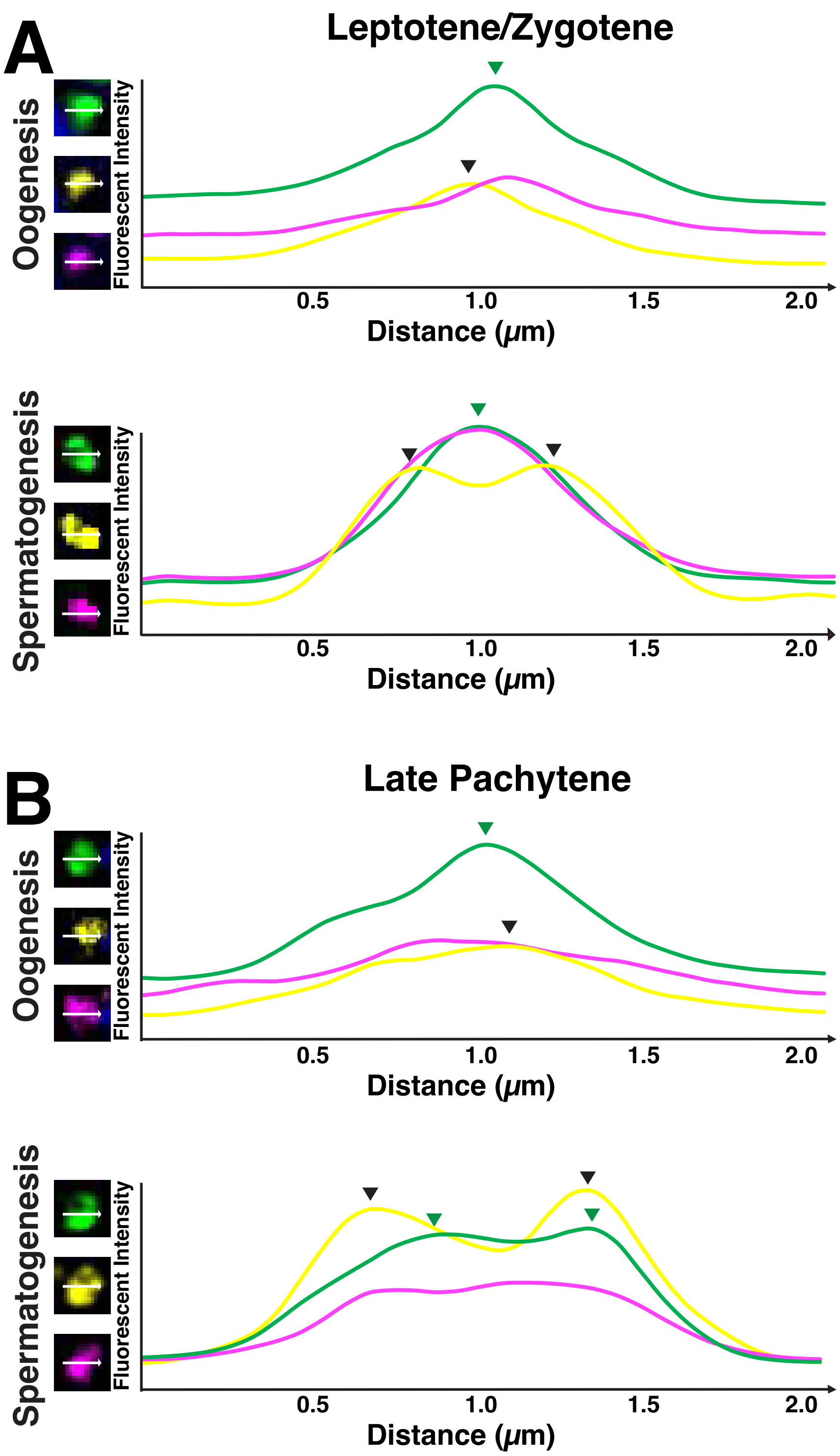

### Supplemental Figure S8

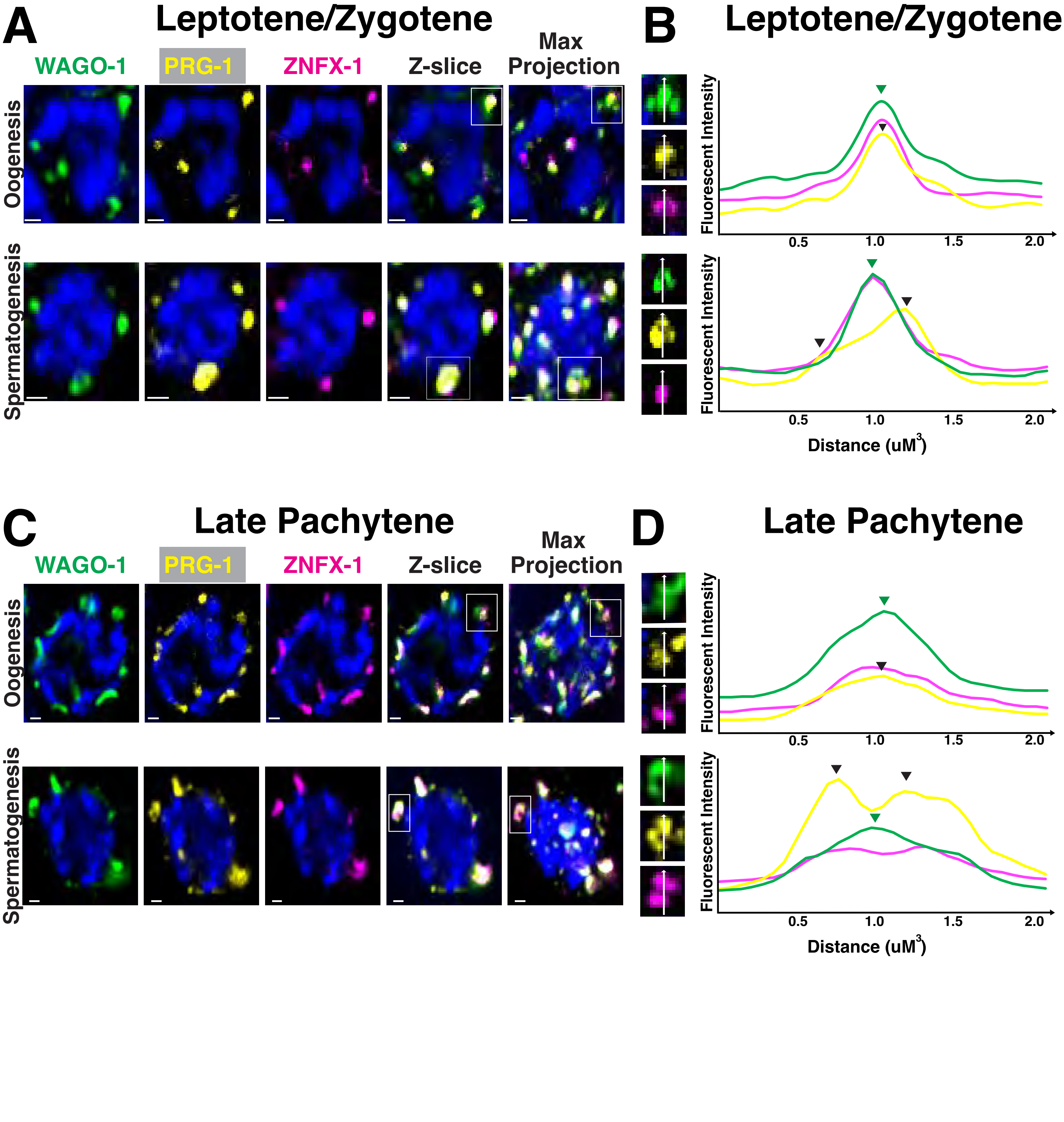

### Supplemental Figure S9

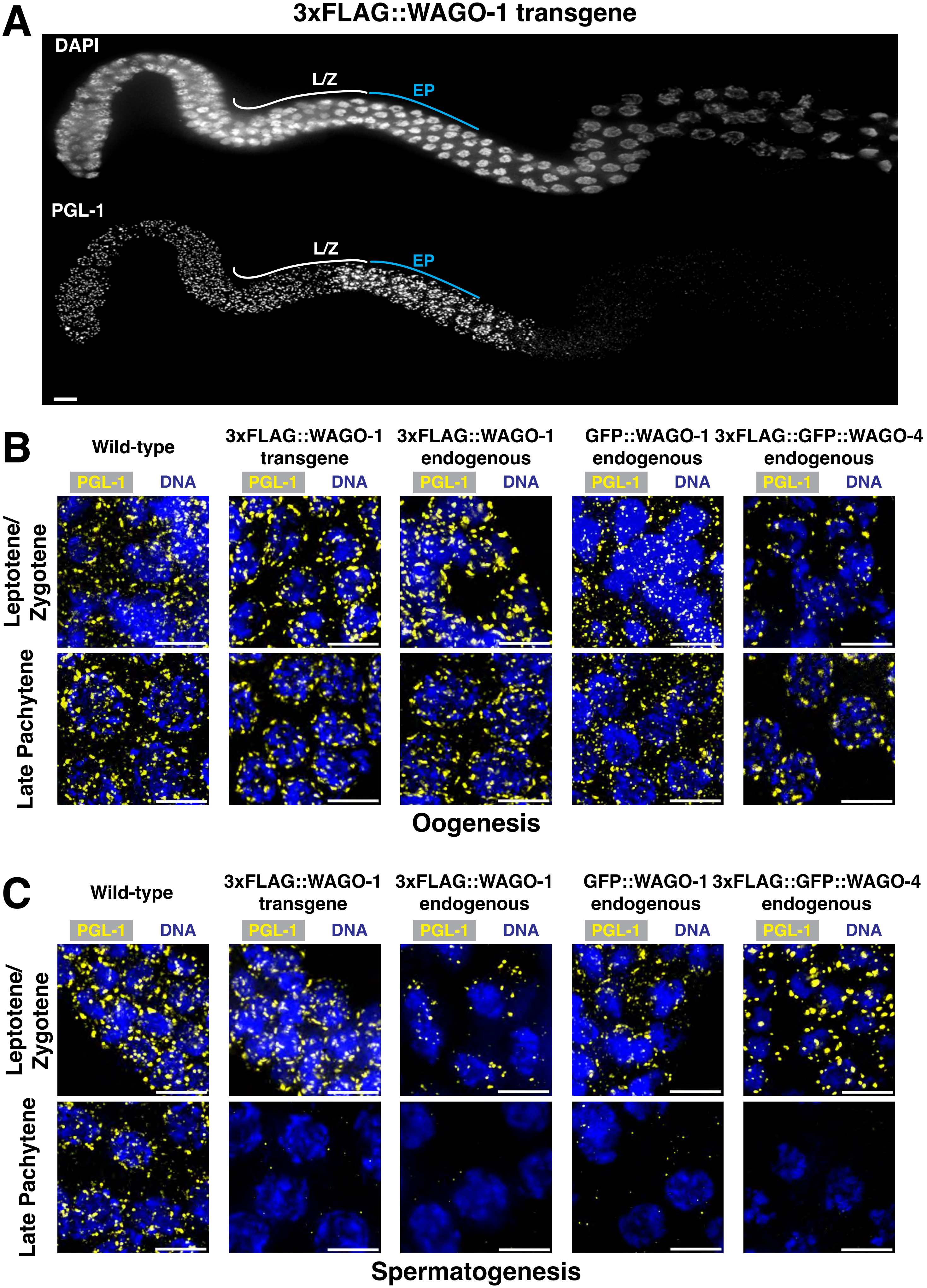

### Supplemental Figure S10

#### Supplemental Figure 10:

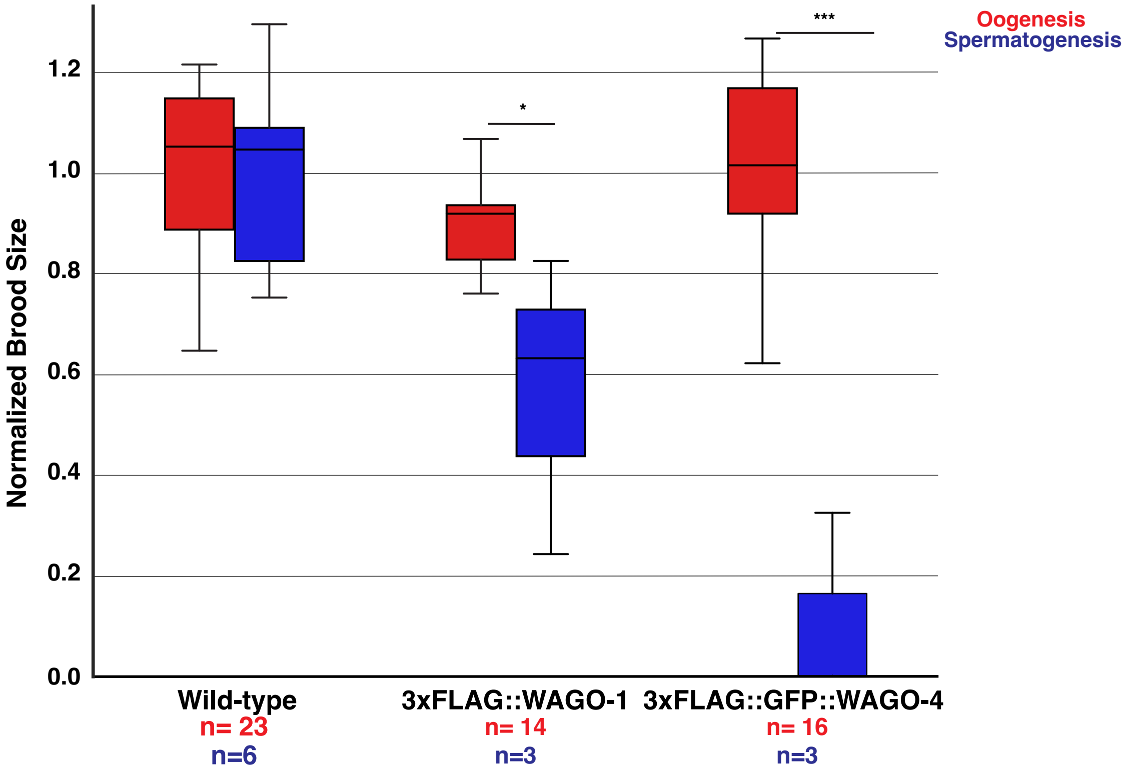

### Supplemental Figure S11

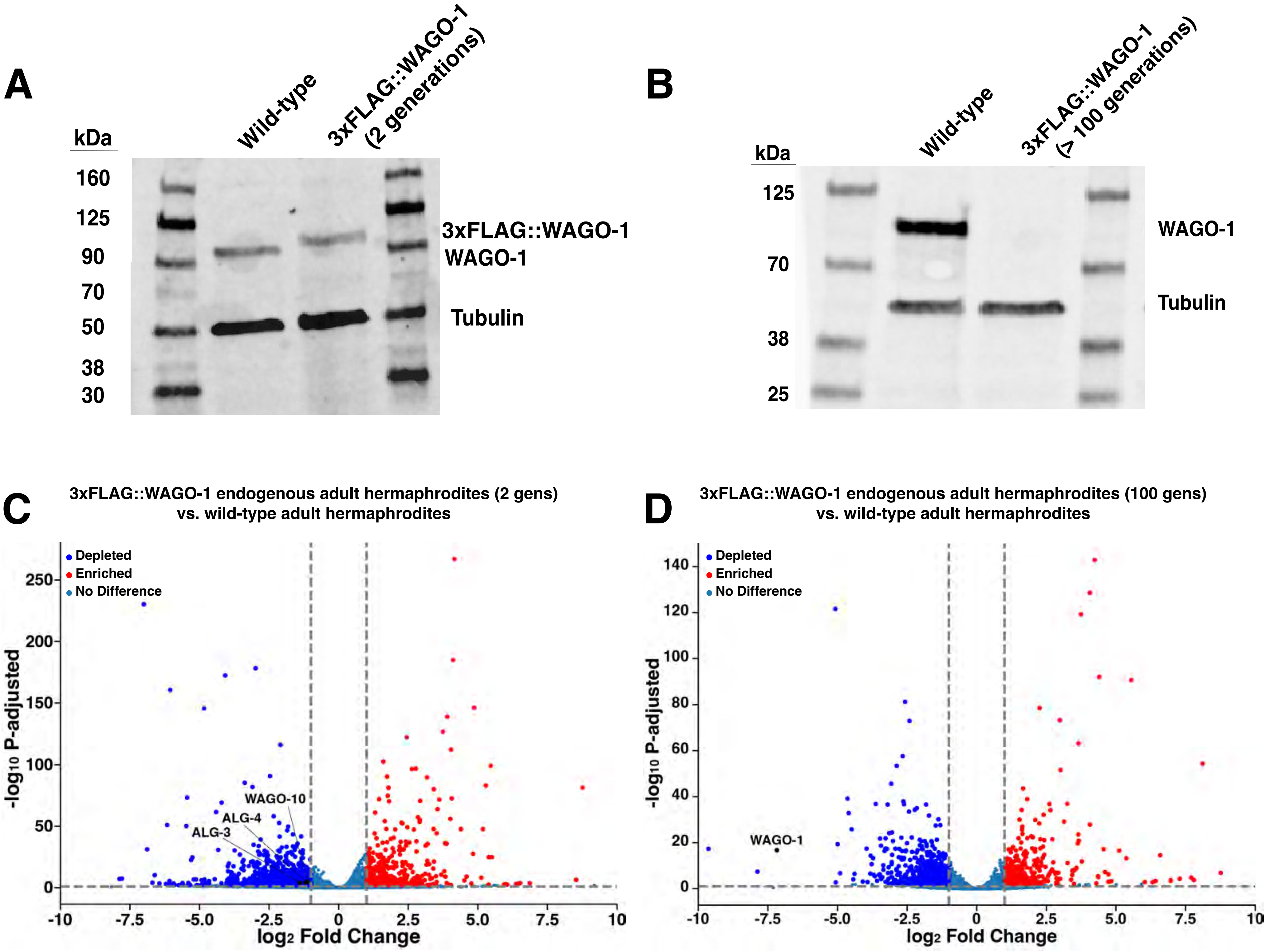

### Supplemental Figure S12

Figure S12

**A**

**Wild-type**

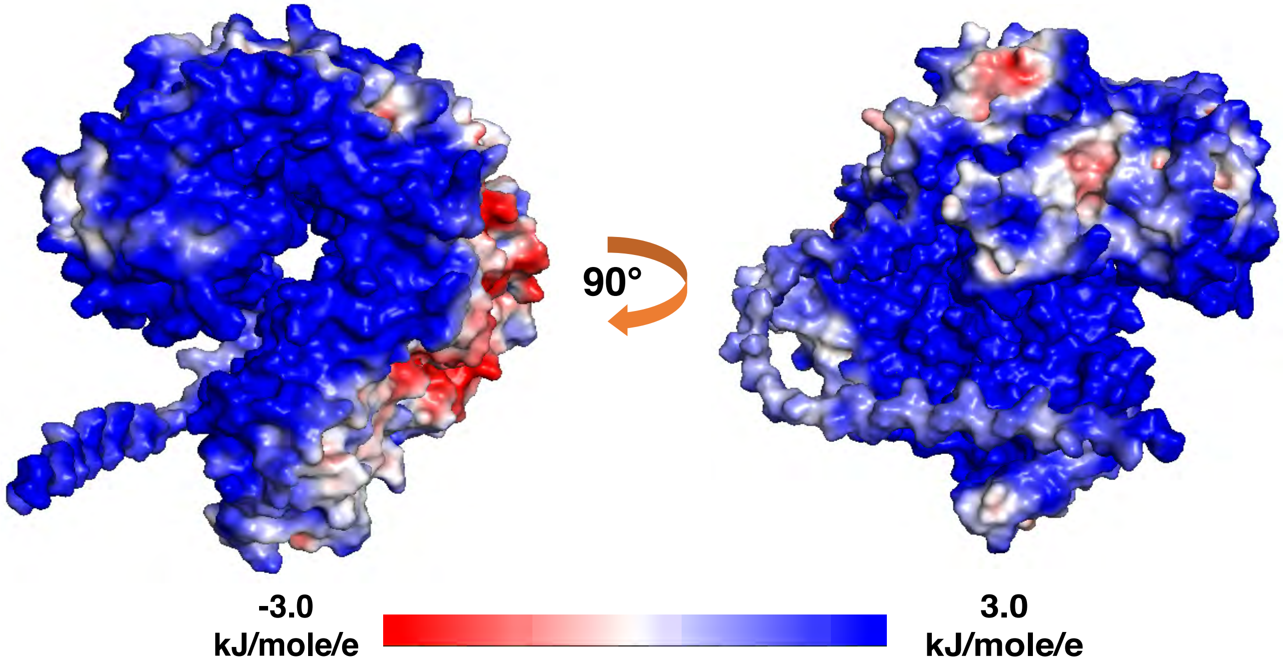


**B**

**3xFLAG::WAGO-1**

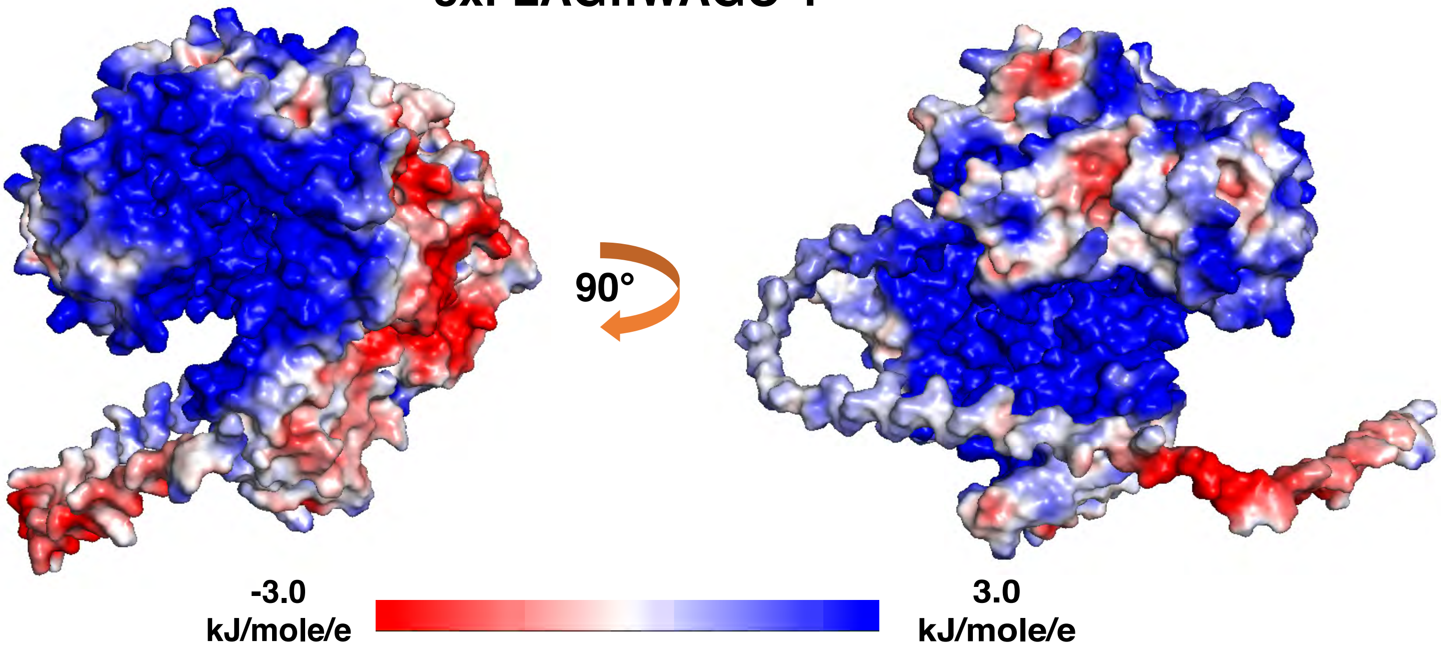

### Supplemental Figure S13

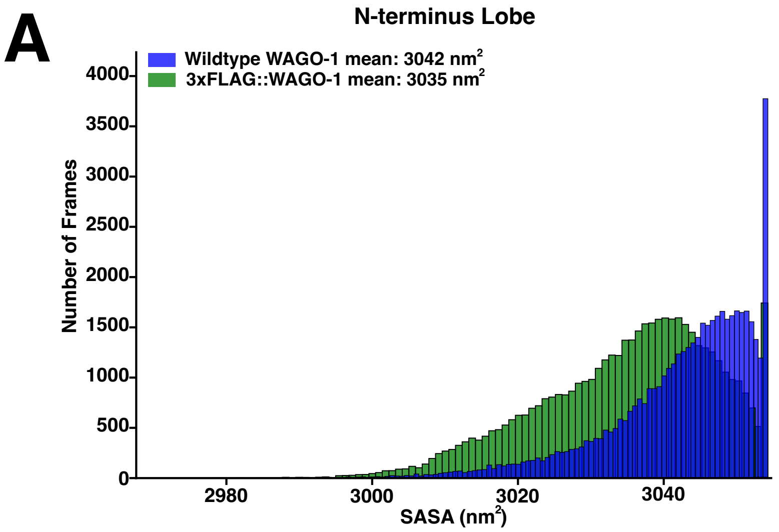

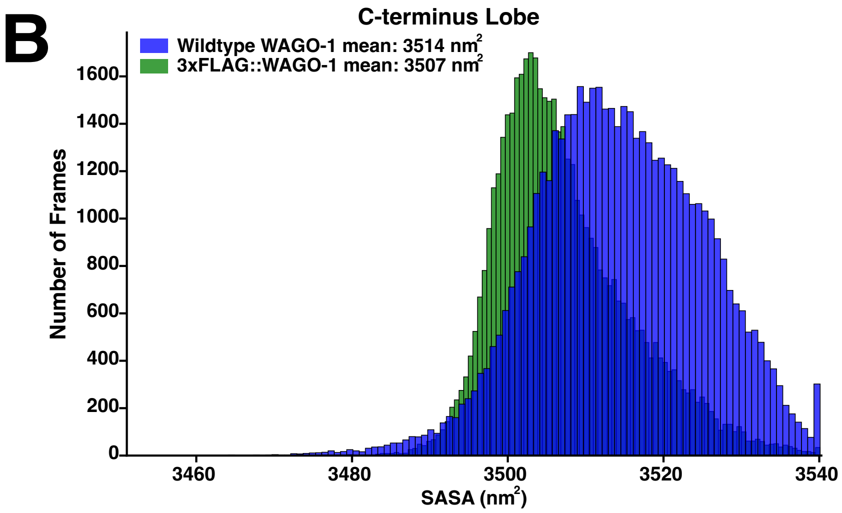

### Supplemental Figure S14

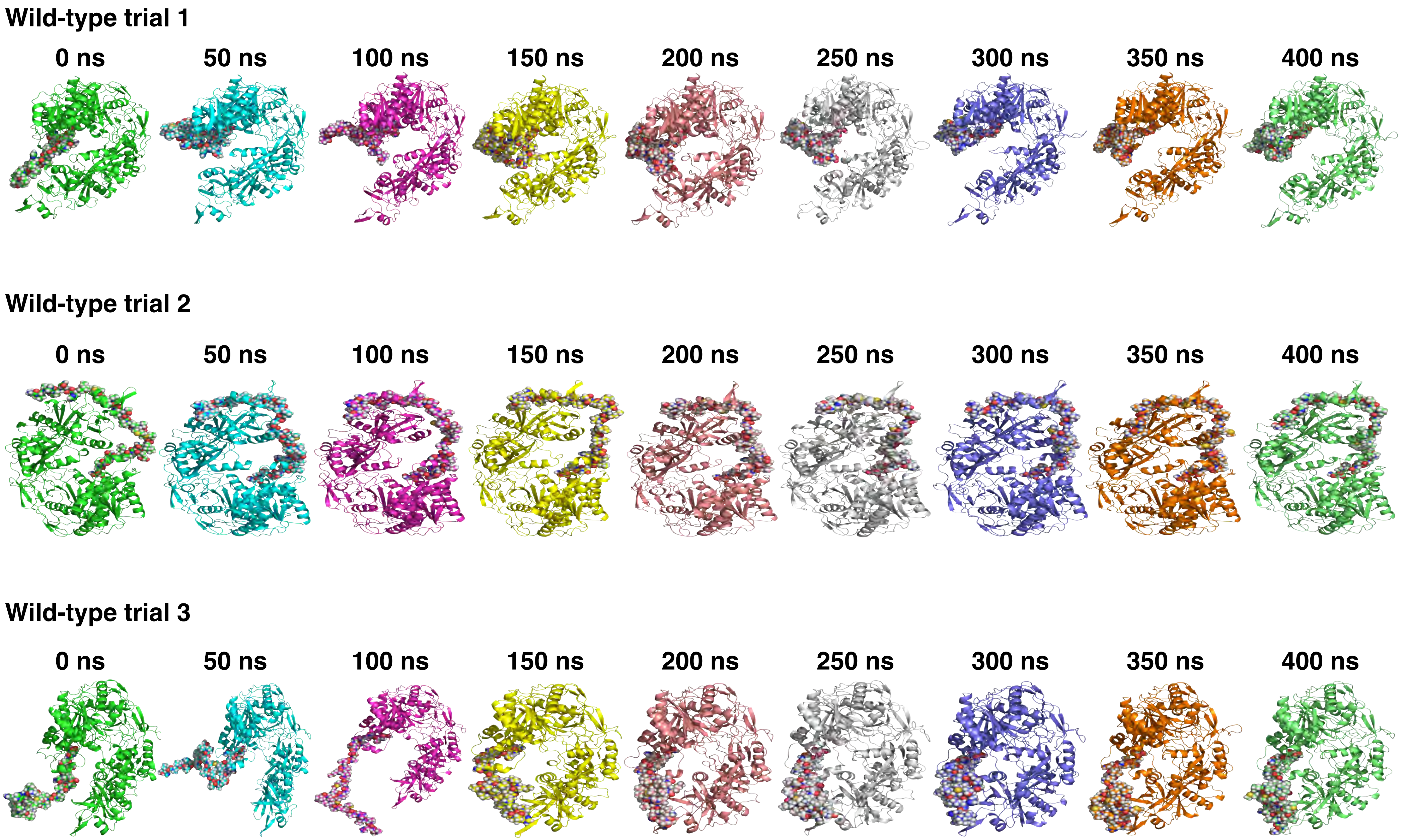

### Supplemental Figure S15

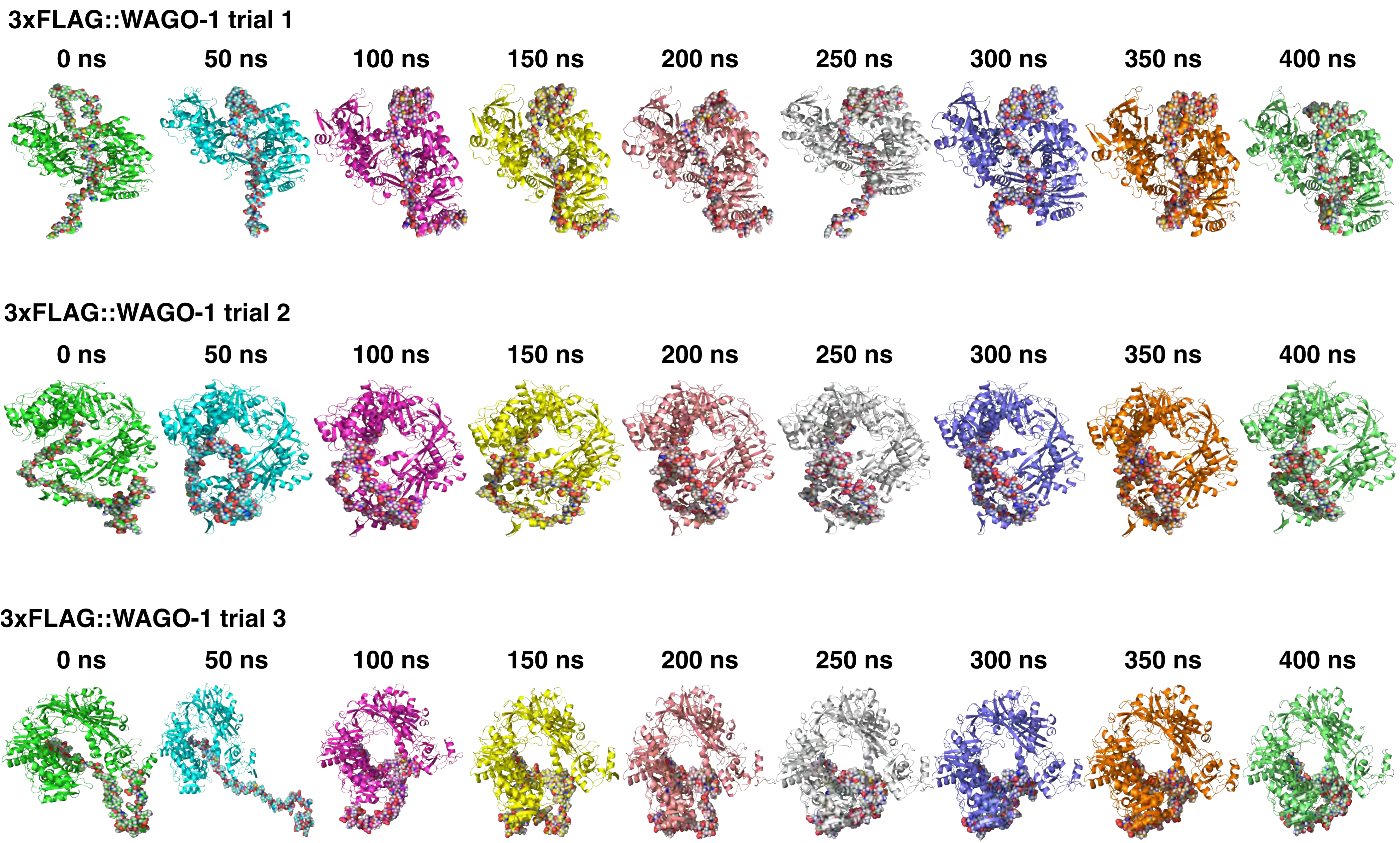

### Supplemental Figure S16

Figure S16

A

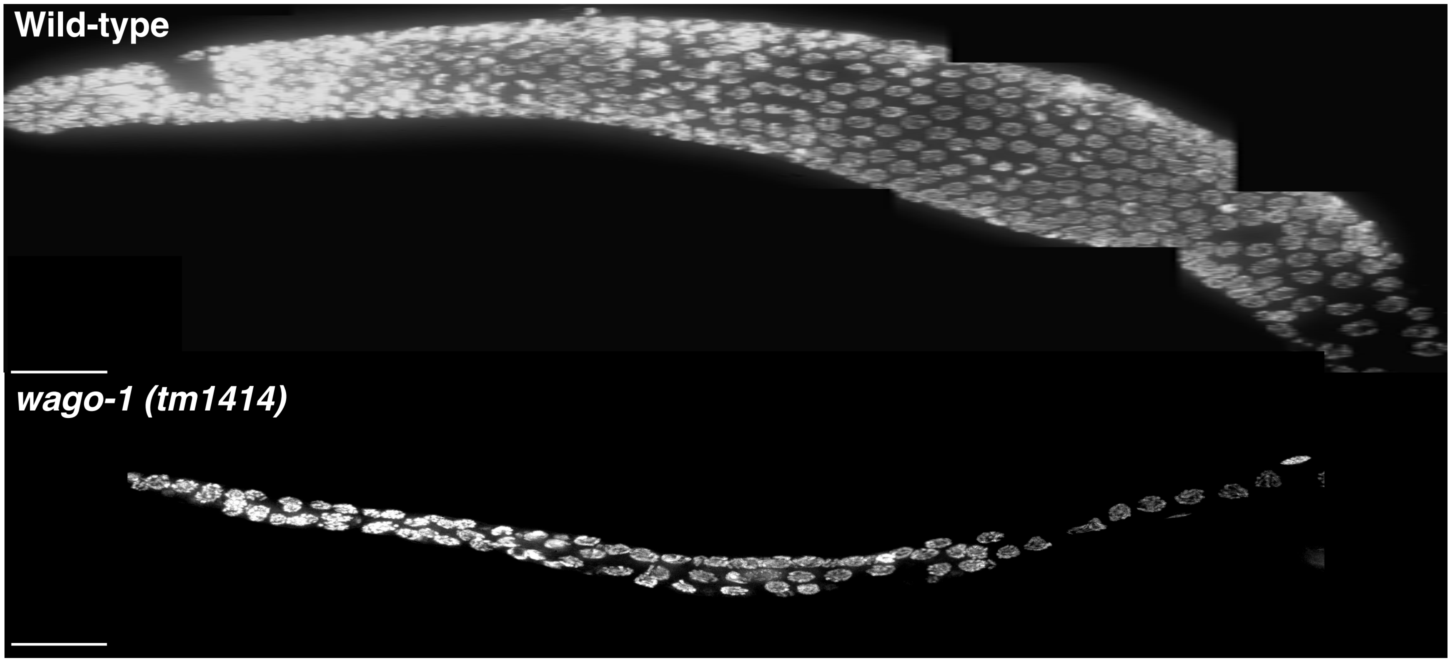


B

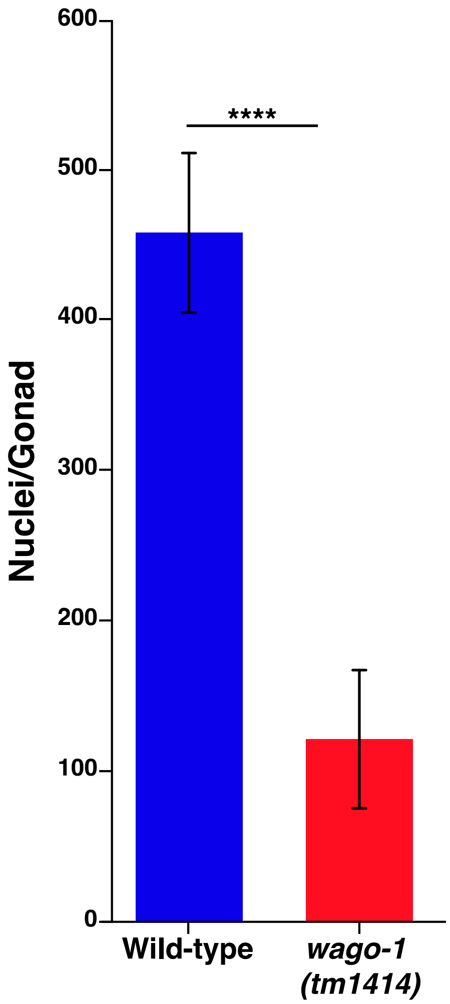

### Supplemental Figure S17

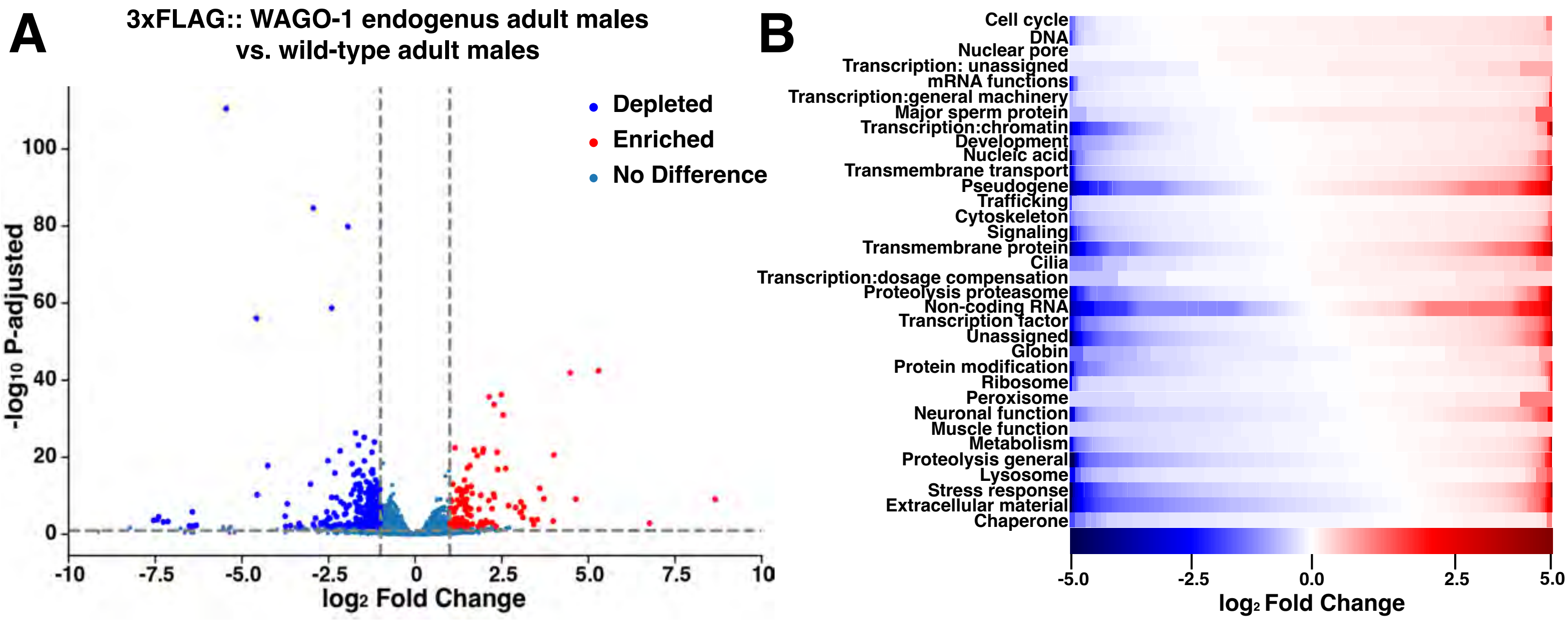

### Supplemental Figure S18

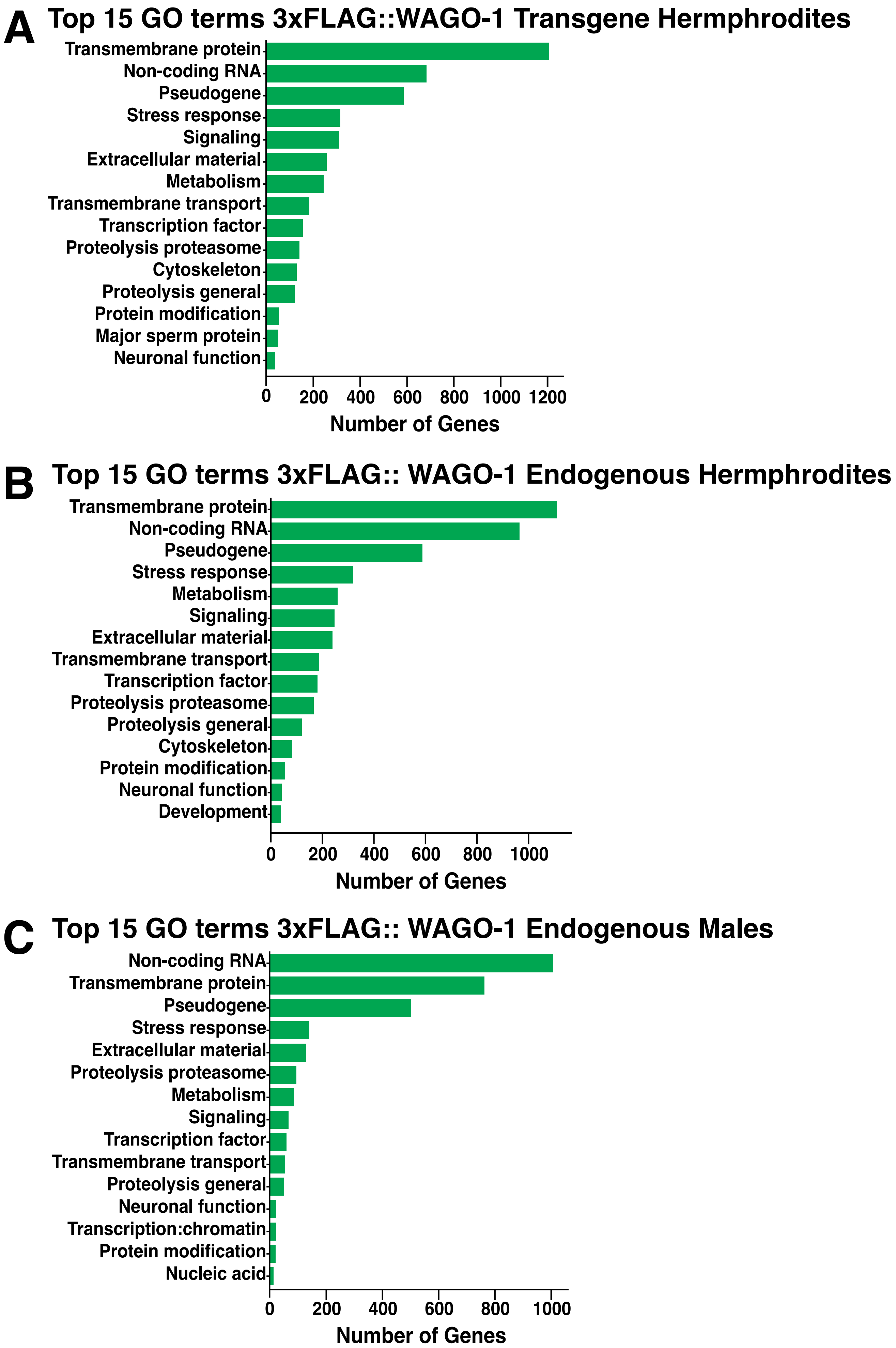

### Supplemental Figure S19

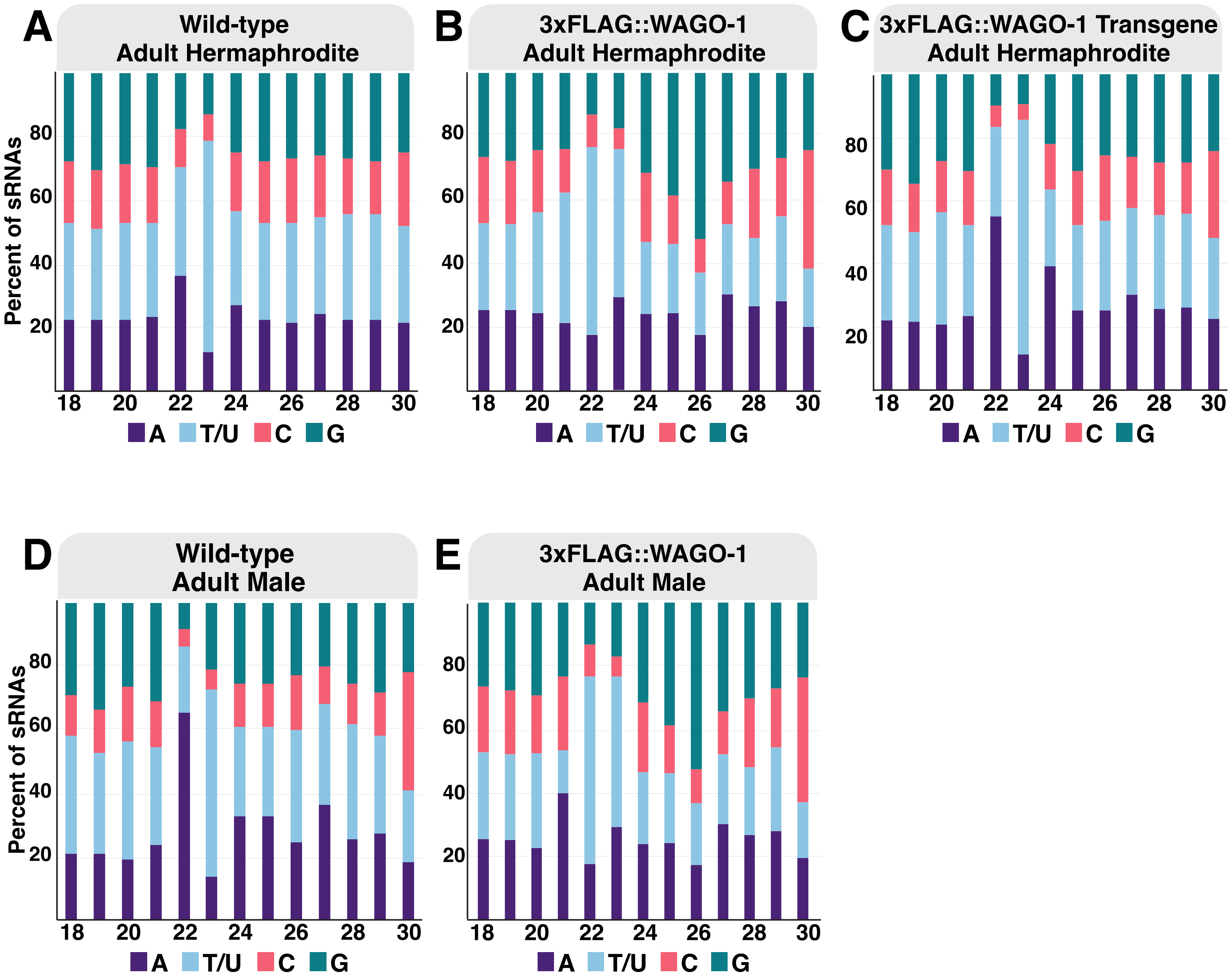

### Supplemental Figure S20

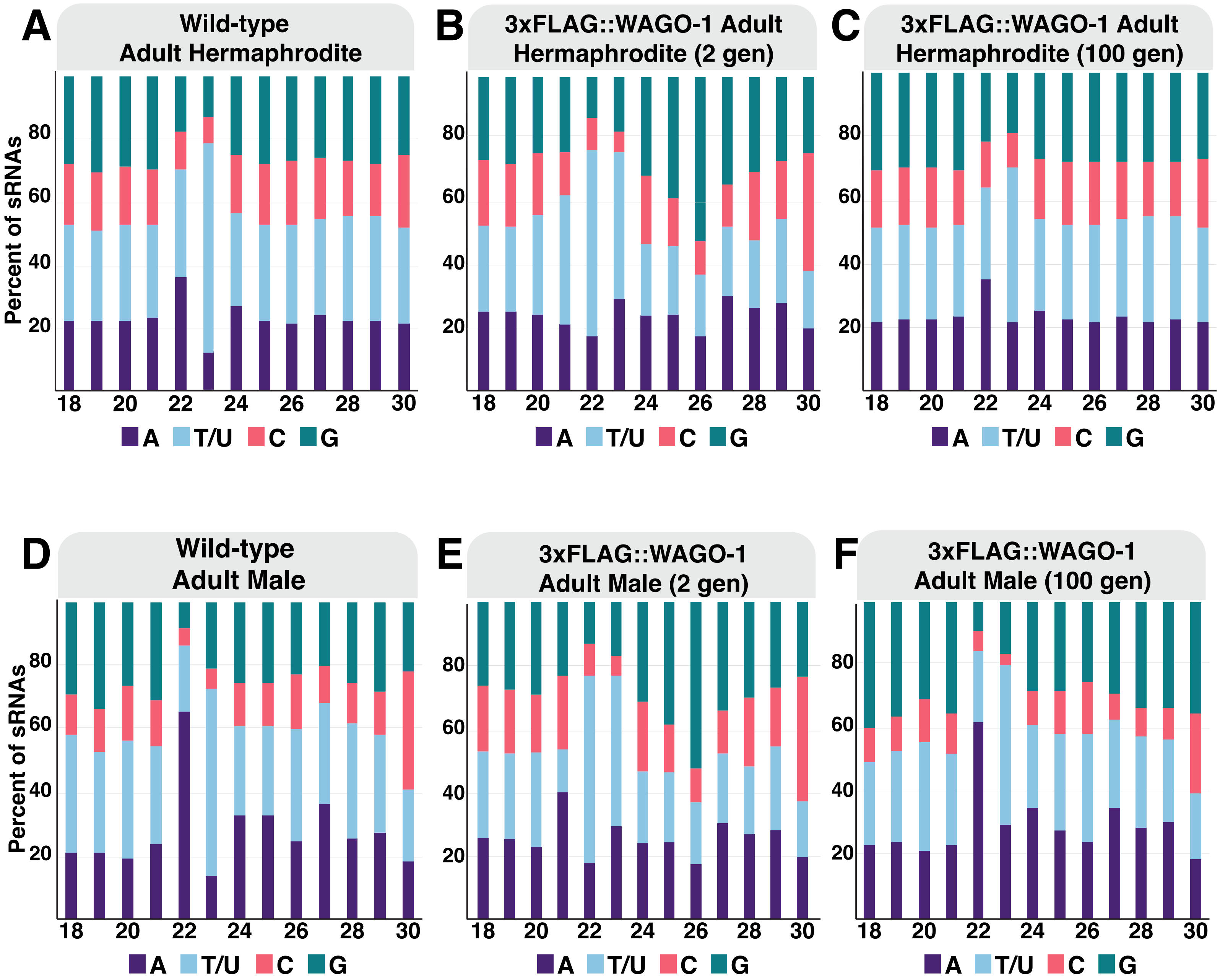

### Supplemental Figure S21

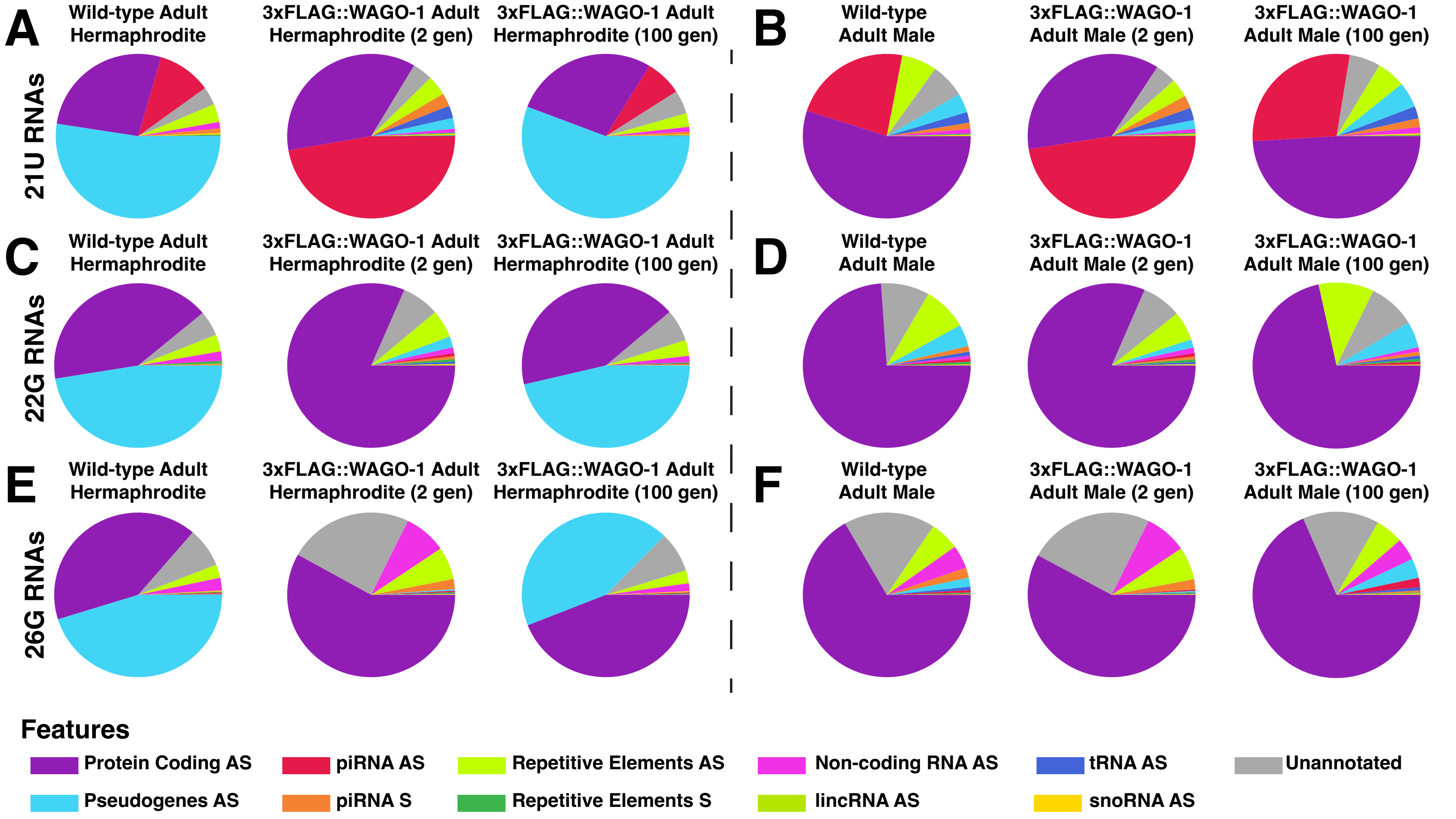

### Supplemental Figure S22

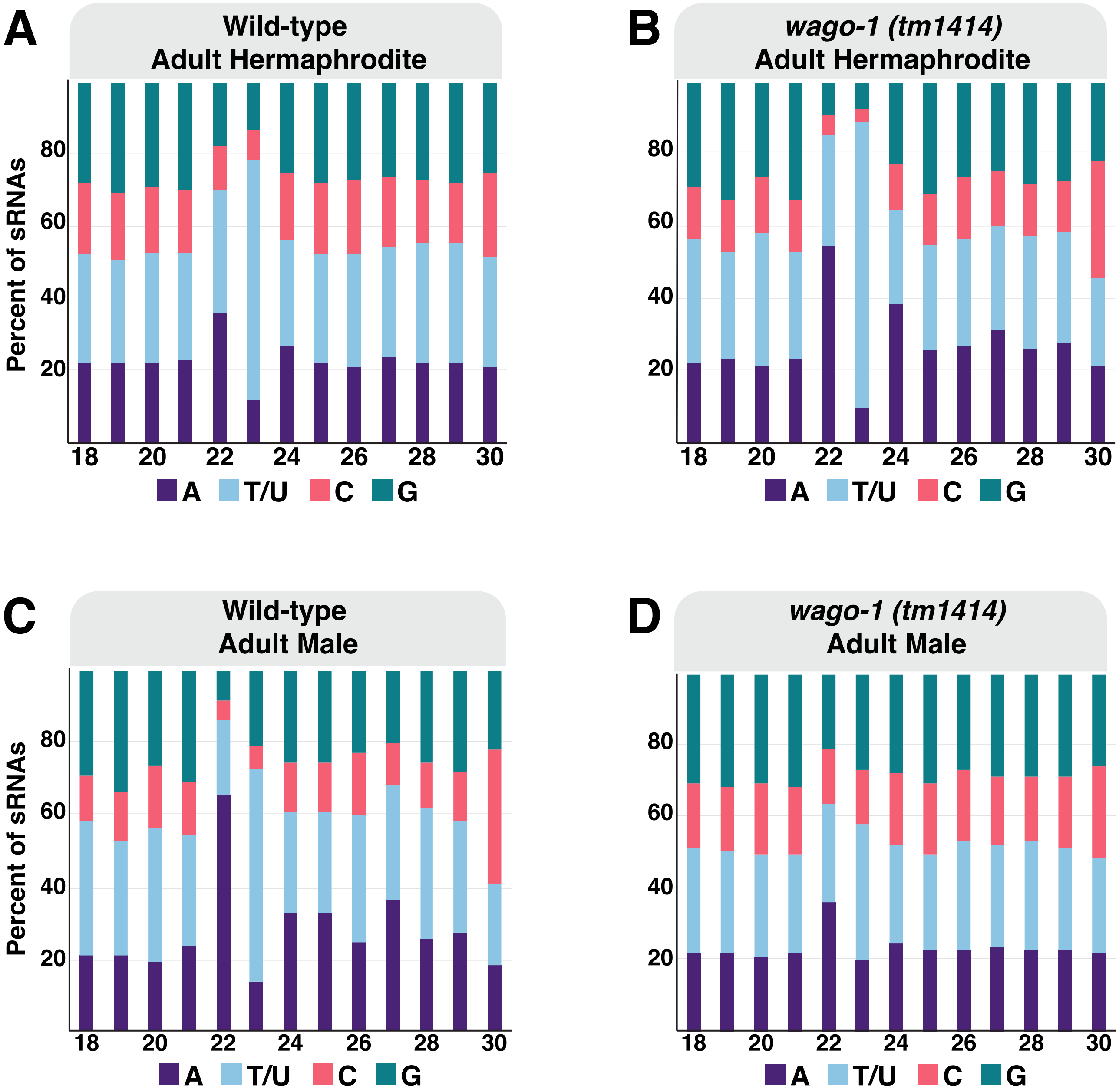
